# Protein Embeddings and Local Alignments

**DOI:** 10.1101/2025.07.23.666343

**Authors:** Julia Malec, G. Brian Golding, Lucian Ilie

## Abstract

**Background:** The advent of protein embeddings has revolutionized bioinformatics by providing contextual representations that capture functional and evolutionary patterns. They have become, alongside sequence alignments, the cornerstone of bioinformatics. While embeddings cannot replace alignments, they can greatly help improving their quality. Our goal is replacing the BLOSUM matrices, the decades-old standard scoring system for protein alignment, with an embedding-based scoring method.

**Results:** We introduce a new scoring function and algorithm for local alignment of protein sequences, we offer a new comprehensive framework for evaluating local alignments. The score between two residues is given by the cosine similarity of their Ankh-embedding vectors and the algorithm uses dynamic programming with affine penalty. For the evaluation, we built multiple datasets, using both natural and inserted sequences, from the Conserved Domain Database, BAL-iBASE, and GPCRdb, designed a new algorithm for local alignment extraction, localization and quality evaluation, and employed five distance metrics to evaluate the similarity with the true alignment. We performed nearly one and a half million tests to compare the new algorithm with the best BLOSUM matrices, specialized GPCRtm matrices, and top programs, such as PEbA, ProtT5-score, DEDAL, vcMSA and pLM-BLAST. Regarding the protein embedding models, Ankh not only surpasses the best combination of ProtT5 and ESM2, but appears to better understand the “language” of proteins, as it behaves much better on natural sequences compared to artificial ones obtained by inserting domains in random protein sequences. Also, while ProtT5 and ESM2 combine to produce better results, Ankh does not combine well with other embeddings.

**Conclusions:** The new Ankh-score-based program is vastly superior to the BLOSUM matrices and clearly superior to all existing methods. New light shed on the protein embeddings can guide future improvements. In order to facilitate the use of the new method and protocol, they are freely available as a web server at `e-score.csd.uwo.ca` and as source code at github.com/lucian-ilie/E-score.

## Background

The advent of protein embeddings such as ProtT5 (Elnaggar et al. 2021), ESM2 (Rives et al. 2021), or Ankh (Elnaggar et al. 2023), has revolutionized bioinformatics by providing contextual representations that capture functional and evolutionary patterns. They are helping to create state-of-the-art solutions for many problems in proteomics, such as structure prediction (Senior et al. 2020; Yang et al. 2020; Jumper et al. 2021), function prediction (Kulmanov and Hoehndorf 2020; Gligorijević et al. 2021; Lai and Xu 2022), interaction site prediction (Manfredi et al. 2023; Hosseini et al. 2024), etc. Embeddings have become, alongside sequence alignments, the cornerstone of bioinformatics.

Local alignments are crucial for comparing protein sequences as they discover regions of similarity within different sequences that may be functionally or evolutionarily related. This can help identify conserved domains, motifs, or functional regions, providing insights into protein function, structural characteristics, and evolutionary relationships. Detecting these conserved subsequences helps in annotating uncharacterized proteins, understanding protein families, and inferring potential functional roles of unknown proteins. Local alignment search is the most widely used procedure in bioinformatics and biological sciences. The well known BLAST program is one of the most cited contributions in the history of science; the two BLAST papers (Altschul et al. 1990, 1997) combined exceed 205,000 citations on Google Scholar (as of Apr. 2025).

In spite of the thorough investigation of this problem over many years, alignment of distantly related protein sequences remains challenging. Accurate local alignment is particularly important when searching for homologous sequences that have diverged significantly, making detection of relevant similarities difficult. More precise alignment algorithms increase the chance of identifying subtle similarities, allowing researchers to uncover distant evolutionary relationships and functional conservation across species.

The key feature in all local alignments algorithms is the scoring method that evaluates the similarity between amino acid residues. BLOSUM matrices (Henikoff and Henikoff 1992) provided the standard scoring procedure for more than three decades, with another popular method provided by PAM matrices (Dayhoff et al. 1978). Different BLOSUM matrices, such as BLOSUM45, BLOSUM50, BLOSUM62, BLOSUM80, and BLOSUM90, are optimized for detecting homology at various sequence similarity levels, with lower numbers matrices being designed to discover more distantly related sequences.

Recent developments in deep learning provide the ideas for improvement. Techniques from Natural Language Processing involving self-supervised learning on very large unlabelled data helped create word embeddings, that is, numerical vectors associated with words, which can preserve their meaning (Mikolov et al. 2013; Pennington et al. 2014; Peters et al. 2018; Devlin et al. 2018; Liu et al. 2019; Yang et al. 2019; Raffel et al. 2020). These ideas were applied to protein sequences, with entire proteins seen as sentences and residues as words, by training on many protein sequences from UniProt (UniProt Consortium 2018). Top protein embeddings include: ProtVec (Asgari and Mofrad 2015), SeqVec (Heinzinger et al. 2019), PRoBERTa (Nambiar et al. 2020), MSA-transformer (Rao et al. 2021), ESM-1b and ESM2 (Rives et al. 2021), ProtAlbert, ProtBert, ProtT5, ProtElectra, ProtXLNet (Elnaggar et al. 2021), Ankh (Elnaggar et al. 2023).

Alignments will continue to provide the most basic sequence-level information, critical for many applications, while embeddings provide the most basic functional or contextual information, ideal for predictive analyses. While embeddings cannot replace alignments they can help improving their quality. The cosine similarity of embedding vectors associated with amino acid residues has been used recently as a successful scoring scheme by several independent studies, for similarity search with pLM-BLAST (Kaminski et al. 2023), multiple sequence alignment with vcMSA (McWhite et al. 2023), pairwise alignment with PEbA (Iovino and Ye 2024) and E-score (Ashrafzadeh et al. 2024).

Our goal here is two fold. First, we give the best local alignment algorithm for protein sequences, together with a new comprehensive framework for evaluating local alignments. The algorithm uses dynamic programming (Smith and Waterman 1981; Gotoh 1982) with the score provided by the cosine similarity of embedding vectors associated with amino acid residues by Ankh. To assess its performance, we introduce a thorough protocol for the evaluation of local alignment computation, including building multiple datasets, suitable for local alignment computation, from the Conserved Domain Database (CDD) (Marchler-Bauer et al. 2015), BALiBASE database (Thompson et al. 2005), and GPCRdb (Pándy-Szekeres et al. 2022), designing an algorithm for local alignment extraction, localization and quality evaluation, and using five distance metrics for evaluating the similarity with the true alignment. Second, our investigations shed new, important light on the protein embeddings themselves. Ankh not only surpasses the best combination of ProtT5 and ESM2, but appears to better understand the “language” of proteins, as it behaves much better on natural sequences compared to artificial ones obtained by inserting domains in random protein sequences. Also, while ProtT5 and ESM2 combine to produce better results, Ankh does not combine well with other embeddings.

We hope that using the new method for computing local alignments will have a significant impact in the fields of bioinformatics and evolutionary biology, improving on how protein sequences are aligned, compared, and interpreted, enhancing the precision of identifying homologous sequences, especially for distantly related proteins, where BLOSUM matrices often struggle. It should improve the detection of functional domains, rare motifs, and evolutionary relationships, which are critical for annotating proteins in newly sequenced genomes. This could have direct implications in various fields, such as drug discovery, protein engineering, and functional genomics. We hope as well that our findings regarding protein embedding methods indicate the possibility of future improvement and can help guide such developments.

## Methods

### Local alignment evaluation setup

A successful method for local alignment computation has to correctly address two aspects:

- *alignment localization*: identification of the subsequences participating in the local alignment, and
- *alignment quality* : alignment of the subsequences involved, as measured by the distance to the reference alignment.

This basic setup is used below in designing the datasets and algorithms used for proper evaluation.

### Datasets

A domain consists of a set of similar sequences for which a reference multiple sequence alignment (MSA) is known. A test consists of two protein sequences containing two similar subsequences from the same domain. Ideally, these two subsequences are the only significant similarities between the two protein sequences. We have imposed two additional conditions on the data. First, each domain occurrence inside a protein sequence has to be away from the sequence ends (usually we require at least 30 residues away from either end), to be considered a proper local alignment; otherwise the alignment is essentially global. Second, in order to have statistically significant p-values, we used, whenever possible, domains with at least 15 sequences, such that the number of tests is over 100. (We made exceptions for the BALiBASE datasets RV11 and RV12, otherwise almost all domains would be eliminated.) We constructed several datasets with these properties. Natural datasets have the domain sequences naturally occurring inside protein sequences. Insert datasets are built by inserting domain sequences into random protein sequences. All datasets are summarized in Table 1. Note the wide amino acid identity range for the entire collection of data.

**Table 1.** Dataset summary. For each dataset, we give the number of domains, sequences, tests performed, average protein and domain length and amino-acid identity range. ^*^Total tests are calculated using the number of times each dataset was used multiplied by the number of distances calculated.

| Database source | Domains | Sequences | Tests | Protein len. | Domain len. | Identity range |
| --- | --- | --- | --- | --- | --- | --- |
| Natural data |  |  |  |  |  |  |
| CDD | 8 | 354 | 9,196 | 427.2 | 185.4 | 8.00% - 24.55% |
| BALiBASE RV11 | 9 | 62 | 224 | 416.9 | 265.0 | 11.44% - 18.32% |
| BALiBASE RV12 | 5 | 51 | 258 | 657.1 | 396.0 | 24.50% - 35.26% |
| BALiBASE RV30 | 10 | 606 | 8,152 | 453.2 | 267.3 | 23.34% - 42.94% |
| Insert data |  |  |  |  |  |  |
| CDD | 8 | 354 | 9,196 | 427.2 | 185.4 | 8.00% - 24.55% |
| BALiBASE RV911 | 29 | 423 | 29,000 | 780.4 | 642.2 | 9.27% - 24.26% |
| BALiBASE RV912 | 28 | 228 | 28,000 | 613.4 | 476.2 | 26.09% - 35.57% |
| BALiBASE RV913 | 27 | 263 | 27,000 | 607.5 | 470.5 | 48.34% - 65.94% |
| GPCRdb | 8 | 290 | 8,000 | 400.4 | 259.9 | 16.64% - 42.77% |
| Total | 132 | 2,631 | *1,457,350 | 531.5 | 349.7 | 8.00% - 65.94% |

We have produced, according to the above requirements, four natural datasets, from the Conserved Domain Database (Marchler-Bauer et al. 2015) and the BALiBASE (Thompson et al. 2005) datasets RV11 -highly divergent, RV12 - moderately divergent, and RV30 - divergent families. Their details are given in Table 2. Then, for greater variety, we have produced several insert datasets. Three are from the BAL-iBASE linear motifs datasets RV911 - highly divergent, RV912 - moderately divergent, and RV913 - closely related sequences, one is from the GPCRdb database (Pándy-Szekeres et al. 2022), for comparison against specialized scoring matrices, and we use the CDD dataset also in insert mode to compare with the natural one. The details of all datasets are given in Supplementary Tables 1-12.

**Table 2.** Natural dataset details. From left to right, domain ID, number of sequences, average length of protein and domain sequences, average sequence length before and after domain occurrences, and amino acid percentage identity.

| Domain | Sequences | Protein length | Domain length | Before | After | Identity |
| --- | --- | --- | --- | --- | --- | --- |
| CDD |  |  |  |  |  |  |
| cd00012 | 60 | 398.07 | 316.18 | 33.37 | 48.52 | 8.00 |
| cd00083 | 47 | 537.87 | 47.36 | 287.45 | 203.06 | 24.55 |
| cd00173 | 26 | 701.23 | 82.00 | 259.92 | 359.31 | 15.76 |
| cd00637 | 78 | 388.44 | 283.62 | 35.05 | 69.77 | 16.86 |
| cd01040 | 62 | 211.84 | 133.60 | 49.84 | 28.40 | 15.68 |
| cd01068 | 19 | 394.95 | 145.47 | 16.32 | 233.16 | 16.30 |
| cd10140 | 36 | 386.08 | 155.81 | 173.31 | 56.97 | 9.56 |
| cd21116 | 26 | 399.31 | 318.77 | 25.23 | 55.31 | 12.22 |
| BALiBASE RV11 |  |  |  |  |  |  |
| BBS11009 | 4 | 216.00 | 91.00 | 59.75 | 65.25 | 18.28 |
| BBS11010 | 4 | 491.50 | 178.25 | 37.75 | 275.50 | 15.16 |
| BBS11016 | 8 | 470.50 | 324.13 | 57.00 | 89.38 | 15.74 |
| BBS11018 | 14 | 547.93 | 432.79 | 73.71 | 41.43 | 13.51 |
| BBS11024 | 4 | 407.00 | 302.75 | 36.50 | 67.75 | 11.44 |
| BBS11026 | 7 | 240.00 | 83.29 | 34.43 | 122.29 | 12.58 |
| BBS11034 | 8 | 491.63 | 336.75 | 56.88 | 98.00 | 15.33 |
| BBS11037 | 5 | 523.40 | 367.80 | 83.60 | 72.00 | 12.28 |
| BBS11038 | 8 | 364.25 | 268.13 | 49.63 | 46.50 | 18.32 |
| BALiBASE RV12 |  |  |  |  |  |  |
| BBS12008 | 13 | 436.77 | 367.23 | 36.15 | 33.38 | 27.63 |
| BBS12011 | 12 | 869.08 | 589.58 | 46.75 | 232.75 | 25.04 |
| BBS12030 | 6 | 790.33 | 236.17 | 56.83 | 497.33 | 35.26 |
| BBS12033 | 7 | 454.71 | 306.14 | 73.57 | 75.00 | 24.50 |
| BBS12037 | 13 | 734.85 | 480.92 | 182.69 | 71.23 | 25.01 |
| BALiBASE RV30 |  |  |  |  |  |  |
| BBS30001 | 116 | 468.01 | 178.41 | 143.79 | 145.81 | 31.51 |
| BBS30004 | 50 | 497.26 | 391.42 | 34.14 | 71.70 | 30.79 |
| BBS30005 | 113 | 479.78 | 235.58 | 214.04 | 30.16 | 29.38 |
| BBS30006 | 18 | 303.39 | 94.67 | 166.33 | 42.39 | 23.34 |
| BBS30008 | 36 | 526.81 | 409.36 | 70.64 | 46.81 | 28.77 |
| BBS30009 | 38 | 171.45 | 67.50 | 30.05 | 73.89 | 28.86 |
| BBS30010 | 50 | 655.54 | 554.74 | 68.78 | 32.02 | 42.94 |
| BBS30013 | 86 | 656.26 | 407.91 | 167.07 | 81.28 | 26.39 |
| BBS30016 | 37 | 291.30 | 61.16 | 148.27 | 81.86 | 34.73 |
| BBS30030 | 62 | 482.53 | 271.77 | 126.84 | 83.92 | 27.98 |

We used multiple testing datasets to evaluate our method under a variety of situations. These included datasets from NCBI’s Conserved Domain Database as the alignment of these sequences are curated by hand according to experimental data of the function of the proteins. The test datasets also included datasets from BAliBASE, as again it is a curated dataset designed to be a dataset of alignment standards. From this we choose subsets that covered a range of divergences.

### Alignment computation

Given a protein embedding method, *E*, and two protein sequences, we use the classic dynamic programming Smith and Waterman (1981) algorithm to compute the local alignment, using the affine gap penalty variant of Gotoh (1982). The score between two residues *a*_1_ and *a*_2_ is the *E*-score we introduced in Ashrafzadeh et al. (2024), given by the cosine similarity between the vectors computed by *E* for the two residues:

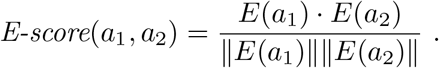

For a fixed method *E*, such an Ankh, we obtain Ankh-score. The penalties are empirically set at −1 for gap opening and −0.1 for gap extension. Note that the scoring function of Iovino and Ye (2024) is essentially the same; they multiply the cosine by 10, but keeping the original cosine and dividing their penalties by 10 produces the same results. (They use slightly different gap penalties, −11 and −1, which correspond to our −1.1 and −.1.)

In order to correctly evaluate the quality of a given local alignment, we need to calculate its distance to the reference alignment using a proper distance metric. Distance metrics require the alignments to be between the *same* sequences. Therefore, we need an algorithm that computes, from any alignment produced by some method, a local alignment between the same sequences as the reference. The idea is to eliminate sequences outside reference and also pad the computed alignment, if necessary, on either side with the worst alignment of the missing subsequences, denoted Worst(·,·), which is the alignment of length the sum of the lengths of the two sequences, with each sequence contiguous and aligned with gaps only, e.g.,

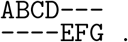

The local alignment computation algorithm is given below.

LocalAlignment 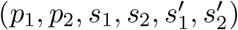

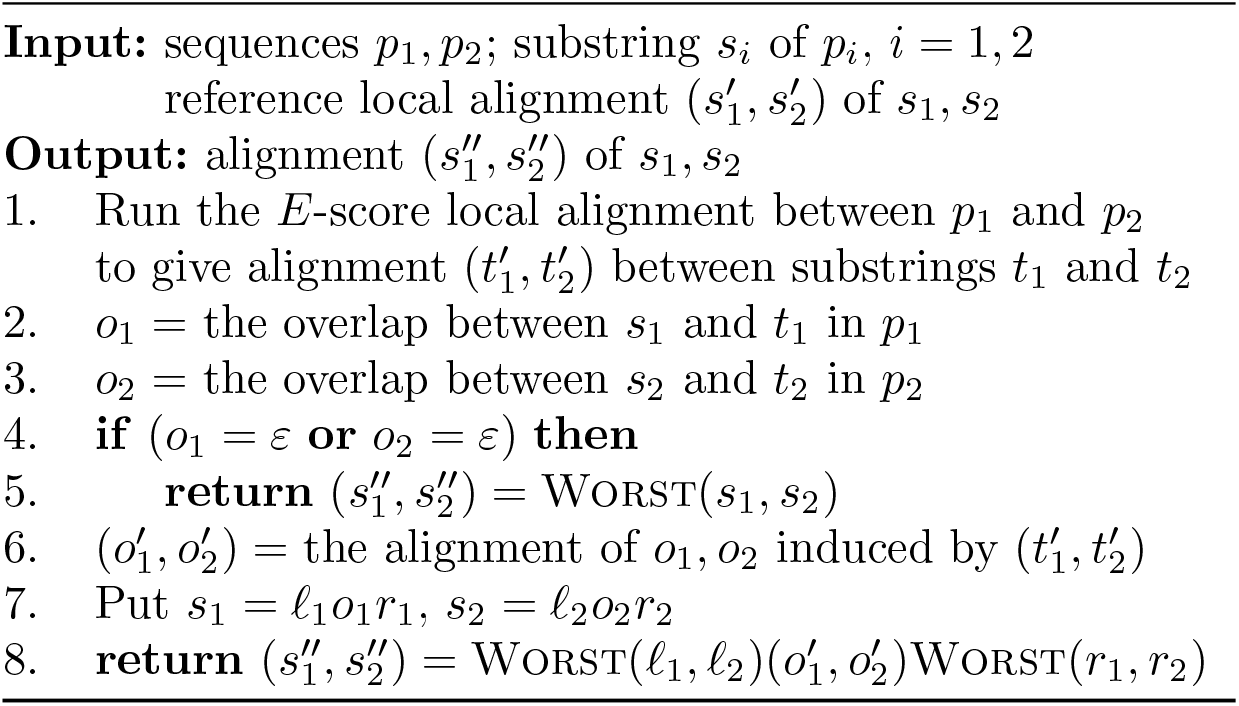

For clarity, we provide below an example of running the algorithm. Consider the following protein sequences *p*_1_ and *p*_2_ containing the domain sequences, *s*_1_ and *s*_2_, shown in red, having the reference alignment 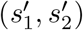:

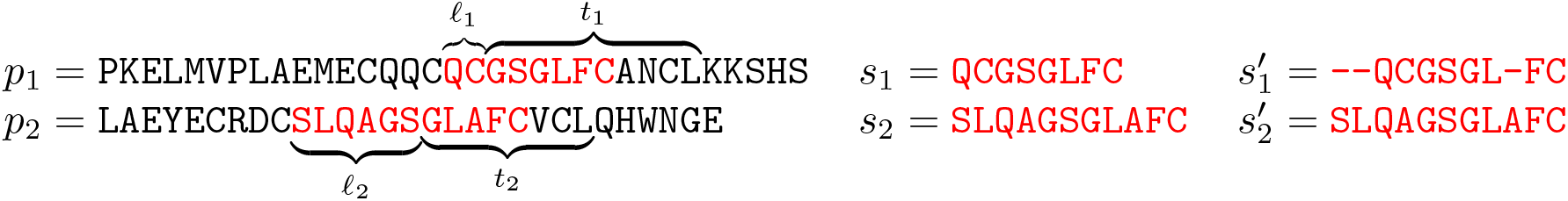

Assume that the *E*-score local alignment algorithm produces the alignment 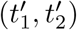 between the substrings *t*_1_ and *t*_2_, indicated by curly braces above and below the sequences; the red parts indicate the overlaps, *o*_1_ and *o*_2_, with the reference domains:

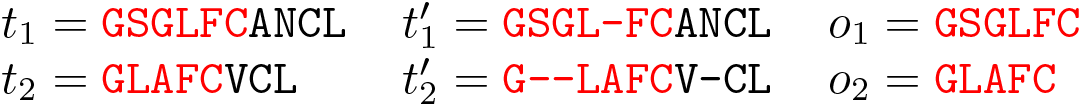

The alignment, 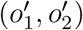, of the overlaps induced by 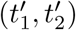, the extra sequences to the left and to the right, and the final alignment, 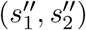, of the domain sequences, *s*_1_ and *s*_2_, induced by the *E*-score alignment are given below:

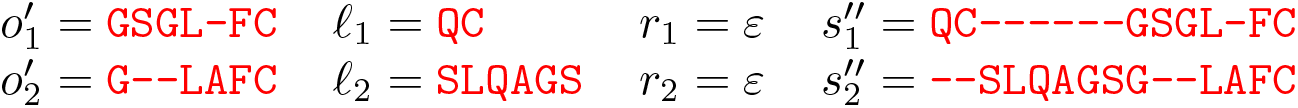

The evaluation of the quality of the alignment produced by this process consists of measuring the distance between the alignment 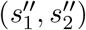 and the reference alignment 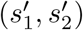. Note that both of these alignments are between the same reference domain sequences *s*_1_ and *s*_2_.

### Comparison procedure

Assume a fixed protein embedding method. Given a domain and two methods for computing local alignment, we perform a number of specified tests, each consisting of a pair of sequences. The local alignment is computed using both methods and then the results are compared according to their distances to the reference alignment, as induced by the domain’s MSA for the given sequences.

Potentially the alignments could differ in multiple ways and hence we used five distance metrics to evaluated the difference in the alignments. Three of those are from Blackburne and Whelan (2012) – d_ssp_, d_seq_, d_pos_ – and two newly introduced in our global alignment study (Ashrafzadeh et al. 2024) – d_d_, d_cc_. The former three are true metrics inspired from the sum-of-pairs score, differing from each other in the way they consider gaps: ignore, label by sequence, and label by sequence and position, resp. This makes d_pos_ the most relevant of the three. The other two are very different; d_d_, relative displacement distance, evaluates all position differences of all residues; d_cc_, closest context distance, takes the middle ground, by measuring the distance to the closest position with the same context.

After evaluating all tests as explained above, we report, for each domain and each distance, two values for each method: (i) the average distance between local alignments computed by the method to the corresponding reference alignment and (ii) the number of tests won by the method. Except for rare cases, a method prevails with respect to both the average distance and the number of tests won. Methods are always compared in pairs, in order to enable the use of the Wilcoxon signed-rank test; we consider p-values smaller than 0.01 significant. Values higher than this threshold are shown in red.

### Score modifications

We have explained above how alignment quality is evaluated. Concerning the alignment localization, the above evaluation penalizes the situation when a computed alignment misses part of the reference alignment by replacing the missing part with worst alignment of the subsequences missed. If, however, the computed alignment extends outside the reference alignment, the length of the overhangs cannot be measured properly because there is no golden standard for that. Instead, we offer the user the possibility of adjusting the size of the overhangs. If we make the scores more negative by shifting them down by a fixed value, then the size of the overhangs decreases. Fig. 1 shows how various shifts affect the amount of alignment outside the reference compared to what BLOSUM matrices produce. The quality of the alignments is little affected by using the shifts, therefore this is a good way for the user to adjust the alignment length, if desired.

**Fig. 1.**
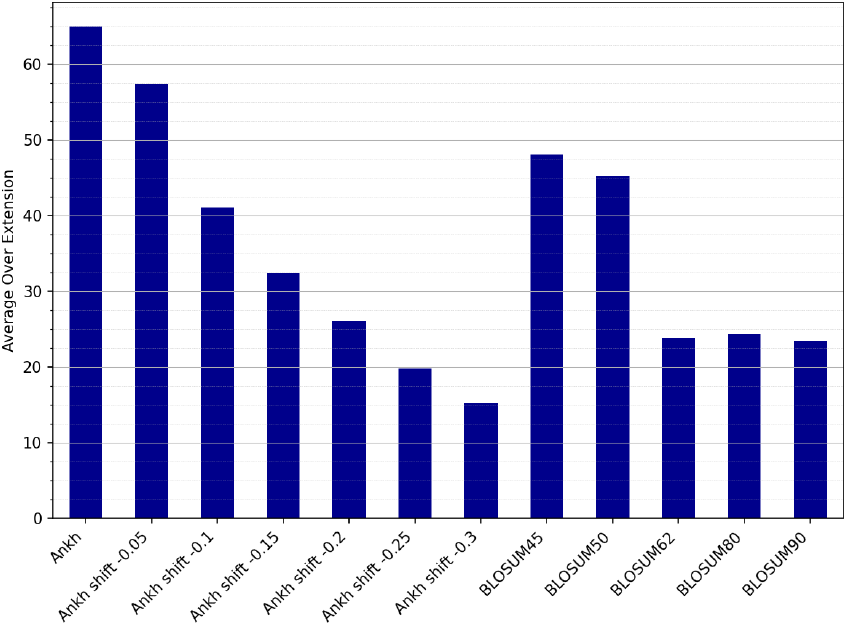
Alignment overhangs. Average lengths of the alignment overhangs for the CDD natural dataset are given for all BLOSUM matrices and several shift values of Ankh-score.

## Results

### Model selection

We have first tested several embedding methods to see which one works best for our task. Supplementary Table 13 gives the comparison of seven top methods on the CDD natural dataset. For all five distances, Ankh-score and ProtT5-score are the best, with Ankh-score usually first, followed at some distance by ESM2. Therefore, we are going to use Ankh-score as our scoring function. Note that the statistical significance of the comparison between Ankh-score and ProtT5-score is assessed thoroughly later in this section, when we compare the two on eight datasets.

### Competing methods

The candidates for comparison include the standard for scoring protein alignments, the BLOSUM matrices, which we feel should be replaced by the new method, specialized matrices such as GPCRtm, and existing top methods, PEbA (Iovino and Ye 2024), ProtT5-score (Ashrafzadeh et al. 2024), DEDAL (Llinares-López et al. 2023), vcMSA (McWhite et al. 2023), and pLM-BLAST (Kaminski et al. 2023).

The first evaluation we did was to assess how much of the alignments are missed, on the average, by each method. We used the CDD natural dataset for this purpose and the results are plotted in Fig. 2; all values are given in Supplementary Table 14. We do not compare with vcMSA because it is a heuristic version of PEbA or E-score, therefore generally weaker, as it is intended for multiple sequence alignments, therefore unable to use dynamic programming. Also, as already mentioned PEbA uses esentially the same score as ProtT5-score, therefore, comparing against ProtT5-score will cover both. We are not comparing with pLM-BLAST due to its poor performance covering alignments, as explained above.

**Fig. 2.**
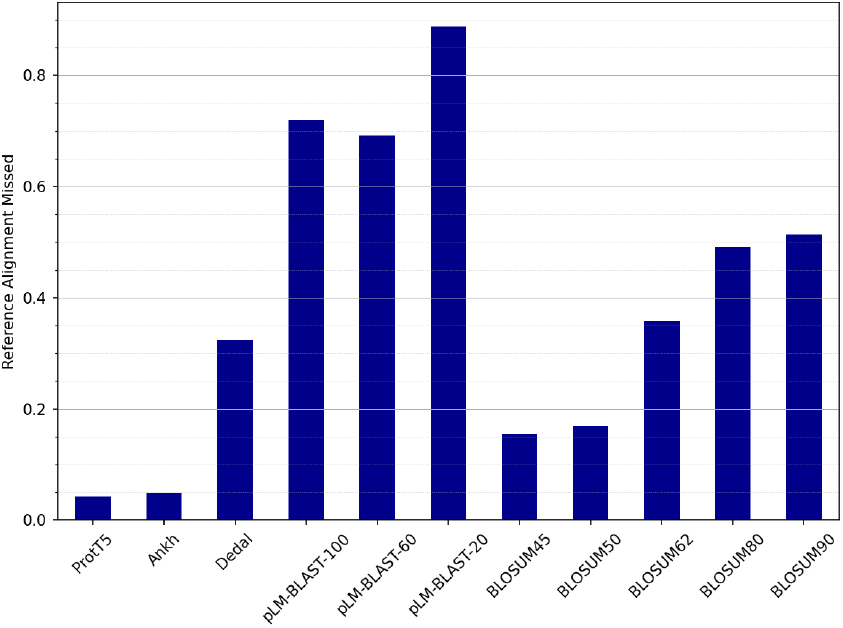
Missed alignments. Average percentage of the alignments missed by each scoring method on the CDD natural dataset. For pLM-BLAST we tested several values (100, 60, 20) for the minimum alignment parameter, attempting to reduce the amount of alignments missed.

### Comparison against BLOSUM

First, we give an idea of what the scores look like in Fig. 3, where we plotted the actual scores for both methods, Ankh-score and BLOSUM, for two typical examples, one natural example and one insert. It is quite striking how clearly visible the alignments are in Ankh-score plots, whereas in the BLOSUM plots, for the insert case the alignment is barely visible, while in the natural case it cannot be detected visually.

**Fig. 3.**
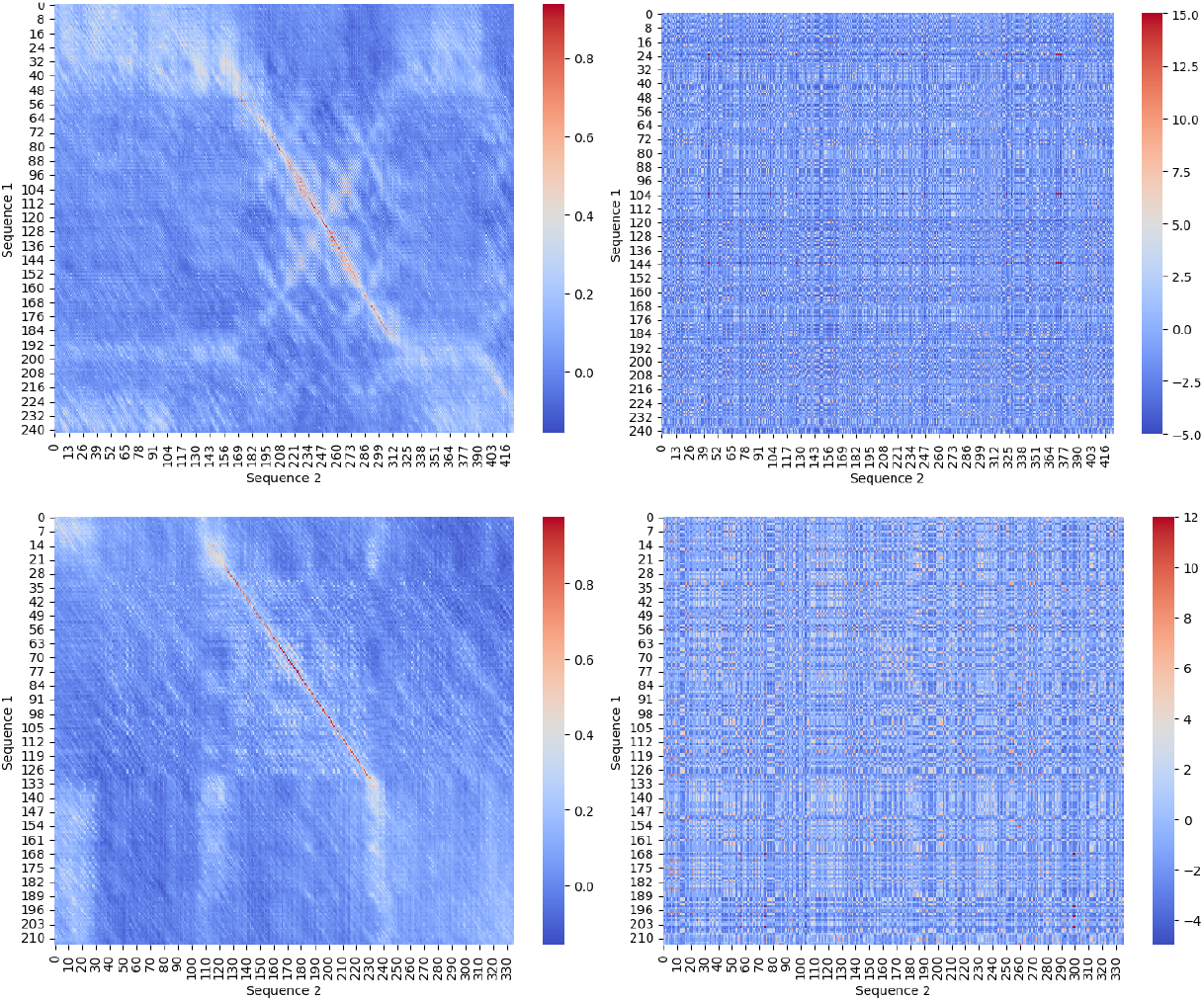
Score examples. Ankh-scores (left column) and BLOSUM scores (right column) are given for two examples, a natural one (top row) and an insert one (bottom row).

As we aim to replace BLOSUM matrices by Ankh-score, we perform a thorough comparison against the five common BLOSUM matrices – 45, 50, 62, 80 and 90 – on all datasets from Table 1 (except GPCR). We give as an example the comparison on the CDD natural datasets for the first three distances in Table 3. Full results for all datasets and all distances are given in Supplementary Tables 15-30, including also the BLOSUM matrices that produced the best results.

**Table 3.** Ankh-score vs best BLOSUM on CDD natural dataset, for the first three distances.

| Domain | d<sub>cc</sub> |  |  | d<sub>d</sub> |  |  | d<sub>pos</sub> |  |  |
| --- | --- | --- | --- | --- | --- | --- | --- | --- | --- |
| Average distances |  |  |  |  |  |  |  |  |  |
|  | Ankh | BLOSUM | P-value | Ankh | BLOSUM | P-value | Ankh | BLOSUM | P-value |
| cd00012 | **0.0446** | 0.1472 | 9.3E-275 | **0.0235** | 0.1445 | 2.2E-262 | **0.6738** | 0.9617 | 3.5E-285 |
| cd00083 | **0.0026** | 0.0356 | 2.1E-131 | **0.0006** | 0.0141 | 1.6E-128 | **0.0430** | 0.3998 | 8.5E-133 |
| cd00173 | **0.0335** | 0.3052 | 5.1E-31 | **0.0081** | 0.2873 | 2.5E-29 | **0.4884** | 0.7891 | 7.2E-34 |
| cd00637 | **0.0044** | 0.0179 | 0.0E+00 | **0.0010** | 0.0053 | 0.0E+00 | **0.1315** | 0.4094 | 0.0E+00 |
| cd01040 | **0.0037** | 0.0338 | 1.8E-298 | **0.0007** | 0.0099 | 7.6E-303 | **0.1006** | 0.4217 | 4.9E-302 |
| cd01068 | **0.0027** | 0.0334 | 2.6E-27 | **0.0006** | 0.0079 | 8.5E-29 | **0.1046** | 0.4230 | 6.9E-29 |
| cd10140 | **0.0596** | 0.0859 | 3.0E-85 | **0.0120** | 0.0367 | 2.7E-90 | **0.7260** | 0.8695 | 2.6E-55 |
| cd21116 | **0.0084** | 0.0456 | 1.7E-51 | **0.0029** | 0.0231 | 1.3E-54 | **0.2944** | 0.6259 | 8.8E-55 |
| Domains won | **8** | 0 |  | **8** | 0 |  | **8** | 0 |  |
| Tests won |  |  |  |  |  |  |  |  |  |
|  | Ankh | BLOSUM | Equal | Ankh | BLOSUM | Equal | Ankh | BLOSUM | Equal |
| cd00012 | **1699** | 70 | 1 | **1639** | 130 | 1 | **1723** | 17 | 30 |
| cd00083 | **870** | 45 | 166 | **864** | 40 | 177 | **866** | 32 | 183 |
| cd00173 | **226** | 30 | 69 | **225** | 31 | 69 | **230** | 22 | 73 |
| cd00637 | **2996** | 7 | 0 | **2960** | 43 | 0 | **3002** | 1 | 0 |
| cd01040 | **1813** | 58 | 20 | **1821** | 45 | 25 | **1814** | 41 | 36 |
| cd01068 | **162** | 4 | 5 | **164** | 2 | 5 | **165** | 1 | 5 |
| cd10140 | **541** | 86 | 3 | **565** | 62 | 3 | **451** | 101 | 78 |
| cd21116 | **313** | 12 | 0 | **319** | 6 | 0 | **320** | 5 | 0 |
| Total | **8620** | 312 | 264 | **8557** | 359 | 280 | **8571** | 220 | 405 |
| Tests won (%) |  |  |  |  |  |  |  |  |  |
| cd00012 | **95.99** | 3.95 | 0.06 | **92.60** | 7.34 | 0.06 | **97.34** | 0.96 | 1.69 |
| cd00083 | **80.48** | 4.16 | 15.36 | **79.93** | 3.70 | 16.37 | **80.11** | 2.96 | 16.93 |
| cd00173 | **69.54** | 9.23 | 21.23 | **69.23** | 9.54 | 21.23 | **70.77** | 6.77 | 22.46 |
| cd00637 | **99.77** | 0.23 | 0.00 | **98.57** | 1.43 | 0.00 | **99.97** | 0.03 | 0.00 |
| cd01040 | **95.88** | 3.07 | 1.06 | **96.30** | 2.38 | 1.32 | **95.93** | 2.17 | 1.90 |
| cd01068 | **94.74** | 2.34 | 2.92 | **95.91** | 1.17 | 2.92 | **96.49** | 0.58 | 2.92 |
| cd10140 | **85.87** | 13.65 | 0.48 | **89.68** | 9.84 | 0.48 | **71.59** | 16.03 | 12.38 |
| cd21116 | **96.31** | 3.69 | 0.00 | **98.15** | 1.85 | 0.00 | **98.46** | 1.54 | 0.00 |
| Average | **89.82** | 5.04 | 5.14 | **90.05** | 4.66 | 5.30 | **88.83** | 3.88 | 7.29 |

The full results are summarized by the dot plots in Fig. 4. It can be seen from the plots that Ankh-score clearly dominates the best BLOSUM scores for each case.

**Fig. 4.**
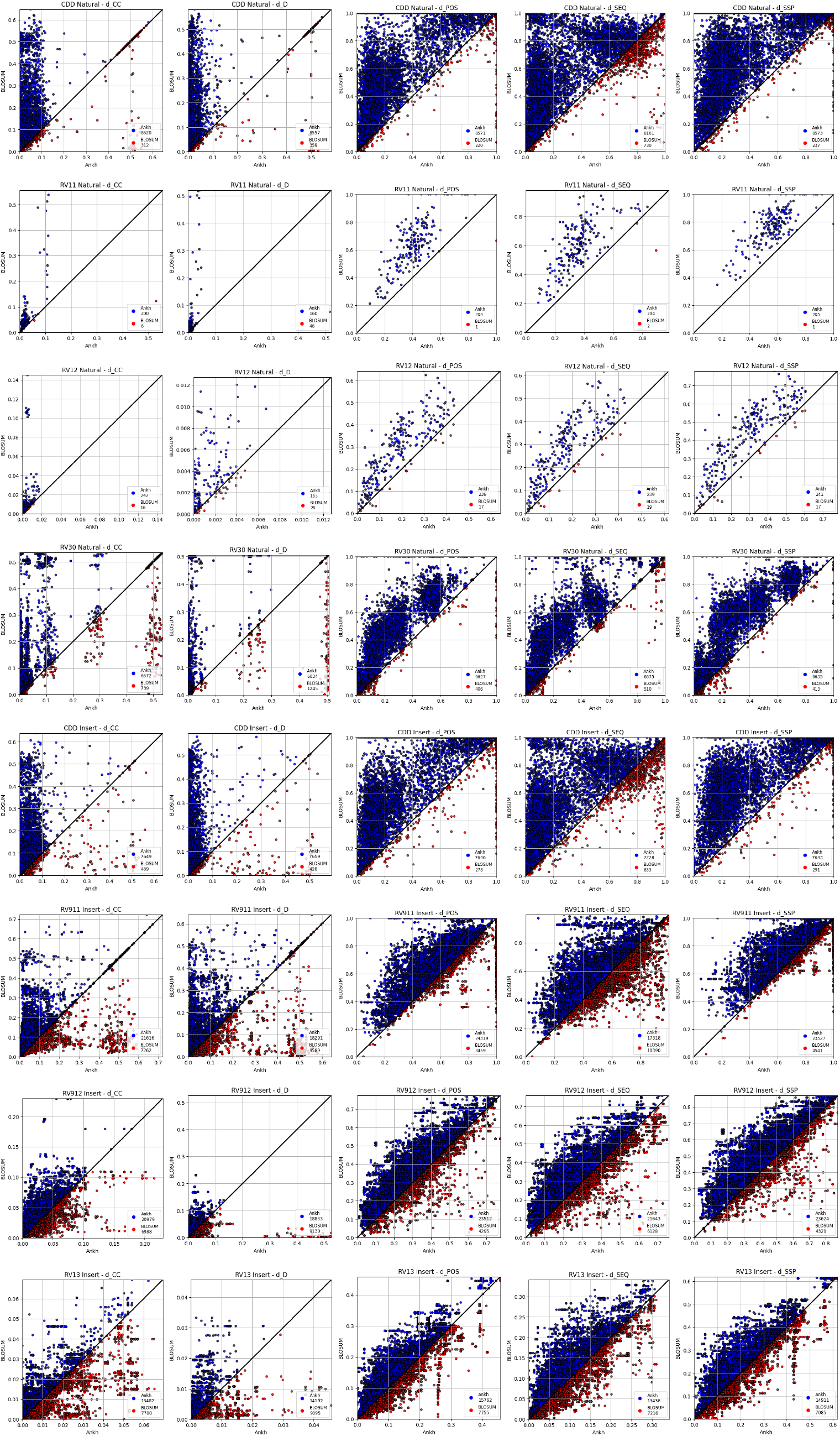
Ankh-score vs best BLOSUM. One row for each natural dataset gives the distribution of each of the five distances, for the alignments produced using Ankh-score (blue) and the best BLOSUM matrix for each test (red). The legend gives the number of tests won by each.

We give also the summaries for the number of domains won by each in Table 4 and the average percentage of the tests won for each dataset in Table 5. The number of BLOSUM matrices, by dataset, achieving the best results is given in Table 6. As expected, more higher order BLOSUM matrices produce the best performance for datasets with higher amino acid identity level.

**Table 4.**
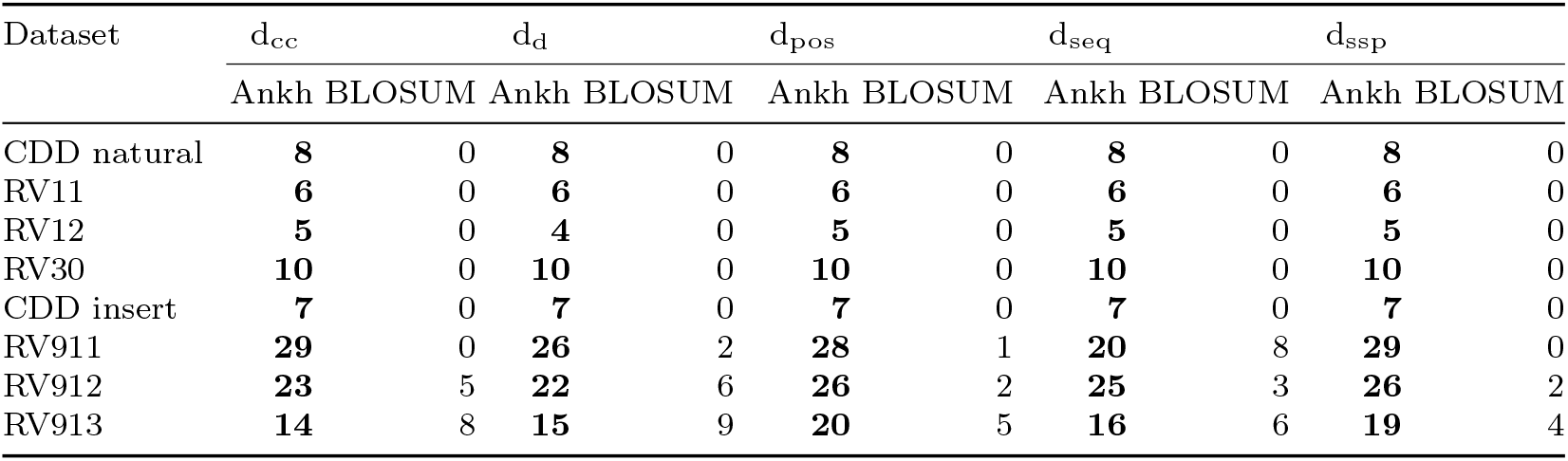
Summary of Ankh-score vs best BLOSUM for all datasets: number of domains won by each method for each dataset.

**Table 5.** Summary of Ankh-score vs best BLOSUM for all datasets: number of domains won by each method for each dataset.

| Dataset | d<sub>cc</sub> |  |  | d<sub>d</sub> |  |  | d<sub>pos</sub> |  |  | d<sub>seq</sub> |  |  | d<sub>ssp</sub> |  |  |
| --- | --- | --- | --- | --- | --- | --- | --- | --- | --- | --- | --- | --- | --- | --- | --- |
|  | Ankh | BLSM | Equal | Ankh | BLSM | Equal | Ankh | BLSM | Equal | Ankh | BLSM | Equal | Ankh | BLSM | Equal |
| CDDn | **89.82** | 5.04 | 5.14 | **90.05** | 4.66 | 5.30 | **88.83** | 3.88 | 7.29 | **84.77** | 9.53 | 5.71 | **88.81** | 4.20 | 6.99 |
| RV11 | **96.25** | 3.75 | 0.00 | **85.62** | 14.38 | 0.00 | **98.41** | 0.79 | 0.79 | **98.41** | 1.59 | 0.00 | **99.21** | 0.79 | 0.00 |
| RV12 | **78.67** | 10.56 | 10.76 | **72.82** | 16.04 | 11.14 | **80.30** | 5.13 | 14.57 | **80.64** | 6.82 | 12.54 | **80.33** | 5.22 | 14.45 |
| RV30 | **75.73** | 12.54 | 11.74 | **70.09** | 17.79 | 12.12 | **77.53** | 7.48 | 14.99 | **76.71** | 9.22 | 14.07 | **77.67** | 7.81 | 14.52 |
| CDDi | **89.04** | 10.14 | 0.81 | **90.17** | 9.03 | 0.81 | **88.56** | 7.19 | 4.25 | **83.95** | 14.76 | 1.29 | **88.54** | 7.62 | 3.84 |
| RV911 | **74.54** | 25.04 | 0.42 | **65.33** | 34.25 | 0.43 | **83.86** | 11.93 | 4.21 | **61.85** | 37.11 | 1.04 | **81.13** | 15.66 | 3.21 |
| RV912 | **74.93** | 24.96 | 0.12 | **67.26** | 32.61 | 0.13 | **83.97** | 15.34 | 0.69 | **77.30** | 21.89 | 0.82 | **84.37** | 15.43 | 0.20 |
| RV913 | **61.28** | 35.00 | 3.72 | **58.76** | 37.90 | 3.35 | **63.05** | 31.02 | 5.93 | **61.07** | 32.80 | 6.13 | **64.83** | 30.80 | 4.37 |

**Table 6.** Best BLOSUMs. The number of BLOSUM matrices, by dataset, achieving the best results.

| Dataset | BLOSUM45 | BLOSUM50 | BLOSUM62 | BLOSUM80 | BLOSUM90 |
| --- | --- | --- | --- | --- | --- |
| CDD natural | 30 | 5 | 5 | 0 | 0 |
| RV11 | 27 | 18 | 0 | 0 | 0 |
| RV12 | 10 | 1 | 14 | 0 | 0 |
| RV30 | 34 | 10 | 4 | 2 | 0 |
| CDD insert | 34 | 4 | 2 | 0 | 0 |
| RV911 | 90 | 23 | 26 | 3 | 3 |
| RV912 | 76 | 17 | 44 | 2 | 1 |
| RV913 | 42 | 28 | 55 | 3 | 7 |
| Total | 343 | 106 | 150 | 10 | 11 |

### Comparison against GPCRtm

We compared first the GPCRtm matrix with the best BLOSUM matrix and GPCRtm is the winner; see Supplementary Tables 31 and 32. A more fair comparison is against a single BLOSUM matrix. As seen in Supplementary Table 32, BLOSUM62 is most often the best matrix. The comparison between GPCRtm and BLOSUM62 is given in Supplementary Table 33 and indicates GPCRtm as a very clear winner. Finally, we compared Ankh-score against GPCRtm in Supplementary Table 34 and it wins by a very large margin.

### Comparison against ProtT5-score and PEbA

We already mentioned that the PEbA and ProtT5-score produce the same results. ProtT5-score is the closest competitor, therefore we compared Ankh-score against ProtT5-score on all eight datasets, natural and insert. The dot plots are shown in Supplementary Fig. 1 and all they indicate is that the performances are close. We need the actual numbers to decide the winner. Full comparison is given in Supplementary Tables 34-42. We give here the summaries for the number of domains won by each in Table 7 and the average percentage of the tests won for each dataset in Table 8. The numbers present a very interesting picture. Ankh-score is very clearly the winner for the natural datasets, which are the most relevant. For the insert data, Ankh-score is still winning, but not as clearly. This behaviour is considered further in the Discussion section.

**Table 7.**
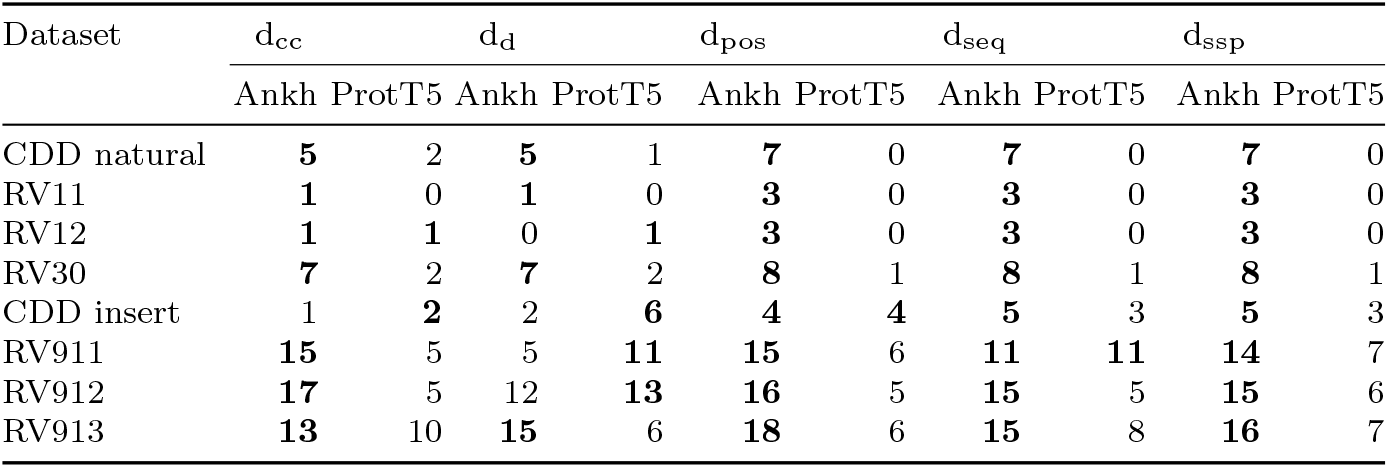
Summary of Ankh-score vs ProtT5-score for all datasets: number of domains won by each method for each dataset.

| Dataset | d<sub>cc</sub> |  | d<sub>d</sub> |  | d<sub>pos</sub> |  | d<sub>seq</sub> |  | d<sub>ssp</sub> |  |
| --- | --- | --- | --- | --- | --- | --- | --- | --- | --- | --- |
|  | Ankh | ProtT5 | Ankh | ProtT5 | Ankh | ProtT5 | Ankh | ProtT5 | Ankh | ProtT5 |
| CDD natural | **5** | 2 | **5** | 1 | **7** | 0 | **7** | 0 | **7** | 0 |
| RV11 | **1** | 0 | **1** | 0 | **3** | 0 | **3** | 0 | **3** | 0 |
| RV12 | **1** | **1** | 0 | **1** | **3** | 0 | **3** | 0 | **3** | 0 |
| RV30 | **7** | 2 | **7** | 2 | **8** | 1 | **8** | 1 | **8** | 1 |
| CDD insert | 1 | **2** | 2 | **6** | **4** | **4** | **5** | 3 | **5** | 3 |
| RV911 | **15** | 5 | 5 | **11** | **15** | 6 | **11** | **11** | **14** | 7 |
| RV912 | **17** | 5 | 12 | **13** | **16** | 5 | **15** | 5 | **15** | 6 |
| RV913 | **13** | 10 | **15** | 6 | **18** | 6 | **15** | 8 | **16** | 7 |

**Table 8.** Summary of Ankh-score vs ProtT5 for all datasets: number of domains won by each method for each dataset.

| Dataset | d<sub>cc</sub> |  |  | d<sub>d</sub> |  |  | d<sub>pos</sub> |  |  | d<sub>seq</sub> |  |  | d<sub>ssp</sub> |  |  |
| --- | --- | --- | --- | --- | --- | --- | --- | --- | --- | --- | --- | --- | --- | --- | --- |
|  | Ankh | ProtT5 | Equal | Ankh | ProtT5 | Equal | Ankh | ProtT5 | Equal | Ankh | ProtT5 | Equal | Ankh | ProtT5 | Equal |
| CDDn | **58.90** | 33.36 | 7.74 | **59.14** | 34.74 | 6.12 | **55.93** | 27.44 | 16.63 | **57.93** | 29.26 | 12.81 | **57.70** | 28.72 | 13.58 |
| RV11 | **78.57** | 21.43 | 0.00 | **90.00** | 10.00 | 0.00 | **74.82** | 24.08 | 1.10 | **75.27** | 23.17 | 1.56 | **74.45** | 24.36 | 1.19 |
| RV12 | 47.16 | **52.84** | 0.00 | 37.88 | **62.12** | 0.00 | **74.53** | 20.68 | 4.79 | **74.96** | 18.46 | 6.58 | **75.38** | 21.11 | 3.50 |
| RV30 | **59.24** | 30.43 | 10.33 | **60.19** | 28.86 | 10.95 | **62.62** | 22.68 | 14.70 | **62.79** | 23.36 | 13.85 | **63.17** | 23.05 | 13.78 |
| CDDi | **53.84** | 37.94 | 8.22 | **52.26** | 38.73 | 9.02 | **54.63** | 35.09 | 10.28 | **55.40** | 34.62 | 9.97 | **55.83** | 36.34 | 7.83 |
| RV911 | **54.01** | 44.76 | 1.24 | 45.68 | **53.58** | 0.74 | **55.20** | 34.83 | 9.97 | **53.13** | 42.32 | 4.55 | **55.88** | 37.71 | 6.41 |
| RV912 | **58.47** | 40.56 | 0.97 | **49.78** | 49.24 | 0.98 | **62.19** | 33.56 | 4.25 | **61.93** | 34.75 | 3.32 | **62.66** | 35.21 | 2.13 |
| RV913 | **55.18** | 35.87 | 8.95 | **49.55** | 38.78 | 11.67 | **54.61** | 31.47 | 13.92 | **56.53** | 31.93 | 11.54 | **57.17** | 32.38 | 10.44 |

### Comparison against DEDAL

The comparison of Ankh-score against DEDAL is given as dot plots in Supplementary Fig. 2 for all natural datasets, with full details in Supplementary Tables 43-46 and they show that Ankh-score is clearly the winner. The comparisons indicate a similar situation as with BLOSUM matrices, therefore we have not performed the comparison using the insert data.

### Comparison against BLOSUM on common overlap

In the light of our results for missed alignments in Fig. 2 and considering that our rigorous comparison of alignments penalizes for such missed alignment segments, we need to evaluate whether the advantage of Ankh-score over BLOSUM matrices is due to this penalization or actual alignment quality. We compared Ankh-score with the commonly best matrix, BLOSUM45, on the common alignments. That is, the alignments were first computed by each method and only the intersection among both computed alignments and the reference alignment was considered for comparison. (Note that we need to first fix a BLOSUM matrix in order to be able to compute the common alignments.) The results are given in Supplementary Table 47 and they indicate fairly similar performance with the original comparison.

### Protein embeddings improvements

We have tried to improve the performance of the embedding vectors by combining them in various ways. We obtained a successful combination between ProtT5 and ESM2 models, using the average of the normalized vectors. As shown in Supplementary Table 48, ProtT5+ESM2-score is better than ProtT5-score. Ankh-score still outperforms the combination quite clearly, as seen in Supplementary Table 49. Interestingly, Ankh does not appear to combine well with other methods.

## Discussion

We have compared the new algorithm, that uses Ankh-score, rigorously using a large number of tests against the current standard of BLOSUM matrices, as well as the closest competitor, ProtT5-score/PEbA. The datasets include natural data, where domains occur naturally inside protein sequences, and insert data, where we insert domains inside random protein sequences. This was done to cover a wide range of amino acid identity levels, 8-66%; see Table 1. The tests were evaluated using the Wilcoxon signed-rank test for statistical significance, with the p-value threshold set conservatively at 0.01.

Ankh-score shows far better performance than the best BLOSUM matrices, winning all natural data cases. Only for three domains of the BALiBASE RV11 dataset and a single case of BALiBASE RV12, the p-values are not significant, shown in red in Supplementary Tables 17 and 19. Of these, only two cases have p-value above 0.05. Ankh-score performs much better than best BLOSUM matrices also for the insert data, the winning margin becoming smaller for the cases with higher identity. This is to be expected as BLOSUM matrices are good at handling higher identity cases, as there are fewer divergent regions and the contextual and long-range dependencies captured by Ankh are less critical. This is captured well by the top row of Fig. 5. The plots indicate the percentage of wins of Ankh-score over best BLOSUM for, from left to right, natural, insert, and all data. The difference is the largest for natural data, then decreases for insert data and with increasing identity level. Importantly, the performance of Ankh-score stays higher than that of the best BLOSUM matrices at all identity levels tested. Note also the different identity range of the first plot. The magnitude of the advantage of Ankh-score is shown in the top row of Fig. 6; note the different range of both the abscissa and ordinate axes for the first plot.

**Fig. 5.**
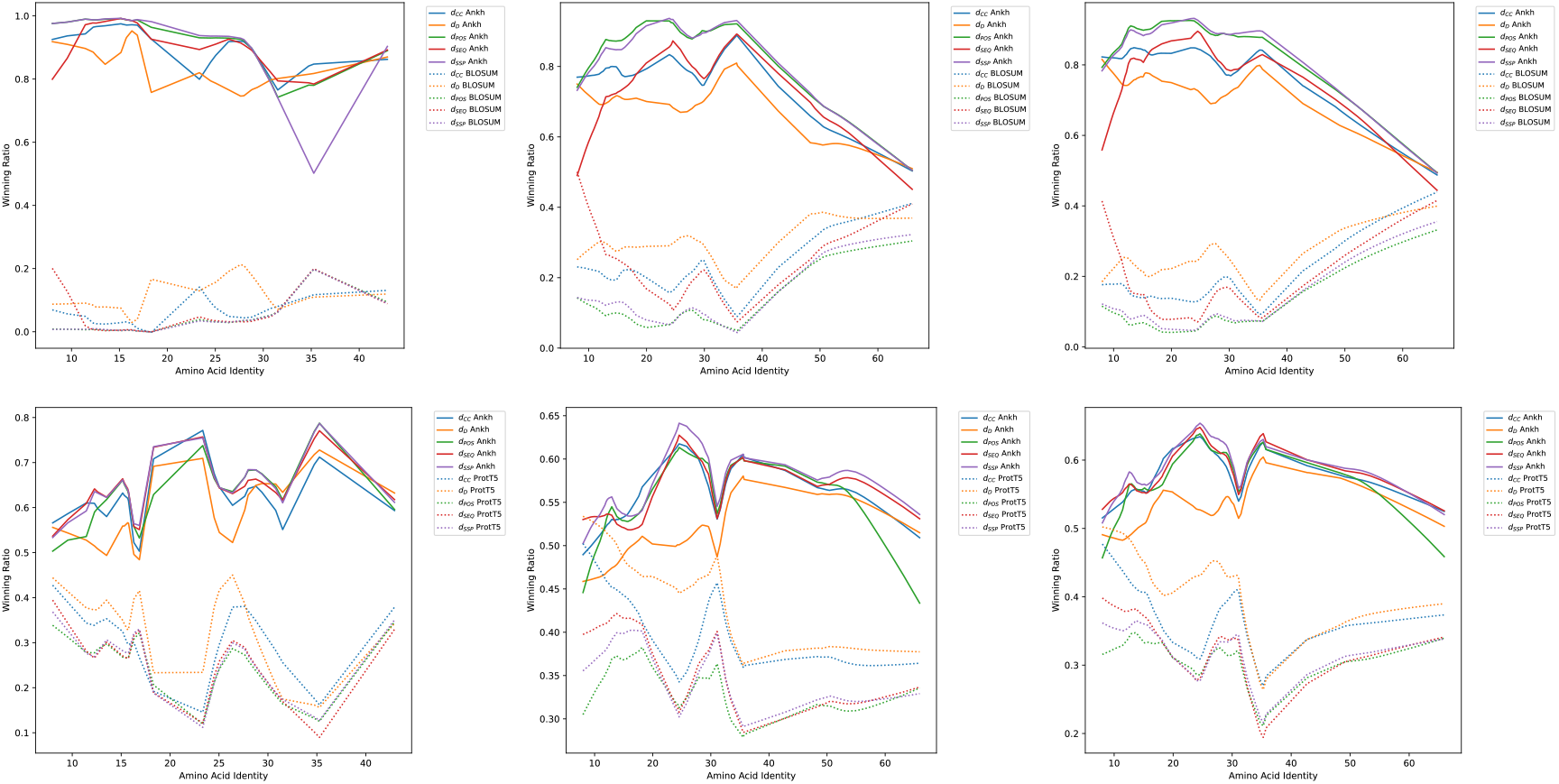
Wins percentage over identity level. The plots show the percentage of wins of Ankh-score vs best BLOSUM (top row) and ProtT5-score (bottom row), for natural (left), insert (middle), and all data (right). Two curves are given for each of the five distances, one solid curve for Ankh-score and a dotted of the same colour for the competitor.

**Fig. 6.**
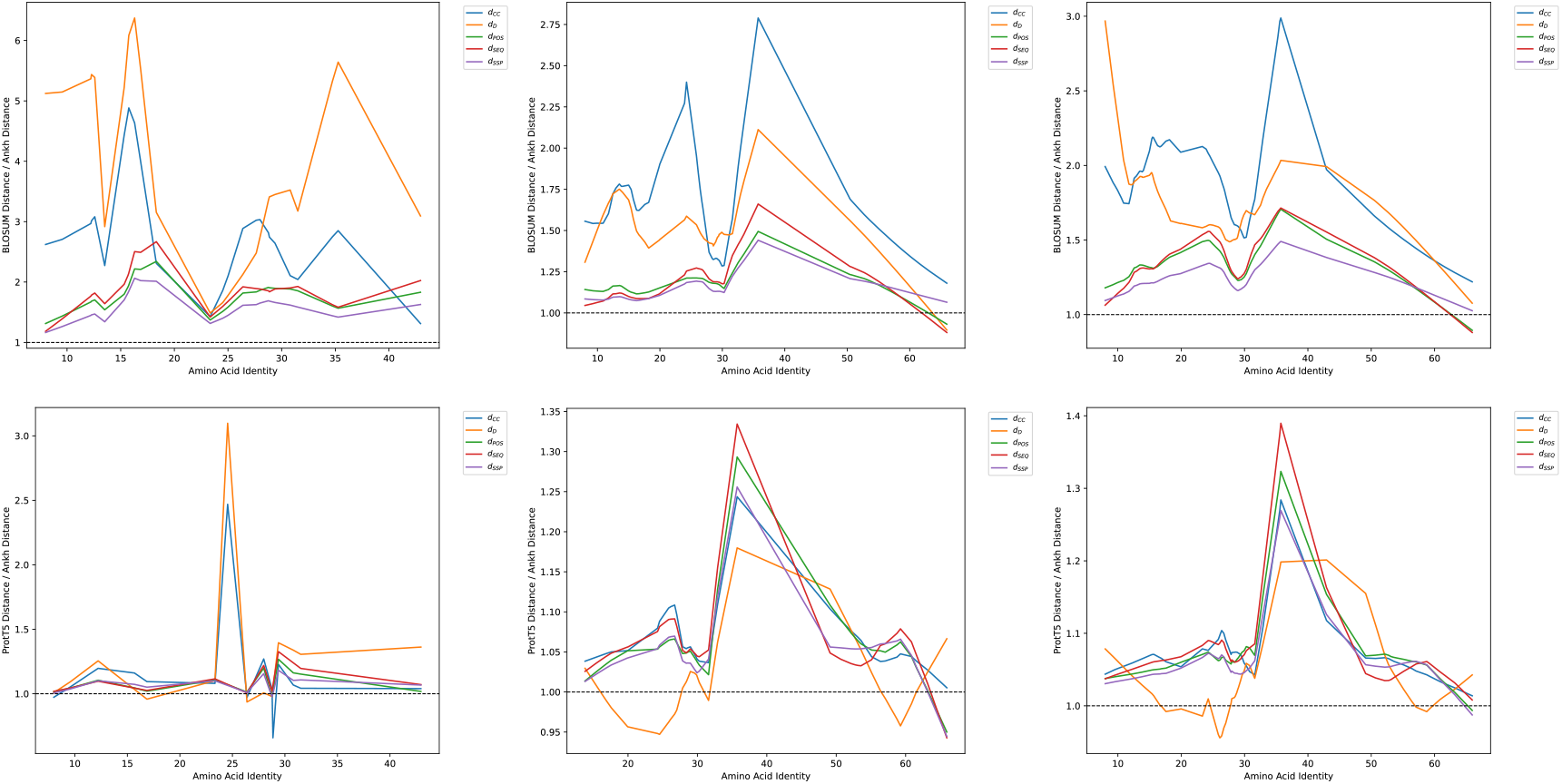
Distance ratio over identity level. The plots show the distance ratio, for all five distance, between Ankh-score vs best BLOSUM (top row) and ProtT5-score (bottom row), for natural (left), insert (middle), and all data (right). Ratio higher than 1 means Ankh-score is better than the competitor.

Compared against the closest competitor, Ankh-score performs, remarkably, consistently better than ProtT5-score. The number of cases that are not statistically significant is much higher than in the BLOSUM comparison – see the p-values in red in Supplementary Tables 35-42. Ankh-score dominates ProtT5-score at all identity levels, as seen in the bottom row of Fig. 5. The magnitude of the advantage is shown in the bottom row of Fig. 6.

Of the five distances considered, the last three, d_pos_, d_seq_, d_ssp_, are true metric distances built on the sum-of-pairs idea. The difference is that d_ssp_ ignores gaps, d_seq_ considers gaps labelled by sequences, whereas d_pos_ labels gaps by both sequence and position. Therefore, d_pos_ is the most relevant of the three, however their behaviour is fairly consistent. The other two distances are very different and they exhibit different behaviour from the three, as well as from each other. The five distances together give a better general picture of the comparison. As seen in Tables 7 and 8, Ankh-score is winning everywhere for the last three distance, but ProtT5-score has some wins for the first two distance, particularly d_d_. The full meaning of this behaviour is yet to be determined.

Our results have interesting implications on the behaviour of the two models, Ankh and ProtT5, themselves. Comparing the first two plots of the bottom row of Fig. 5, we see that the difference is much smaller between the Ankh and ProtT5 sets of curves, indicating that Ankh behaves significantly better for the natural domains, compared with insert ones. This phenomenon is most visible when comparing the ‘CDD natural’ and ‘CDD insert’ rows of Table 7. A possible interpretation is as follows. The embedding models are trained in unsupervised manner on very large number of (natural) protein sequences. It is therefore to be expected that the insert sequences may confuse the models somewhat, as being unnatural. Our results show that Ankh is significantly more confused than ProtT5 is. This can be interpreted as indication that Ankh better captures the true “language” of proteins.

## Conclusions

The methods discussed here are a massive improvement for the analysis of biological sequences. The reconstruction of accurate protein alignments are a first step and a basic step in almost all studies. Alignments are necessary to detect homologues between species and to employ comparative data. These, in turn, aid structural predictions, functional annotations, and domain identification. Active site identification and annotation of pathogenic mutations rely on precise alignments. This knowledge contributes to deeper insights in areas like drug design and disease diagnostics. The detection of remote homologs is a prerequisite for evolutionary studies.

Accurate local protein sequence alignments are the foundation of virtually every bioinformatics and comparative-genomics analysis. A global alignment is restricted to end-to-end alignments regardless of the homology present. The ends of sequences are known to evolve rapidly with little selective constraint. This leads to fragments of irrelevant sequences at the ends of sequences and hinders the correct alignment of subsequences. Similarly, proteins are often composed of multiple functional domains.

But their alignment is needlessly complicated by global alignments that must match the entire sequence rather than local alignments that align just the functional domains.

Most alignment methods use a scoring matrix to evaluate the differences in potential alignments. The most commonly used of these matrices is the empirically determined BLOSUM matrix. This is a static matrix that was determined decades ago from active site blocks. All such empirical matrices suffer from amino acid biases dues to over/under-represented amino acids in the training blocks. BLOSUM assumes uniform substitution rates and assumes no context dependencies.

Here we have used embedding vectors from every known protein sequence. These vectors capture far more of the biology than just what amino acid is present. It includes context for each amino acid and knowledge from similar proteins as well as from distantly related proteins. This information reflects not just amino acid presence but substitution patterns and selective constraints.

Moving beyond generic substitution matrices allows alignments to more faithfully reflect the true, context-dependent evolutionary and structural constraints acting on proteins, thereby enhancing every downstream inference.

The alignments using embedding vectors clearly provide alignments that are as good as the standard methods used and, in almost all cases, far superior. This is due to embedding vectors providing a vast amount of information that relate the biology to the alignment. Alignments are constrained to by the function of the protein and by the phylogenetic history between the two species. With a slightly different function, the proteins can explore more of sequence space without compromising their function. With a great divergence between the species the sequences can explore greater levels of structural divergence without a loss of function. The embedding vectors incorporate individual amino acid differences (as done in a BLOSUM matrix) but in addition, they also include contextual information including some sense of the three-dimensional structure. This information and more is combined into the embedding vector to produce alignments that respect the biological constraints and can therefore produce very accurate alignments.

The last comment concerns the interplay between structure prediction and alignments. AlphaFold (Jumper et al. 2021; Abramson et al. 2024) does a much better job at the alignment of distantly related sequences than does BLOSUM (Lesk and Konagurthu 2022; Rajapaksa et al. 2023). But embedding vectors do a similar or better job at this same problem. In addition, embedding vectors can do a better job when less distant sequences are aligned and, arguably, a better job than BLOSUM with very similar sequences. The reason for this is that embedding vectors incorporate features of the structures from the proteins from which they were constructed and also the finer features of individual amino acid differences. Furthermore, AlphaFold provides an estimate of the structure but often has levels of uncertainty in these structures that are otherwise ignored. Even with a well supported structure, such a structure is not static and can change with different environmental conditions or as the protein binds other moieties to function. Last but not least, AlphaFold relies on sequence alignments to generate its highly accurate protein structure predictions.

## Availability of data and materials

The new method, protocol, and data are freely available as a web server at e-score.csd.uwo.ca and as source code at github.com/lucian-ilie/E-score.

## Supplementary information

Supplementary information is available online: localAlign supp.pdf.

## Author contributions

J.M. selected the data, wrote the software and web server, performed all tests, including installing and running the necessary libraries, and contributed to the methodology and analysis. G.B.G. advised on choosing the data and contributed to performing and writing the biological analysis. L.I. proposed the study, designed the methodology, analyzed the results, and wrote the manuscript.

## Acknowledgements

All our computations were performed on Digital Research Alliance of Canada servers. We would like to thank Paul Moore for helping with some of the code for automating testing on the server.

## Declarations

### Ethics approval and consent to participate

Not applicable.

## Consent for publication

Not applicable.

