## Supplemental material for "Protein Embeddings and Local Alignments"

– Supplementary material –

Julia Malec, G. Brian Golding, Lucian Ilie*

May 17, 2025

### 1 Datasets

#### 1.1 CDD

Supplementary Table 1: CDD natural dataset.

| Domain | | Seq. | Tests | Protein len. | Domain len. | Seq. before | Seq. after | Identity (%) |
|---|---|---|---|---|---|---|---|---|
| *NBD_sugar-kinase_HSP70_actin* | cd00012 | 60 | 1770 | 398.07 | 316.18 | 33.37 | 48.52 | 8.00 |
| *bHLH_SF* | cd00083 | 47 | 1081 | 537.87 | 47.36 | 287.45 | 203.06 | 24.55 |
| *SH2* | cd00173 | 26 | 325 | 701.23 | 82.00 | 259.92 | 359.31 | 15.76 |
| *7tm_classA_rhodopsin-like* | cd00637 | 78 | 3003 | 388.44 | 283.62 | 35.05 | 69.77 | 16.86 |
| *Mb-like* | cd01040 | 62 | 1891 | 211.84 | 133.60 | 49.84 | 28.40 | 15.68 |
| *Globin_sensor* | cd01068 | 19 | 171 | 394.95 | 145.47 | 16.32 | 233.16 | 16.30 |
| *PFM_aerolysin_family* | cd10140 | 36 | 630 | 386.08 | 155.81 | 173.31 | 56.97 | 9.56 |
| *ClyA-like* | cd21116 | 26 | 325 | 399.31 | 318.77 | 25.23 | 55.31 | 12.22 |
| Total / Avg. | | 354 | 9196 | 427.22 | 185.35 | 110.06 | 131.81 | 8.00% - 24.55% |

Supplementary Table 2: CDD insert dataset.

| Domain | | Seq. | Tests | Protein len. | Domain len. | Seq. before | Seq. after | Identity (%) |
|---|---|---|---|---|---|---|---|---|
| *NBD_sugar-kinase_HSP70_actin* | cd00012 | 60 | 1770 | 453.60 | 316.18 | 69.61 | 67.80 | 8.00 |
| *bHLH_SF* | cd00083 | 47 | 1081 | 185.30 | 47.36 | 67.42 | 70.52 | 24.55 |
| *SH2* | cd00173 | 26 | 325 | 220.82 | 82.00 | 73.38 | 65.44 | 15.76 |
| *7tm_classA_rhodopsin-like* | cd00637 | 78 | 3003 | 421.95 | 283.62 | 68.95 | 69.39 | 16.86 |
| *Mb-like* | cd01040 | 62 | 1891 | 271.43 | 133.60 | 69.36 | 68.47 | 15.68 |
| *Globin_sensor* | cd01068 | 19 | 171 | 284.77 | 145.47 | 66.64 | 72.65 | 16.30 |
| *PFM_aerolysin_family* | cd10140 | 36 | 630 | 293.22 | 155.81 | 68.15 | 69.26 | 9.56 |
| *ClyA-like* | cd21116 | 26 | 325 | 455.74 | 318.77 | 66.21 | 70.76 | 12.22 |
| Total / Avg. | | 354 | 9196 | 323.35 | 185.35 | 68.72 | 69.29 | 8.00% - 24.55% |

#### 1.2 BALiBASE

Supplementary Table 3: BALiBASE RV11 original data.

| Domain | Seq. | Protein len. | Domain len. | Seq. before | Seq. after |
|---|---|---|---|---|---|
| BBS11001 | 4 | 86.25 | 76.50 | 3.75 | 6.00 |
| BBS11002 | 8 | 98.00 | 60.50 | 31.63 | 5.88 |
| BBS11003 | 4 | 470.00 | 432.75 | 25.25 | 12.00 |
| BBS11004 | 4 | 421.00 | 381.75 | 14.00 | 25.25 |
| BBS11005 | 14 | 398.71 | 333.14 | 60.00 | 5.57 |
| BBS11006 | 8 | 244.75 | 205.88 | 27.00 | 11.88 |
| BBS11007 | 9 | 418.33 | 394.22 | 6.78 | 17.33 |
| BBS11008 | 4 | 305.00 | 109.50 | 192.50 | 3.00 |
| **BBS11009** | **4** | **216.00** | **91.00** | **59.75** | **65.25** |
| **BBS11010** | **4** | **491.50** | **178.25** | **37.75** | **275.50** |
| BBS11011 | 5 | 201.00 | 176.80 | 13.00 | 11.20 |
| BBS11012 | 4 | 364.00 | 328.50 | 9.50 | 26.00 |
| BBS11013 | 5 | 71.60 | 52.60 | 8.40 | 10.60 |
| BBS11014 | 6 | 539.50 | 509.83 | 23.50 | 6.17 |
| BBS11015 | 4 | 308.50 | 303.00 | 1.75 | 3.75 |
| **BBS11016** | **8** | **470.50** | **324.13** | **57.00** | **89.38** |
| BBS11017 | 4 | 252.75 | 234.50 | 0.75 | 17.50 |
| **BBS11018** | **14** | **547.93** | **432.79** | **73.71** | **41.43** |
| BBS11019 | 10 | 348.40 | 344.80 | 2.10 | 1.50 |
| BBS11020 | 9 | 219.22 | 203.44 | 4.78 | 11.00 |
| BBS11021 | 4 | 117.25 | 100.25 | 9.25 | 7.75 |
| BBS11022 | 4 | 118.00 | 61.50 | 40.00 | 16.50 |
| BBS11023 | 7 | 309.86 | 268.00 | 1.71 | 40.14 |
| **BBS11024** | **4** | **407.00** | **302.75** | **36.50** | **67.75** |
| BBS11025 | 4 | 82.75 | 60.00 | 7.00 | 15.75 |
| **BBS11026** | **7** | **240.00** | **83.29** | **34.43** | **122.29** |
| BBS11027 | 7 | 232.57 | 195.71 | 2.29 | 34.57 |
| BBS11028 | 10 | 158.80 | 89.20 | 18.40 | 51.20 |
| BBS11029 | 4 | 102.00 | 91.50 | 9.25 | 1.25 |
| BBS11030 | 14 | 291.79 | 184.21 | 3.57 | 104.00 |
| BBS11031 | 11 | 376.27 | 313.73 | 39.36 | 23.18 |
| BBS11032 | 8 | 294.75 | 229.25 | 36.50 | 29.00 |
| BBS11033 | 11 | 151.27 | 92.91 | 53.82 | 4.55 |
| **BBS11034** | **8** | **491.63** | **336.75** | **56.88** | **98.00** |
| BBS11035 | 5 | 97.40 | 89.20 | 6.80 | 1.40 |
| BBS11036 | 8 | 370.00 | 333.13 | 1.88 | 35.00 |
| **BBS11037** | **5** | **523.40** | **367.80** | **83.60** | **72.00** |
| **BBS11038** | **8** | **364.25** | **268.13** | **49.63** | **46.50** |

Supplementary Table 4: BALiBASE RV11 dataset; includes domains with sequences having, on the average, at least 30 residues before and after domain occurrences.

| Domain | Seq. | Tests | Protein len. | Domain len. | Seq. before | Seq. after | Identity (%) |
|---|---|---|---|---|---|---|---|
| BBS11009 | 4 | 6 | 216.00 | 91.00 | 59.75 | 65.25 | 18.28 |
| BBS11010 | 4 | 6 | 491.50 | 178.25 | 37.75 | 275.50 | 15.16 |
| BBS11016 | 8 | 28 | 470.50 | 324.13 | 57.00 | 89.38 | 15.74 |
| BBS11018 | 14 | 91 | 547.93 | 432.79 | 73.71 | 41.43 | 13.51 |
| BBS11024 | 4 | 6 | 407.00 | 302.75 | 36.50 | 67.75 | 11.44 |
| BBS11026 | 7 | 21 | 240.00 | 83.29 | 34.43 | 122.29 | 12.58 |
| BBS11034 | 8 | 28 | 491.63 | 336.75 | 56.88 | 98.00 | 15.33 |
| BBS11037 | 5 | 10 | 523.40 | 367.80 | 83.60 | 72.00 | 12.28 |
| BBS11038 | 8 | 28 | 364.25 | 268.13 | 49.63 | 46.50 | 18.32 |
| Total / Avg. | 62 | 224 | 416.91 | 264.99 | 54.36 | 97.57 | 11.44% - 18.32% |

Supplementary Table 5: BALiBASE RV12 original data.

| Domain | Seq. | Protein len. | Domain len. | Seq. before | Seq. after |
|---|---|---|---|---|---|
| BBS12001 | 11 | 458.27 | 438.55 | 16.09 | 3.64 |
| BBS12002 | 6 | 208.00 | 182.00 | 3.67 | 22.33 |
| BBS12003 | 8 | 71.25 | 62.00 | 8.00 | 1.25 |
| BBS12004 | 15 | 248.60 | 244.93 | 2.47 | 1.20 |
| BBS12005 | 9 | 209.00 | 197.44 | 7.11 | 4.44 |
| BBS12006 | 4 | 232.00 | 228.00 | 0.50 | 3.50 |
| BBS12007 | 8 | 1020.50 | 982.38 | 0.00 | 38.13 |
| **BBS12008** | **13** | **436.77** | **367.23** | **36.15** | **33.38** |
| BBS12009 | 5 | 112.60 | 67.80 | 16.60 | 28.20 |
| BBS12010 | 7 | 348.29 | 311.57 | 34.14 | 2.57 |
| **BBS12011** | **12** | **869.08** | **589.58** | **46.75** | **232.75** |
| BBS12012 | 4 | 373.50 | 309.25 | 62.25 | 2.00 |
| BBS12013 | 8 | 709.88 | 685.75 | 21.75 | 2.38 |
| BBS12014 | 9 | 97.22 | 49.33 | 25.11 | 22.78 |
| BBS12015 | 12 | 192.08 | 117.42 | 63.33 | 11.33 |
| BBS12016 | 5 | 494.20 | 105.20 | 24.80 | 364.20 |
| BBS12017 | 7 | 472.86 | 459.86 | 5.14 | 7.86 |
| BBS12018 | 4 | 833.75 | 810.50 | 20.75 | 2.50 |
| BBS12019 | 5 | 727.20 | 534.20 | 188.40 | 4.60 |
| BBS12020 | 4 | 125.50 | 123.00 | 2.00 | 0.50 |
| BBS12021 | 6 | 75.67 | 73.17 | 1.33 | 1.17 |
| BBS12022 | 5 | 250.40 | 81.20 | 14.80 | 154.40 |
| BBS12023 | 5 | 704.20 | 686.80 | 14.40 | 3.00 |
| BBS12024 | 4 | 243.25 | 233.75 | 5.75 | 3.75 |
| BBS12025 | 4 | 202.00 | 118.75 | 69.50 | 13.75 |
| BBS12026 | 18 | 267.39 | 253.78 | 8.50 | 5.11 |
| BBS12027 | 13 | 470.62 | 203.15 | 22.54 | 244.92 |
| BBS12028 | 8 | 294.50 | 228.25 | 64.63 | 1.63 |
| BBS12029 | 12 | 444.92 | 269.33 | 173.75 | 1.83 |
| **BBS12030** | **6** | **790.33** | **236.17** | **56.83** | **497.33** |
| BBS12031 | 10 | 919.20 | 917.50 | 1.40 | 0.30 |
| BBS12032 | 9 | 66.11 | 62.11 | 2.33 | 1.67 |
| **BBS12033** | **7** | **454.71** | **306.14** | **73.57** | **75.00** |
| BBS12034 | 4 | 276.25 | 274.25 | 2.00 | 0.00 |
| BBS12035 | 27 | 260.26 | 212.96 | 21.11 | 26.19 |
| BBS12036 | 7 | 209.57 | 201.29 | 4.86 | 3.43 |
| **BBS12037** | **13** | **734.85** | **480.92** | **182.69** | **71.23** |
| BBS12038 | 10 | 492.40 | 248.10 | 26.00 | 218.30 |
| BBS12039 | 11 | 103.55 | 67.00 | 8.82 | 27.73 |
| BBS12040 | 5 | 132.20 | 123.80 | 0.20 | 8.20 |
| BBS12041 | 7 | 113.57 | 106.71 | 5.71 | 1.14 |
| BBS12042 | 4 | 503.50 | 492.25 | 9.50 | 1.75 |
| BBS12043 | 34 | 318.09 | 231.50 | 81.44 | 5.15 |
| BBS12044 | 11 | 573.18 | 564.18 | 8.09 | 0.91 |

Supplementary Table 6: BALiBASE RV12 dataset; includes domains with sequences having, on the average, at least 30 residues before and after domain occurrences.

| Domain | Seq. | Tests | Protein len. | Domain len. | Seq. before | Seq. after | Identity (%) |
|---|---|---|---|---|---|---|---|
| BBS12008 | 13 | 78 | 436.77 | 367.23 | 36.15 | 33.38 | 27.63 |
| BBS12011 | 12 | 66 | 869.08 | 589.58 | 46.75 | 232.75 | 25.04 |
| BBS12030 | 6 | 15 | 790.33 | 236.17 | 56.83 | 497.33 | 35.26 |
| BBS12033 | 7 | 21 | 454.71 | 306.14 | 73.57 | 75.00 | 24.50 |
| BBS12037 | 13 | 78 | 734.85 | 480.92 | 182.69 | 71.23 | 25.01 |
| Total / Avg. | 51 | 258 | 657.15 | 396.01 | 79.20 | 181.94 | 24.50% - 35.26% |

Supplementary Table 7: BALiBASE RV30 original data.

| Domain | Seq. | Protein len. | Domain len. | Seq. before | Seq. after |
|---|---|---|---|---|---|
| **BBS30001** | **116** | **468.01** | **178.41** | **143.79** | **145.81** |
| BBS30002 | 31 | 540.68 | 471.74 | 40.77 | 28.16 |
| BBS30003 | 142 | 407.39 | 325.45 | 72.26 | 9.68 |
| **BBS30004** | **50** | **497.26** | **391.42** | **34.14** | **71.70** |
| **BBS30005** | **113** | **479.78** | **235.58** | **214.04** | **30.16** |
| **BBS30006** | **18** | **303.39** | **94.67** | **166.33** | **42.39** |
| BBS30007 | 24 | 140.29 | 100.54 | 19.25 | 20.50 |
| **BBS30008** | **36** | **526.81** | **409.36** | **70.64** | **46.81** |
| **BBS30009** | **38** | **171.45** | **67.50** | **30.05** | **73.89** |
| **BBS30010** | **50** | **655.54** | **554.74** | **68.78** | **32.02** |
| BBS30011 | 65 | 328.98 | 309.11 | 14.57 | 5.31 |
| BBS30012 | 65 | 397.37 | 316.11 | 23.17 | 58.09 |
| **BBS30013** | **86** | **656.26** | **407.91** | **167.07** | **81.28** |
| BBS30014 | 44 | 217.32 | 203.36 | 3.64 | 10.32 |
| BBS30015 | 21 | 270.48 | 103.90 | 163.71 | 2.86 |
| **BBS30016** | **37** | **291.30** | **61.16** | **148.27** | **81.86** |
| BBS30017 | 15 | 313.00 | 255.93 | 21.20 | 35.87 |
| BBS30018 | 78 | 438.59 | 291.32 | 126.09 | 21.18 |
| BBS30019 | 34 | 356.53 | 233.21 | 14.88 | 108.44 |
| BBS30020 | 56 | 151.34 | 83.14 | 47.29 | 20.91 |
| BBS30021 | 140 | 218.47 | 192.63 | 6.94 | 18.90 |
| BBS30023 | 77 | 275.55 | 240.09 | 7.56 | 27.90 |
| BBS30024 | 69 | 490.75 | 209.62 | 257.16 | 23.97 |
| BBS30025 | 74 | 141.41 | 90.57 | 36.82 | 14.01 |
| BBS30026 | 81 | 411.37 | 323.36 | 23.31 | 64.70 |
| BBS30027 | 20 | 105.50 | 91.15 | 11.05 | 3.30 |
| BBS30028 | 89 | 425.78 | 341.36 | 54.65 | 29.76 |
| BBS30029 | 98 | 514.59 | 421.66 | 24.67 | 68.26 |
| **BBS30030** | **62** | **482.53** | **271.77** | **126.84** | **83.92** |

Supplementary Table 8: BALiBASE RV30 dataset; includes domains with sequences having, on the average, at least 30 residues before and after domain occurrences.

| Domain | Seq. | Tests | Protein len. | Domain len. | Seq. before | Seq. after | Identity (%) |
|---|---|---|---|---|---|---|---|
| BBS30001 | 116 | 1000 | 468.01 | 178.41 | 143.79 | 145.81 | 31.51 |
| BBS30004 | 50 | 1000 | 497.26 | 391.42 | 34.14 | 71.70 | 30.79 |
| BBS30005 | 113 | 1000 | 479.78 | 235.58 | 214.04 | 30.16 | 29.38 |
| BBS30006 | 18 | 153 | 303.39 | 94.67 | 166.33 | 42.39 | 23.34 |
| BBS30008 | 36 | 630 | 526.81 | 409.36 | 70.64 | 46.81 | 28.77 |
| BBS30009 | 38 | 703 | 171.45 | 67.50 | 30.05 | 73.89 | 28.86 |
| BBS30010 | 50 | 1000 | 655.54 | 554.74 | 68.78 | 32.02 | 42.94 |
| BBS30013 | 86 | 1000 | 656.26 | 407.91 | 167.07 | 81.28 | 26.39 |
| BBS30016 | 37 | 666 | 291.30 | 61.16 | 148.27 | 81.86 | 34.73 |
| BBS30030 | 62 | 1000 | 482.53 | 271.77 | 126.84 | 83.92 | 27.98 |
| Total / Avg. | 606 | 8152 | 453.23 | 267.25 | 117.00 | 68.98 | 23.34% - 42.94% |

Supplementary Table 9: BALiBASE RV911 dataset.

| Domain | Seq. | Tests | Protein len. | Domain len. | Seq. before | Seq. after | Identity (%) | Orig. before | Orig. after |
|---|---|---|---|---|---|---|---|---|---|
| BOX001 | 18 | 1000 | 1286.52 | 1133.22 | 106.79 | 164.04 | 12.47 | 0.00 | 0.00 |
| BOX022 | 13 | 1000 | 672.53 | 541.92 | 79.97 | 178.13 | 13.45 | 0.00 | 0.00 |
| BOX032 | 6 | 1000 | 1061.21 | 923.17 | 71.14 | 90.60 | 14.74 | 0.00 | 0.00 |
| BOX034 | 7 | 1000 | 617.28 | 482.29 | 110.58 | 77.55 | 15.49 | 0.00 | 0.00 |
| BOX035 | 9 | 1000 | 1161.55 | 1021.33 | 178.39 | 174.38 | 14.04 | 0.00 | 0.00 |
| BOX046 | 7 | 1000 | 990.41 | 835.57 | 376.58 | 156.90 | 15.61 | 0.00 | 0.00 |
| BOX060 | 9 | 1000 | 530.85 | 388.00 | 83.64 | 115.43 | 10.64 | 0.00 | 0.00 |
| BOX063 | 5 | 1000 | 1607.36 | 1473.80 | 70.96 | 101.91 | 24.26 | 0.00 | 0.00 |
| BOX076 | 26 | 1000 | 595.99 | 462.38 | 101.26 | 67.14 | 10.55 | 0.00 | 0.00 |
| BOX096 | 12 | 1000 | 927.28 | 793.17 | 83.09 | 166.07 | 10.87 | 0.00 | 0.00 |
| BOX115 | 7 | 1000 | 885.03 | 750.00 | 80.11 | 111.18 | 9.94 | 0.00 | 0.00 |
| BOX121 | 13 | 1000 | 702.04 | 563.62 | 180.01 | 73.53 | 13.84 | 0.00 | 0.00 |
| BOX122 | 7 | 1000 | 744.28 | 607.14 | 144.32 | 67.80 | 18.20 | 0.00 | 0.00 |
| BOX123 | 5 | 1000 | 693.95 | 555.80 | 104.68 | 85.13 | 15.68 | 0.00 | 0.00 |
| BOX142 | 5 | 1000 | 658.49 | 529.20 | 109.93 | 179.33 | 11.99 | 0.00 | 0.00 |
| BOX172 | 6 | 1000 | 566.36 | 431.00 | 72.86 | 76.00 | 9.27 | 0.00 | 0.00 |
| BOX175 | 5 | 1000 | 674.18 | 536.20 | 95.47 | 183.23 | 15.39 | 0.00 | 0.00 |
| BOX177 | 11 | 1000 | 556.30 | 419.18 | 94.15 | 68.01 | 10.92 | 0.00 | 0.00 |
| BOX180 | 24 | 1000 | 379.23 | 243.04 | 97.97 | 72.44 | 13.81 | 0.00 | 0.00 |
| BOX181 | 7 | 1000 | 362.83 | 226.57 | 75.05 | 100.11 | 17.59 | 0.00 | 0.00 |
| BOX192 | 13 | 1000 | 512.67 | 377.85 | 82.04 | 118.26 | 14.99 | 0.00 | 0.00 |
| BOX212 | 8 | 1000 | 532.15 | 397.00 | 71.10 | 111.09 | 12.00 | 0.00 | 0.00 |
| BOX214 | 37 | 1000 | 630.00 | 493.65 | 268.06 | 102.56 | 12.96 | 0.00 | 0.00 |
| BOX222 | 69 | 1000 | 1065.50 | 926.51 | 182.10 | 153.12 | 11.76 | 0.00 | 0.00 |
| BOX240 | 13 | 1000 | 760.17 | 608.23 | 179.10 | 159.26 | 14.93 | 0.00 | 0.00 |
| BOX246 | 22 | 1000 | 1072.67 | 933.64 | 215.97 | 206.37 | 18.15 | 0.00 | 0.00 |
| BOX258 | 6 | 1000 | 668.43 | 528.17 | 159.72 | 82.01 | 19.96 | 0.00 | 0.00 |
| BOX270 | 47 | 1000 | 1178.90 | 1045.55 | 257.31 | 254.14 | 16.22 | 0.00 | 0.00 |
| BOX284 | 6 | 1000 | 536.09 | 396.67 | 73.33 | 74.57 | 13.52 | 0.00 | 0.00 |
| Total / Avg. | 423 | 29000 | 780.35 | 642.20 | 131.23 | 123.11 | 9.27% - 24.26% | 0.00 | 0.00 |

Supplementary Table 10: BALiBASE RV912 dataset.

| Domain | Seq. | Tests | Protein len. | Domain len. | Seq. before | Seq. after | Identity (%) | Orig. before | Orig. after |
|---|---|---|---|---|---|---|---|---|---|
| BOX011 | 5 | 1000 | 616.85 | 479.80 | 90.86 | 71.67 | 26.09 | 0.00 | 0.00 |
| BOX032 | 7 | 1000 | 1111.94 | 975.71 | 72.26 | 74.09 | 28.40 | 0.00 | 0.00 |
| BOX045 | 9 | 1000 | 441.11 | 304.22 | 95.17 | 70.63 | 29.49 | 0.00 | 0.00 |
| BOX047 | 5 | 1000 | 813.98 | 678.00 | 69.01 | 85.05 | 31.61 | 0.00 | 0.00 |
| BOX049 | 18 | 1000 | 234.82 | 97.61 | 70.29 | 69.28 | 28.98 | 0.00 | 0.00 |
| BOX050 | 7 | 1000 | 391.56 | 254.00 | 69.78 | 67.81 | 29.42 | 0.00 | 0.00 |
| BOX054 | 5 | 1000 | 894.51 | 757.20 | 101.98 | 72.65 | 29.60 | 0.00 | 0.00 |
| BOX060 | 7 | 1000 | 341.08 | 206.57 | 71.92 | 88.69 | 33.46 | 0.00 | 0.00 |
| BOX075 | 13 | 1000 | 594.80 | 457.69 | 74.32 | 82.11 | 29.90 | 0.00 | 0.00 |
| BOX076 | 7 | 1000 | 653.14 | 515.57 | 77.07 | 67.93 | 28.55 | 0.00 | 0.00 |
| BOX096 | 5 | 1000 | 805.45 | 669.00 | 74.60 | 104.08 | 26.72 | 0.00 | 0.00 |
| BOX121 | 11 | 1000 | 551.71 | 416.18 | 98.41 | 72.06 | 29.29 | 0.00 | 0.00 |
| BOX122 | 22 | 1000 | 657.03 | 519.36 | 76.41 | 70.72 | 27.03 | 0.00 | 0.00 |
| BOX142 | 5 | 1000 | 464.71 | 328.00 | 68.83 | 74.21 | 26.72 | 0.00 | 0.00 |
| BOX149 | 8 | 1000 | 622.17 | 485.38 | 77.10 | 86.71 | 26.14 | 0.00 | 0.00 |
| BOX154 | 5 | 1000 | 1111.77 | 977.80 | 78.14 | 119.96 | 26.46 | 0.00 | 0.00 |
| BOX175 | 7 | 1000 | 620.49 | 482.57 | 69.21 | 68.68 | 29.25 | 0.00 | 0.00 |
| BOX177 | 6 | 1000 | 648.82 | 512.50 | 72.68 | 66.77 | 28.96 | 0.00 | 0.00 |
| BOX187 | 6 | 1000 | 505.04 | 367.50 | 72.11 | 79.80 | 27.87 | 0.00 | 0.00 |
| BOX192 | 7 | 1000 | 437.02 | 299.00 | 93.54 | 84.95 | 26.88 | 0.00 | 0.00 |
| BOX202 | 8 | 1000 | 526.34 | 391.00 | 82.35 | 120.95 | 31.08 | 0.00 | 0.00 |
| BOX214 | 9 | 1000 | 306.47 | 169.22 | 89.10 | 70.14 | 29.21 | 0.00 | 0.00 |
| BOX240 | 6 | 1000 | 743.78 | 608.67 | 100.12 | 116.02 | 32.53 | 0.00 | 0.00 |
| BOX246 | 8 | 1000 | 792.70 | 649.13 | 107.43 | 118.98 | 29.39 | 0.00 | 0.00 |
| BOX258 | 16 | 1000 | 748.19 | 607.69 | 120.30 | 69.41 | 28.51 | 0.00 | 0.00 |
| BOX259 | 5 | 1000 | 404.47 | 268.60 | 71.92 | 87.62 | 35.57 | 0.00 | 0.00 |
| BOX281 | 6 | 1000 | 633.38 | 490.33 | 78.82 | 131.07 | 28.99 | 0.00 | 0.00 |
| BOX290 | 5 | 1000 | 500.99 | 364.20 | 75.89 | 68.79 | 30.28 | 0.00 | 0.00 |
| Total / Avg. | 228 | 28000 | 613.37 | 476.16 | 82.13 | 84.32 | 26.09% - 35.57% | 0.00 | 0.00 |

Supplementary Table 11: BALiBASE RV913 dataset.

| Domain | Seq. | Tests | Protein len. | Domain len. | Seq. before | Seq. after | Identity (%) | Orig. before | Orig. after |
|---|---|---|---|---|---|---|---|---|---|
| BOX012 | 8 | 1000 | 633.76 | 497.63 | 70.93 | 67.47 | 55.10 | 0.00 | 0.00 |
| BOX017 | 5 | 1000 | 314.89 | 178.60 | 72.21 | 71.13 | 51.67 | 0.00 | 0.00 |
| BOX032 | 5 | 1000 | 1109.33 | 972.00 | 74.57 | 74.11 | 52.70 | 0.00 | 0.00 |
| BOX034 | 10 | 1000 | 527.24 | 389.20 | 69.65 | 70.91 | 50.67 | 0.00 | 0.00 |
| BOX036 | 8 | 1000 | 641.25 | 503.50 | 76.44 | 70.46 | 54.89 | 0.00 | 0.00 |
| BOX043 | 8 | 1000 | 693.72 | 556.13 | 68.85 | 78.20 | 60.19 | 0.00 | 0.00 |
| BOX049 | 9 | 1000 | 237.70 | 98.89 | 68.56 | 70.48 | 52.80 | 0.00 | 0.00 |
| BOX063 | 8 | 1000 | 1512.36 | 1374.13 | 70.63 | 85.26 | 52.83 | 0.00 | 0.00 |
| BOX075 | 7 | 1000 | 579.67 | 443.86 | 69.82 | 72.01 | 61.59 | 0.00 | 0.00 |
| BOX076 | 10 | 1000 | 634.08 | 495.90 | 70.34 | 69.00 | 49.94 | 0.00 | 0.00 |
| BOX079 | 6 | 1000 | 441.23 | 305.00 | 66.99 | 69.24 | 60.84 | 0.00 | 0.00 |
| BOX082 | 7 | 1000 | 758.39 | 621.86 | 68.43 | 68.40 | 57.07 | 0.00 | 0.00 |
| BOX121 | 10 | 1000 | 549.40 | 413.70 | 78.13 | 70.18 | 53.57 | 0.00 | 0.00 |
| BOX122 | 18 | 1000 | 667.88 | 530.33 | 68.72 | 70.19 | 53.13 | 0.00 | 0.00 |
| BOX126 | 6 | 1000 | 429.72 | 292.50 | 67.10 | 70.15 | 57.07 | 0.00 | 0.00 |
| BOX132 | 10 | 1000 | 266.22 | 130.70 | 74.38 | 68.32 | 58.79 | 0.00 | 0.00 |
| BOX133 | 10 | 1000 | 548.77 | 411.30 | 71.53 | 67.98 | 50.44 | 0.00 | 0.00 |
| BOX142 | 6 | 1000 | 469.02 | 332.17 | 67.93 | 70.23 | 65.94 | 0.00 | 0.00 |
| BOX146 | 6 | 1000 | 566.68 | 430.50 | 75.18 | 75.90 | 58.76 | 0.00 | 0.00 |
| BOX158 | 55 | 1000 | 799.15 | 663.64 | 77.82 | 69.48 | 57.92 | 0.00 | 0.00 |
| BOX180 | 6 | 1000 | 342.46 | 205.67 | 71.22 | 67.69 | 52.14 | 0.00 | 0.00 |
| BOX183 | 12 | 1000 | 369.33 | 232.42 | 74.39 | 71.02 | 59.26 | 0.00 | 0.00 |
| BOX214 | 5 | 1000 | 336.57 | 199.20 | 68.07 | 69.29 | 63.19 | 0.00 | 0.00 |
| BOX222 | 7 | 1000 | 1120.51 | 983.14 | 68.33 | 80.30 | 48.34 | 0.00 | 0.00 |
| BOX246 | 5 | 1000 | 740.99 | 603.80 | 108.38 | 122.22 | 61.44 | 0.00 | 0.00 |
| BOX258 | 7 | 1000 | 585.07 | 448.86 | 98.46 | 69.39 | 49.11 | 0.00 | 0.00 |
| BOX290 | 9 | 1000 | 527.03 | 388.67 | 68.69 | 69.14 | 56.37 | 0.00 | 0.00 |
| Total / Avg. | 263 | 27000 | 607.50 | 470.49 | 73.55 | 73.26 | 48.34% - 65.94% | 0.00 | 0.00 |

#### 1.3 GPCR

Supplementary Table 12: GPCR dataset.

| Domain | Seq. | Tests | Protein len. | Domain len. | Seq. before | Seq. after | Identity (%) |
|---|---|---|---|---|---|---|---|
| Alicarboxylic | 12 | 1000 | 399.89 | 258.44 | 70.80 | 70.66 | 42.77 |
| Aminergic | 36 | 1000 | 403.85 | 264.95 | 68.40 | 70.50 | 35.73 |
| Lipid | 36 | 1000 | 398.35 | 255.63 | 72.23 | 70.49 | 19.92 |
| Nucleotide | 12 | 1000 | 402.35 | 263.06 | 68.55 | 70.74 | 25.85 |
| Orphan | 80 | 1000 | 395.04 | 253.77 | 69.60 | 71.68 | 16.64 |
| Peptide | 76 | 1000 | 402.51 | 260.15 | 69.75 | 72.61 | 23.94 |
| Protein | 30 | 1000 | 400.57 | 260.06 | 71.24 | 69.27 | 29.94 |
| Sensory | 8 | 1000 | 400.93 | 262.86 | 67.31 | 70.76 | 32.92 |
| Total / Avg. | 290 | 8000 | 400.44 | 259.87 | 69.73 | 70.84 | 16.64% - 42.77% |

#### 2 Model comparison

Supplementary Table 13: Model comparison on CDD natural dataset.

| Domain | Ankh | ESM1b | ESM2 | ProtAlbert | ProtBert | ProtT5 | ProtXLNet |
|---|---|---|---|---|---|---|---|
| **d_cc** | | | | | | | |
| cd00012 | 0.0446 | 0.0741 | 0.0670 | 0.0754 | 0.0819 | **0.0441** | 0.0863 |
| cd00083 | **0.0026** | 0.2820 | 0.2325 | 0.1945 | 0.2684 | 0.0063 | 0.2808 |
| cd00173 | **0.0335** | 0.3763 | 0.3248 | 0.3285 | 0.3565 | 0.0346 | 0.3912 |
| cd00637 | 0.0044 | 0.0285 | 0.0052 | 0.0060 | 0.0190 | **0.0041** | 0.0248 |
| cd01040 | **0.0037** | 0.0558 | 0.0155 | 0.0279 | 0.0299 | 0.0052 | 0.0844 |
| cd01068 | **0.0027** | 0.0493 | 0.0036 | 0.0274 | 0.0490 | 0.0028 | 0.0854 |
| cd10140 | **0.0596** | 0.1005 | 0.0808 | 0.1051 | 0.1022 | 0.0611 | 0.1421 |
| cd21116 | **0.0084** | 0.0627 | 0.0392 | 0.0416 | 0.0551 | 0.0101 | 0.0565 |
| **d_d** | | | | | | | |
| cd00012 | **0.0235** | 0.0480 | 0.0416 | 0.0507 | 0.0616 | 0.0242 | 0.0697 |
| cd00083 | **0.0006** | 0.2558 | 0.2223 | 0.1856 | 0.2513 | 0.0020 | 0.2582 |
| cd00173 | **0.0081** | 0.3564 | 0.3071 | 0.3154 | 0.3332 | 0.0082 | 0.3688 |
| cd00637 | 0.0010 | 0.0063 | 0.0010 | 0.0011 | 0.0057 | **0.0009** | 0.0051 |
| cd01040 | **0.0007** | 0.0108 | 0.0023 | 0.0043 | 0.0073 | 0.0010 | 0.0152 |
| cd01068 | **0.0006** | 0.0046 | 0.0006 | 0.0022 | 0.0027 | 0.0007 | 0.0231 |
| cd10140 | **0.0120** | 0.0585 | 0.0429 | 0.0744 | 0.0648 | 0.0126 | 0.1204 |
| cd21116 | **0.0029** | 0.0262 | 0.0139 | 0.0188 | 0.0229 | 0.0037 | 0.0239 |
| **d_pos** | | | | | | | |
| cd00012 | **0.6738** | 0.9292 | 0.8208 | 0.9102 | 0.9345 | 0.6956 | 0.9483 |
| cd00083 | **0.0430** | 0.6975 | 0.5199 | 0.4463 | 0.6206 | 0.0869 | 0.6642 |
| cd00173 | 0.4884 | 1.0000 | 0.8062 | 0.7908 | 0.9638 | **0.4589** | 1.0000 |
| cd00637 | **0.1315** | 0.4186 | 0.1510 | 0.1708 | 0.3103 | 0.1316 | 0.4549 |
| cd01040 | **0.1006** | 0.5644 | 0.2028 | 0.3654 | 0.3835 | 0.1208 | 0.7838 |
| cd01068 | **0.1046** | 0.4887 | 0.1144 | 0.3384 | 0.4442 | 0.1093 | 0.9463 |
| cd10140 | **0.7260** | 0.9539 | 0.7884 | 0.9099 | 0.9016 | 0.7372 | 1.0000 |
| cd21116 | **0.2944** | 0.8148 | 0.5874 | 0.6168 | 0.7046 | 0.3255 | 0.7975 |
| **d_seq** | | | | | | | |
| cd00012 | **0.5926** | 0.8576 | 0.7631 | 0.8548 | 0.8521 | 0.6100 | 0.8653 |
| cd00083 | **0.0333** | 0.6750 | 0.4963 | 0.4243 | 0.5976 | 0.0765 | 0.6424 |
| cd00173 | 0.4576 | 0.9168 | 0.7683 | 0.7399 | 0.9141 | **0.4310** | 0.9253 |
| cd00637 | **0.1055** | 0.4137 | 0.1246 | 0.1416 | 0.2980 | 0.1083 | 0.4417 |
| cd01040 | **0.0795** | 0.5599 | 0.1914 | 0.3552 | 0.3760 | 0.1005 | 0.7792 |
| cd01068 | **0.1028** | 0.4884 | 0.1131 | 0.3370 | 0.4437 | 0.1068 | 0.9457 |
| cd10140 | **0.6739** | 0.8731 | 0.7472 | 0.8581 | 0.8619 | 0.6907 | 0.9430 |
| cd21116 | **0.2594** | 0.8066 | 0.5672 | 0.5988 | 0.6944 | 0.2846 | 0.7827 |
| **d_ssp** | | | | | | | |
| cd00012 | **0.7964** | 0.9623 | 0.9038 | 0.9528 | 0.9658 | 0.8139 | 0.9725 |
| cd00083 | **0.0885** | 0.8689 | 0.5588 | 0.4851 | 0.6624 | 0.1559 | 0.8137 |
| cd00173 | 0.5958 | 1.0000 | 0.8966 | 0.8708 | 0.9850 | **0.5767** | 1.0000 |
| cd00637 | **0.2211** | 0.5842 | 0.2466 | 0.2758 | 0.4763 | 0.2239 | 0.6224 |
| cd01040 | **0.1741** | 0.6764 | 0.3619 | 0.4799 | 0.4976 | 0.2036 | 0.8766 |
| cd01068 | **0.1720** | 0.5727 | 0.2000 | 0.4358 | 0.5311 | 0.1778 | 0.9806 |
| cd10140 | **0.8359** | 0.9777 | 0.8724 | 0.9541 | 0.9447 | 0.8435 | 1.0000 |
| cd21116 | **0.4129** | 0.8920 | 0.6838 | 0.7149 | 0.8052 | 0.4562 | 0.8805 |

### 3 Missed alignments

Supplementary Table 14: Percentage of missed alignments for all methods on the natural datasets

| Domain | Ankh | ProtT5 | DEDAL | pLM-BLAST | | | BLOSUMs | | | | |
|---|---|---|---|---|---|---|---|---|---|---|---|
| | | | | min 100 | min 60 | min 20 | 45 | 50 | 62 | 80 | 90 |
| CDD | | | | | | | | | | | |
| cd00012 | 6.06 | 6.02 | 53.85 | 91.58 | 81.96 | 92.02 | 39.90 | 44.99 | 80.62 | 87.58 | 88.92 |
| cd00083 | 8.33 | 3.91 | 51.12 | 84.32 | 42.02 | 82.12 | 26.21 | 28.62 | 29.19 | 41.61 | 44.32 |
| cd00173 | 27.06 | 20.32 | 84.89 | 68.89 | 51.05 | 89.67 | 61.26 | 60.55 | 77.52 | 89.21 | 89.69 |
| cd00637 | 0.02 | 0.01 | 2.11 | 54.89 | 65.59 | 91.72 | 2.90 | 3.27 | 11.37 | 35.07 | 41.91 |
| cd01040 | 0.07 | 0.07 | 5.64 | 66.28 | 46.41 | 80.05 | 10.63 | 13.45 | 38.00 | 61.04 | 65.65 |
| cd01068 | 0.48 | 0.48 | 20.88 | 28.43 | 40.16 | 88.76 | 15.32 | 22.38 | 38.79 | 63.47 | 67.37 |
| cd10140 | 5.64 | 5.75 | 44.80 | 67.92 | 63.14 | 87.88 | 24.29 | 25.91 | 43.85 | 59.42 | 63.05 |
| cd21116 | 1.43 | 1.03 | 22.79 | 80.66 | 78.61 | 92.36 | 12.01 | 13.29 | 37.94 | 63.52 | 65.15 |
| Average | 6.14 | 4.70 | 35.76 | 67.87 | 58.62 | 88.07 | 24.06 | 26.56 | 44.66 | 62.61 | 65.76 |
| BALiBASE RV11 | | | | | | | | | | | |
| BBS11009 | 17.12 | 17.12 | 14.53 | 100.00 | 67.53 | 91.15 | 26.46 | 27.95 | 33.61 | 48.60 | 52.84 |
| BBS11010 | 3.25 | 3.89 | 34.10 | 86.78 | 81.67 | 97.26 | 17.56 | 39.23 | 58.85 | 78.65 | 74.09 |
| BBS11016 | 0.18 | 0.20 | 22.86 | 71.59 | 78.57 | 91.81 | 9.22 | 7.02 | 54.53 | 69.47 | 73.70 |
| BBS11018 | 0.60 | 0.64 | 27.57 | 81.37 | 82.61 | 95.03 | 7.66 | 7.34 | 52.96 | 73.02 | 74.48 |
| BBS11024 | 2.23 | 2.26 | 78.98 | 92.63 | 75.58 | 94.31 | 11.93 | 13.82 | 80.15 | 85.85 | 87.63 |
| BBS11026 | 11.55 | 14.20 | 51.00 | 100.00 | 81.79 | 56.52 | 60.67 | 58.78 | 67.27 | 75.42 | 76.01 |
| BBS11034 | 0.25 | 0.23 | 24.10 | 78.42 | 78.63 | 92.65 | 8.91 | 8.22 | 53.13 | 65.44 | 72.03 |
| BBS11037 | 0.95 | 0.89 | 44.35 | 65.91 | 69.18 | 90.92 | 10.37 | 11.25 | 59.65 | 84.57 | 87.15 |
| BBS11038 | 0.35 | 0.32 | 8.77 | 68.91 | 67.73 | 88.10 | 9.52 | 10.99 | 24.53 | 41.07 | 44.62 |
| Average | 4.05 | 4.42 | 34.03 | 82.85 | 75.92 | 88.64 | 18.03 | 20.51 | 53.85 | 69.12 | 71.39 |
| BALiBASE RV12 | | | | | | | | | | | |
| BBS12008 | 0.08 | 0.09 | 5.95 | 50.90 | 61.41 | 90.12 | 1.72 | 2.43 | 3.71 | 6.84 | 10.78 |
| BBS12011 | 0.16 | 0.16 | 61.72 | 78.33 | 90.79 | 96.91 | 0.90 | 0.98 | 1.54 | 5.23 | 5.04 |
| BBS12030 | 0.24 | 0.24 | 64.41 | 52.08 | 67.57 | 98.71 | 3.88 | 3.16 | 4.69 | 7.63 | 7.12 |
| BBS12033 | 0.83 | 0.93 | 11.05 | 64.41 | 69.83 | 91.53 | 2.98 | 3.72 | 11.00 | 22.95 | 23.18 |
| BBS12037 | 0.10 | 0.18 | 23.87 | 58.87 | 69.99 | 92.70 | 1.96 | 1.88 | 8.20 | 28.00 | 31.47 |
| Average | 0.28 | 0.32 | 33.40 | 60.92 | 71.92 | 93.99 | 2.29 | 2.43 | 5.83 | 14.13 | 15.52 |
| BALiBASE RV30 | | | | | | | | | | | |
| BBS30001 | 0.80 | 1.19 | 18.14 | 49.52 | 67.58 | 96.95 | 2.58 | 2.94 | 21.17 | 36.40 | 39.11 |
| BBS30004 | 0.25 | 0.22 | 23.04 | 54.96 | 62.75 | 92.02 | 3.48 | 4.51 | 16.70 | 26.41 | 27.74 |
| BBS30005 | 0.35 | 0.29 | 5.42 | 77.59 | 86.75 | 97.33 | 2.52 | 3.02 | 11.40 | 23.26 | 26.49 |
| BBS30006 | 10.62 | 11.38 | 51.95 | 91.40 | 61.42 | 60.02 | 19.58 | 20.93 | 35.93 | 45.08 | 46.92 |
| BBS30008 | 2.79 | 2.89 | 39.03 | 76.08 | 72.33 | 95.06 | 10.57 | 9.54 | 32.63 | 41.98 | 42.72 |
| BBS30009 | 17.86 | 6.43 | 14.62 | 95.10 | 68.17 | 66.52 | 29.02 | 29.56 | 38.80 | 44.88 | 45.57 |
| BBS30010 | 0.09 | 0.04 | 32.53 | 46.06 | 64.75 | 94.62 | 1.37 | 1.53 | 6.93 | 17.47 | 19.09 |
| BBS30013 | 0.23 | 0.15 | 45.71 | 78.62 | 81.84 | 94.91 | 3.99 | 5.20 | 38.69 | 50.30 | 50.85 |
| BBS30016 | 35.44 | 32.69 | 23.63 | 80.44 | 72.29 | 80.80 | 50.83 | 49.08 | 57.14 | 58.83 | 59.29 |
| BBS30030 | 0.18 | 0.18 | 21.09 | 58.44 | 61.30 | 88.77 | 4.55 | 5.26 | 14.58 | 34.73 | 38.26 |
| Average | 6.86 | 5.55 | 27.51 | 70.82 | 69.92 | 86.70 | 12.85 | 13.16 | 27.40 | 37.93 | 39.60 |
| Average all | 4.86 | 4.20 | 32.33 | 71.92 | 69.09 | 88.73 | 15.46 | 16.90 | 35.78 | 49.16 | 51.32 |

### 4 Comparison against BLOSUM

#### 4.1 CDD natural

Supplementary Table 15: Ankh-score vs best BLOSUM on CDD natural dataset.

| Domain | $d_{cc}$ | | | $d_d$ | | | $d_{pos}$ | | | $d_{seq}$ | | | $d_{ssp}$ | | |
|---|---|---|---|---|---|---|---|---|---|---|---|---|---|---|---|
| Average distances | | | | | | | | | | | | | | | |
| | Ankh | BLOSUM | P-value | Ankh | BLOSUM | P-value | Ankh | BLOSUM | P-value | Ankh | BLOSUM | P-value | Ankh | BLOSUM | P-value |
| cd00012 | **0.0446** | 0.1472 | 9.3E-275 | **0.0235** | 0.1445 | 2.2E-262 | **0.6738** | 0.9617 | 3.5E-285 | **0.5926** | 0.7604 | 1.1E-169 | **0.7964** | 0.9802 | 8.6E-285 |
| cd00083 | **0.0026** | 0.0356 | 2.1E-131 | **0.0006** | 0.0141 | 1.6E-128 | **0.0430** | 0.3998 | 8.5E-133 | **0.0333** | 0.3813 | 1.5E-133 | **0.0885** | 0.4771 | 6.2E-135 |
| cd00173 | **0.0335** | 0.3052 | 5.1E-31 | **0.0081** | 0.2873 | 2.5E-29 | **0.4884** | 0.7891 | 7.2E-34 | **0.4576** | 0.7385 | 3.8E-34 | **0.5958** | 0.8658 | 1.5E-34 |
| cd00637 | **0.0044** | 0.0179 | 0.0E+00 | **0.0010** | 0.0053 | 0.0E+00 | **0.1315** | 0.4094 | 0.0E+00 | **0.1055** | 0.3880 | 0.0E+00 | **0.2211** | 0.5701 | 0.0E+00 |
| cd01040 | **0.0037** | 0.0338 | 1.8E-298 | **0.0007** | 0.0099 | 7.6E-303 | **0.1006** | 0.4217 | 4.9E-302 | **0.0795** | 0.4078 | 3.4E-303 | **0.1741** | 0.5481 | 1.3E-303 |
| cd01068 | **0.0027** | 0.0334 | 2.6E-27 | **0.0006** | 0.0079 | 8.5E-29 | **0.1046** | 0.4230 | 6.9E-29 | **0.1028** | 0.4199 | 6.7E-29 | **0.1720** | 0.5325 | 6.3E-29 |
| cd10140 | **0.0596** | 0.0859 | 3.0E-85 | **0.0120** | 0.0367 | 2.7E-90 | **0.7260** | 0.8695 | 2.6E-55 | **0.6739** | 0.7825 | 7.8E-21 | **0.8359** | 0.9297 | 2.5E-54 |
| cd21116 | **0.0084** | 0.0456 | 1.7E-51 | **0.0029** | 0.0231 | 1.3E-54 | **0.2944** | 0.6259 | 8.8E-55 | **0.2594** | 0.6012 | 1.7E-54 | **0.4129** | 0.7563 | 9.1E-55 |
| Domains won | 8 | 0 | | 8 | 0 | | 8 | 0 | | 8 | 0 | | 8 | 0 | |
| Tests won | | | | | | | | | | | | | | | |
| | Ankh | BLOSUM | Equal | Ankh | BLOSUM | Equal | Ankh | BLOSUM | Equal | Ankh | BLOSUM | Equal | Ankh | BLOSUM | Equal |
| cd00012 | **1699** | 70 | 1 | **1639** | 130 | 1 | **1723** | 17 | 30 | **1396** | 369 | 5 | **1723** | 15 | 32 |
| cd00083 | **870** | 45 | 166 | **864** | 40 | 177 | **866** | 32 | 183 | **866** | 35 | 180 | **866** | 35 | 180 |
| cd00173 | **226** | 30 | 69 | **225** | 31 | 69 | **230** | 22 | 73 | **231** | 22 | 72 | **232** | 21 | 72 |
| cd00637 | **2996** | 7 | 0 | **2960** | 43 | 0 | **3002** | 1 | 0 | **2999** | 3 | 1 | **2999** | 3 | 1 |
| cd01040 | **1813** | 58 | 20 | **1821** | 45 | 25 | **1814** | 41 | 36 | **1823** | 39 | 29 | **1824** | 40 | 27 |
| cd01068 | **162** | 4 | 5 | **164** | 2 | 5 | **165** | 1 | 5 | **165** | 1 | 5 | **165** | 1 | 5 |
| cd10140 | **541** | 86 | 3 | **565** | 62 | 3 | **451** | 101 | 78 | **364** | 253 | 13 | **445** | 116 | 69 |
| cd21116 | **313** | 12 | 0 | **319** | 6 | 0 | **320** | 5 | 0 | **317** | 8 | 0 | **319** | 6 | 0 |
| Total | **8620** | 312 | 264 | **8557** | 359 | 280 | **8571** | 220 | 405 | **8161** | 730 | 305 | **8573** | 237 | 386 |
| Tests won (%) | | | | | | | | | | | | | | | |
| cd00012 | **95.99** | 3.95 | 0.06 | **92.60** | 7.34 | 0.06 | **97.34** | 0.96 | 1.69 | **78.87** | 20.85 | 0.28 | **97.34** | 0.85 | 1.81 |
| cd00083 | **80.48** | 4.16 | 15.36 | **79.93** | 3.70 | 16.37 | **80.11** | 2.96 | 16.93 | **80.11** | 3.24 | 16.65 | **80.11** | 3.24 | 16.65 |
| cd00173 | **69.54** | 9.23 | 21.23 | **69.23** | 9.54 | 21.23 | **70.77** | 6.77 | 22.46 | **71.08** | 6.77 | 22.15 | **71.38** | 6.46 | 22.15 |
| cd00637 | **99.77** | 0.23 | 0.00 | **98.57** | 1.43 | 0.00 | **99.97** | 0.03 | 0.00 | **99.87** | 0.10 | 0.03 | **99.87** | 0.10 | 0.03 |
| cd01040 | **95.88** | 3.07 | 1.06 | **96.30** | 2.38 | 1.32 | **95.93** | 2.17 | 1.90 | **96.40** | 2.06 | 1.53 | **96.46** | 2.12 | 1.43 |
| cd01068 | **94.74** | 2.34 | 2.92 | **95.91** | 1.17 | 2.92 | **96.49** | 0.58 | 2.92 | **96.49** | 0.58 | 2.92 | **96.49** | 0.58 | 2.92 |
| cd10140 | **85.87** | 13.65 | 0.48 | **89.68** | 9.84 | 0.48 | **71.59** | 16.03 | 12.38 | **57.78** | 40.16 | 2.06 | **70.63** | 18.41 | 10.95 |
| cd21116 | **96.31** | 3.69 | 0.00 | **98.15** | 1.85 | 0.00 | **98.46** | 1.54 | 0.00 | **97.54** | 2.46 | 0.00 | **98.15** | 1.85 | 0.00 |
| Average | **89.82** | 5.04 | 5.14 | **90.05** | 4.66 | 5.30 | **88.83** | 3.88 | 7.29 | **84.77** | 9.53 | 5.71 | **88.81** | 4.20 | 6.99 |

Supplementary Table 16: Best BLOSUM matrices for the previous table.

| Domain | $d_{cc}$ | $d_d$ | $d_{pos}$ | $d_{seq}$ | $d_{ssp}$ |
|---|---|---|---|---|---|
| cd00012 | 45 | 45 | 45 | 62 | 45 |
| cd00083 | 45 | 45 | 45 | 45 | 45 |
| cd00173 | 50 | 50 | 50 | 50 | 50 |
| cd00637 | 45 | 62 | 45 | 62 | 62 |
| cd01040 | 45 | 45 | 45 | 45 | 45 |
| cd01068 | 45 | 45 | 45 | 45 | 45 |
| cd10140 | 45 | 45 | 45 | 62 | 45 |
| cd21116 | 45 | 45 | 45 | 45 | 45 |
| BLOSUM | 45 | 50 | 62 | 80 | 90 |
| Total | 30 | 5 | 5 | 0 | 0 |

#### 4.2 BALiBASE RV11

Supplementary Table 17: Ankh-score vs best BLOSUM on BALiBASE RV11 dataset.

| Domain | $d_{cc}$ | | | $d_d$ | | | $d_{pos}$ | | | $d_{seq}$ | | | $d_{ssp}$ | | |
|---|---|---|---|---|---|---|---|---|---|---|---|---|---|---|---|
| Average distances | | | | | | | | | | | | | | | |
| | Ankh | BLOSUM | P-value | Ankh | BLOSUM | P-value | Ankh | BLOSUM | P-value | Ankh | BLOSUM | P-value | Ankh | BLOSUM | P-value |
| BBS11009 | 0.0207 | 0.0358 | 4.3E-02 | 0.0103 | 0.0164 | 8.0E-02 | 0.2317 | 0.3901 | 4.3E-02 | 0.2134 | 0.3707 | 4.3E-02 | 0.3654 | 0.6148 | 4.3E-02 |
| BBS11010 | 0.0232 | 0.0630 | 3.1E-02 | 0.0039 | 0.0291 | 3.1E-02 | 0.2844 | 0.6431 | 3.1E-02 | 0.2093 | 0.5842 | 3.1E-02 | 0.4258 | 0.7493 | 3.1E-02 |
| BBS11016 | **0.0111** | 0.0307 | 1.5E-08 | **0.0041** | 0.0115 | 1.5E-08 | **0.3578** | 0.6084 | 7.5E-09 | **0.3195** | 0.5951 | 7.5E-09 | **0.5197** | 0.7678 | 7.5E-09 |
| BBS11018 | **0.0190** | 0.0408 | 1.2E-16 | **0.0096** | 0.0150 | 1.1E-06 | **0.4264** | 0.6538 | 1.2E-16 | **0.3623** | 0.6039 | 1.2E-16 | **0.5856** | 0.7803 | 1.2E-16 |
| BBS11024 | 0.0244 | 0.0457 | 3.1E-02 | 0.0069 | 0.0278 | 6.3E-02 | 0.4017 | 0.7209 | 3.1E-02 | 0.3226 | 0.7182 | 3.1E-02 | 0.5443 | 0.8557 | 3.1E-02 |
| BBS11026 | **0.0873** | 0.2578 | 7.2E-04 | **0.0298** | 0.2296 | 3.5E-04 | **0.6577** | 0.8642 | 6.8E-04 | **0.6351** | 0.7692 | 3.5E-04 | **0.7841** | 0.9151 | 2.0E-04 |
| BBS11034 | **0.0196** | 0.0349 | 1.5E-08 | **0.0061** | 0.0150 | 5.7E-06 | **0.3908** | 0.6368 | 7.5E-09 | **0.3499** | 0.6059 | 7.5E-09 | **0.5500** | 0.7863 | 7.5E-09 |
| BBS11037 | **0.0229** | 0.0426 | 2.0E-03 | **0.0043** | 0.0151 | 2.0E-03 | **0.4064** | 0.7201 | 2.0E-03 | **0.3575** | 0.7011 | 2.0E-03 | **0.5548** | 0.8308 | 2.0E-03 |
| BBS11038 | **0.0167** | 0.0351 | 7.5E-09 | **0.0032** | 0.0069 | 5.7E-06 | **0.2270** | 0.4361 | 7.5E-09 | **0.1905** | 0.4053 | 7.5E-09 | **0.3564** | 0.5910 | 7.5E-09 |
| Domains won | **6** | 0 | | **6** | 0 | | **6** | 0 | | **6** | 0 | | **6** | 0 | |
| Tests won | | | | | | | | | | | | | | | |
| | Ankh | BLOSUM | Equal | Ankh | BLOSUM | Equal | Ankh | BLOSUM | Equal | Ankh | BLOSUM | Equal | Ankh | BLOSUM | Equal |
| BBS11009 | 5 | 0 | 1 | 4 | 1 | 1 | 5 | 0 | 1 | 5 | 0 | 1 | 5 | 0 | 1 |
| BBS11010 | 6 | 0 | 0 | 6 | 0 | 0 | 6 | 0 | 0 | 6 | 0 | 0 | 6 | 0 | 0 |
| BBS11016 | **27** | 1 | 0 | **27** | 1 | 0 | **28** | 0 | 0 | **28** | 0 | 0 | **28** | 0 | 0 |
| BBS11018 | **90** | 1 | 0 | **58** | 33 | 0 | **91** | 0 | 0 | **91** | 0 | 0 | **91** | 0 | 0 |
| BBS11024 | 6 | 0 | 0 | 5 | 1 | 0 | 6 | 0 | 0 | 6 | 0 | 0 | 6 | 0 | 0 |
| BBS11026 | **18** | 3 | 0 | **18** | 3 | 0 | **19** | 1 | 1 | **19** | 2 | 0 | **20** | 1 | 0 |
| BBS11034 | **27** | 1 | 0 | **24** | 4 | 0 | **28** | 0 | 0 | **28** | 0 | 0 | **28** | 0 | 0 |
| BBS11037 | **10** | 0 | 0 | **10** | 0 | 0 | **10** | 0 | 0 | **10** | 0 | 0 | **10** | 0 | 0 |
| BBS11038 | **28** | 0 | 0 | **23** | 5 | 0 | **28** | 0 | 0 | **28** | 0 | 0 | **28** | 0 | 0 |
| Total | **200** | 6 | 0 | **160** | 46 | 0 | **204** | 1 | 1 | **204** | 2 | 0 | **205** | 1 | 0 |
| Tests won (%) | | | | | | | | | | | | | | | |
| BBS11009 | 83.33 | 0.00 | 16.67 | 66.67 | 16.67 | 16.67 | 83.33 | 0.00 | 16.67 | 83.33 | 0.00 | 16.67 | 83.33 | 0.00 | 16.67 |
| BBS11010 | 100.00 | 0.00 | 0.00 | 100.00 | 0.00 | 0.00 | 100.00 | 0.00 | 0.00 | 100.00 | 0.00 | 0.00 | 100.00 | 0.00 | 0.00 |
| BBS11016 | **96.43** | 3.57 | 0.00 | **96.43** | 3.57 | 0.00 | **100.00** | 0.00 | 0.00 | **100.00** | 0.00 | 0.00 | **100.00** | 0.00 | 0.00 |
| BBS11018 | **98.90** | 1.10 | 0.00 | **63.74** | 36.26 | 0.00 | **100.00** | 0.00 | 0.00 | **100.00** | 0.00 | 0.00 | **100.00** | 0.00 | 0.00 |
| BBS11024 | 100.00 | 0.00 | 0.00 | 83.33 | 16.67 | 0.00 | 100.00 | 0.00 | 0.00 | 100.00 | 0.00 | 0.00 | 100.00 | 0.00 | 0.00 |
| BBS11026 | **85.71** | 14.29 | 0.00 | **85.71** | 14.29 | 0.00 | **90.48** | 4.76 | 4.76 | **90.48** | 9.52 | 0.00 | **95.24** | 4.76 | 0.00 |
| BBS11034 | **96.43** | 3.57 | 0.00 | **85.71** | 14.29 | 0.00 | **100.00** | 0.00 | 0.00 | **100.00** | 0.00 | 0.00 | **100.00** | 0.00 | 0.00 |
| BBS11037 | **100.00** | 0.00 | 0.00 | **100.00** | 0.00 | 0.00 | **100.00** | 0.00 | 0.00 | **100.00** | 0.00 | 0.00 | **100.00** | 0.00 | 0.00 |
| BBS11038 | **100.00** | 0.00 | 0.00 | **82.14** | 17.86 | 0.00 | **100.00** | 0.00 | 0.00 | **100.00** | 0.00 | 0.00 | **100.00** | 0.00 | 0.00 |
| Average | **96.25** | 3.75 | 0.00 | **85.62** | 14.38 | 0.00 | **98.41** | 0.79 | 0.79 | **98.41** | 1.59 | 0.00 | **99.21** | 0.79 | 0.00 |

Supplementary Table 18: Best BLOSUM matrices for the previous table.

| Domain | $d_{cc}$ | $d_d$ | $d_{pos}$ | $d_{seq}$ | $d_{ssp}$ |
|---|---|---|---|---|---|
| BBS11009 | 45 | 45 | 45 | 45 | 50 |
| BBS11010 | 45 | 45 | 45 | 50 | 45 |
| BBS11016 | 45 | 50 | 50 | 50 | 50 |
| BBS11018 | 50 | 50 | 45 | 45 | 45 |
| BBS11024 | 50 | 45 | 45 | 45 | 45 |
| BBS11026 | 50 | 50 | 50 | 50 | 50 |
| BBS11034 | 45 | 45 | 50 | 50 | 50 |
| BBS11037 | 50 | 45 | 45 | 45 | 45 |
| BBS11038 | 45 | 45 | 45 | 45 | 45 |
| BLOSUM | 45 | 50 | 62 | 80 | 90 |
| Total | 27 | 18 | 0 | 0 | 0 |

#### 4.3 BALiBASE RV12

Supplementary Table 19: Ankh-score vs best BLOSUM on BALiBASE RV12 dataset.

| Domain | $d_{cc}$ | | | $d_d$ | | | $d_{pos}$ | | | $d_{seq}$ | | | $d_{ssp}$ | | |
|---|---|---|---|---|---|---|---|---|---|---|---|---|---|---|---|
| Average distances | | | | | | | | | | | | | | | |
| | Ankh | BLOSUM | P-value | Ankh | BLOSUM | P-value | Ankh | BLOSUM | P-value | Ankh | BLOSUM | P-value | Ankh | BLOSUM | P-value |
| BBS11009 | **0.0011** | 0.0051 | 2.8E-14 | **0.0003** | 0.0015 | 9.7E-13 | **0.0674** | 0.1519 | 9.2E-14 | **0.0635** | 0.1498 | 8.1E-14 | **0.1263** | 0.2590 | 6.0E-14 |
| BBS11026 | **0.0068** | 0.0084 | 1.0E-08 | 0.0031 | 0.0036 | 1.9E-01 | **0.3188** | 0.3759 | 7.3E-10 | **0.3062** | 0.3672 | 7.8E-10 | **0.4780** | 0.5517 | 8.8E-11 |
| BBS11034 | **0.0018** | 0.0075 | 6.1E-04 | **0.0002** | 0.0023 | 6.1E-04 | **0.0721** | 0.1344 | 3.4E-03 | **0.0701** | 0.1320 | 2.6E-03 | **0.1412** | 0.2297 | 2.6E-03 |
| BBS11037 | **0.0129** | 0.0239 | 1.9E-06 | **0.0032** | 0.0066 | 9.5E-06 | **0.2973** | 0.4267 | 4.8E-06 | **0.2360** | 0.3886 | 1.3E-05 | **0.4392** | 0.6003 | 4.8E-06 |
| BBS11038 | **0.0038** | 0.0091 | 1.7E-14 | **0.0022** | 0.0037 | 3.9E-08 | **0.1880** | 0.3437 | 1.7E-14 | **0.1726** | 0.3302 | 1.7E-14 | **0.3136** | 0.5088 | 1.7E-14 |
| Domains won | **5** | 0 | | **4** | 0 | | **5** | 0 | | **5** | 0 | | **5** | 0 | |
| Tests won | | | | | | | | | | | | | | | |
| | Ankh | BLOSUM | Equal | Ankh | BLOSUM | Equal | Ankh | BLOSUM | Equal | Ankh | BLOSUM | Equal | Ankh | BLOSUM | Equal |
| BBS11009 | **77** | 1 | 0 | **72** | 6 | 0 | **73** | 4 | 1 | **73** | 5 | 0 | **74** | 4 | 0 |
| BBS11026 | **54** | 12 | 0 | 33 | 33 | 0 | **56** | 9 | 1 | **56** | 10 | 0 | **57** | 9 | 0 |
| BBS11034 | **13** | 2 | 0 | **13** | 2 | 0 | **12** | 3 | 0 | **12** | 3 | 0 | **12** | 3 | 0 |
| BBS11037 | **20** | 1 | 0 | **19** | 2 | 0 | **20** | 1 | 0 | **20** | 1 | 0 | **20** | 1 | 0 |
| BBS11038 | **78** | 0 | 0 | **59** | 19 | 0 | **78** | 0 | 0 | **78** | 0 | 0 | **78** | 0 | 0 |
| Total | **242** | 16 | 0 | **163** | 29 | 0 | **239** | 17 | 2 | **239** | 19 | 0 | **241** | 17 | 0 |
| Tests won (%) | | | | | | | | | | | | | | | |
| BBS11009 | **98.72** | 1.28 | 0.00 | **92.31** | 7.69 | 0.00 | **93.59** | 5.13 | 1.28 | **93.59** | 6.41 | 0.00 | **94.87** | 5.13 | 0.00 |
| BBS11026 | **81.82** | 18.18 | 0.00 | 50.00 | 50.00 | 0.00 | **84.85** | 13.64 | 1.52 | **84.85** | 15.15 | 0.00 | **86.36** | 13.64 | 0.00 |
| BBS11034 | **86.67** | 13.33 | 0.00 | **86.67** | 13.33 | 0.00 | **80.00** | 20.00 | 0.00 | **80.00** | 20.00 | 0.00 | **80.00** | 20.00 | 0.00 |
| BBS11037 | **95.24** | 4.76 | 0.00 | **90.48** | 9.52 | 0.00 | **95.24** | 4.76 | 0.00 | **95.24** | 4.76 | 0.00 | **95.24** | 4.76 | 0.00 |
| BBS11038 | **100.00** | 0.00 | 0.00 | **75.64** | 24.36 | 0.00 | **100.00** | 0.00 | 0.00 | **100.00** | 0.00 | 0.00 | **100.00** | 0.00 | 0.00 |
| Average | **92.49** | 7.51 | 0.00 | **86.27** | 13.73 | 0.00 | **90.74** | 8.71 | 0.56 | **90.74** | 9.26 | 0.00 | **91.29** | 8.71 | 0.00 |

Supplementary Table 20: Best BLOSUM matrices for the previous table.

| Domain | $d_{cc}$ | $d_d$ | $d_{pos}$ | $d_{seq}$ | $d_{ssp}$ |
|---|---|---|---|---|---|
| BBS11009 | 45 | 45 | 62 | 62 | 62 |
| BBS11026 | 62 | 62 | 62 | 62 | 50 |
| BBS11034 | 62 | 62 | 62 | 62 | 62 |
| BBS11037 | 45 | 45 | 45 | 62 | 45 |
| BBS11038 | 62 | 45 | 45 | 45 | 45 |
| BLOSUM | 45 | 50 | 62 | 80 | 90 |
| Total | 10 | 1 | 14 | 0 | 0 |

#### 4.4 BALiBASE RV30

Supplementary Table 21: Ankh-score vs best BLOSUM on BALiBASE RV30 dataset.

| Domain | $d_{cc}$ | | | $d_d$ | | | $d_{pos}$ | | | $d_{seq}$ | | | $d_{ssp}$ | | |
|---|---|---|---|---|---|---|---|---|---|---|---|---|---|---|---|
| Average distances | | | | | | | | | | | | | | | |
| | Ankh | BLOSUM | P-value | Ankh | BLOSUM | P-value | Ankh | BLOSUM | P-value | Ankh | BLOSUM | P-value | Ankh | BLOSUM | P-value |
| BBS30001 | **0.0157** | 0.0165 | 8.9E-101 | **0.0010** | 0.0025 | 3.7E-109 | **0.1513** | 0.2649 | 5.5E-117 | **0.1453** | 0.2605 | 4.2E-118 | **0.2360** | 0.3515 | 1.1E-117 |
| BBS30004 | **0.0019** | 0.0068 | 5.4E-157 | **0.0005** | 0.0021 | 5.4E-150 | **0.1171** | 0.2338 | 2.4E-154 | **0.1042** | 0.2237 | 6.1E-155 | **0.1935** | 0.3293 | 3.0E-155 |
| BBS30005 | **0.0078** | 0.0172 | 1.7E-139 | **0.0005** | 0.0031 | 1.1E-144 | **0.0817** | 0.2439 | 2.2E-147 | **0.0605** | 0.2277 | 2.2E-147 | **0.1422** | 0.3546 | 2.3E-147 |
| BBS30006 | **0.0566** | 0.0744 | 3.4E-10 | **0.0224** | 0.0292 | 8.7E-11 | **0.3454** | 0.4582 | 7.7E-22 | **0.3159** | 0.4120 | 7.2E-21 | **0.4445** | 0.5643 | 8.3E-22 |
| BBS30008 | **0.0200** | 0.0313 | 1.0E-83 | **0.0086** | 0.0134 | 6.6E-34 | **0.3952** | 0.4797 | 2.8E-94 | **0.3591** | 0.4188 | 2.0E-76 | **0.5062** | 0.5708 | 1.0E-95 |
| BBS30009 | **0.0694** | 0.1520 | 3.0E-14 | **0.0074** | 0.1257 | 1.1E-12 | **0.4691** | 0.5270 | 2.6E-34 | **0.4563** | 0.5101 | 3.5E-30 | **0.5083** | 0.5547 | 1.1E-31 |
| BBS30010 | **0.0013** | 0.0015 | 2.0E-118 | **0.0003** | 0.0008 | 2.7E-131 | **0.0845** | 0.1556 | 7.2E-144 | **0.0731** | 0.1495 | 5.1E-146 | **0.1404** | 0.2296 | 6.0E-148 |
| BBS30013 | **0.0135** | 0.0250 | 2.1E-144 | **0.0046** | 0.0059 | 3.7E-56 | **0.3144** | 0.4613 | 3.5E-154 | **0.2862** | 0.4375 | 7.8E-155 | **0.4238** | 0.5453 | 2.4E-155 |
| BBS30016 | **0.1917** | 0.2473 | 1.1E-20 | **0.1741** | 0.2291 | 2.4E-20 | **0.4616** | 0.5881 | 6.4E-35 | **0.4311** | 0.5522 | 6.2E-38 | **0.5044** | 0.6169 | 3.1E-39 |
| BBS30030 | **0.0043** | 0.0193 | 9.6E-155 | **0.0043** | 0.0054 | 1.3E-34 | **0.1262** | 0.2926 | 4.6E-155 | **0.1113** | 0.2790 | 1.8E-155 | **0.2148** | 0.4222 | 6.1E-156 |
| Domains won | **10** | 0 | | **10** | 0 | | **10** | 0 | | **10** | 0 | | **10** | 0 | |
| Tests won | | | | | | | | | | | | | | | |
| | Ankh | BLOSUM | Equal | Ankh | BLOSUM | Equal | Ankh | BLOSUM | Equal | Ankh | BLOSUM | Equal | Ankh | BLOSUM | Equal |
| BBS30001 | **708** | 119 | 173 | **727** | 93 | 180 | **733** | 76 | 191 | **740** | 76 | 184 | **740** | 76 | 184 |
| BBS30004 | **941** | 25 | 34 | **916** | 47 | 37 | **927** | 33 | 40 | **933** | 27 | 40 | **933** | 29 | 38 |
| BBS30005 | **859** | 43 | 98 | **869** | 33 | 98 | **879** | 21 | 100 | **879** | 21 | 100 | **879** | 21 | 100 |
| BBS30006 | **118** | 31 | 4 | **122** | 27 | 4 | **142** | 6 | 5 | **138** | 8 | 7 | **143** | 5 | 5 |
| BBS30008 | **552** | 63 | 15 | **451** | 164 | 15 | **571** | 35 | 24 | **525** | 78 | 27 | **571** | 39 | 20 |
| BBS30009 | **405** | 187 | 111 | **381** | 197 | 125 | **324** | 90 | 289 | **382** | 154 | 167 | **325** | 91 | 287 |
| BBS30010 | **858** | 128 | 14 | **869** | 116 | 15 | **891** | 80 | 29 | **896** | 75 | 29 | **904** | 75 | 21 |
| BBS30013 | **902** | 70 | 28 | **747** | 217 | 36 | **934** | 27 | 39 | **934** | 29 | 37 | **934** | 29 | 37 |
| BBS30016 | **293** | 59 | 314 | **293** | 59 | 314 | **288** | 30 | 348 | **310** | 33 | 323 | **267** | 39 | 360 |
| BBS30030 | **936** | 14 | 50 | **659** | 292 | 49 | **938** | 8 | 54 | **938** | 9 | 53 | **939** | 9 | 52 |
| Total | **6572** | 739 | 841 | **6034** | 1245 | 873 | **6627** | 406 | 1119 | **6675** | 510 | 967 | **6635** | 413 | 1104 |
| Tests won (%) | | | | | | | | | | | | | | | |
| BBS30001 | **70.80** | 11.90 | 17.30 | **72.70** | 9.30 | 18.00 | **73.30** | 7.60 | 19.10 | **74.00** | 7.60 | 18.40 | **74.00** | 7.60 | 18.40 |
| BBS30004 | **94.10** | 2.50 | 3.40 | **91.60** | 4.70 | 3.70 | **92.70** | 3.30 | 4.00 | **93.30** | 2.70 | 4.00 | **93.30** | 2.90 | 3.80 |
| BBS30005 | **85.90** | 4.30 | 9.80 | **86.90** | 3.30 | 9.80 | **87.90** | 2.10 | 10.00 | **87.90** | 2.10 | 10.00 | **87.90** | 2.10 | 10.00 |
| BBS30006 | **77.12** | 20.26 | 2.61 | **79.74** | 17.65 | 2.61 | **92.81** | 3.92 | 3.27 | **90.20** | 5.23 | 4.58 | **93.46** | 3.27 | 3.27 |
| BBS30008 | **87.62** | 10.00 | 2.38 | **71.59** | 26.03 | 2.38 | **90.63** | 5.56 | 3.81 | **83.33** | 12.38 | 4.29 | **90.63** | 6.19 | 3.17 |
| BBS30009 | **57.61** | 26.60 | 15.79 | **54.20** | 28.02 | 17.78 | **46.09** | 12.80 | 41.11 | **54.34** | 21.91 | 23.76 | **46.23** | 12.94 | 40.83 |
| BBS30010 | **85.80** | 12.80 | 1.40 | **86.90** | 11.60 | 1.50 | **89.10** | 8.00 | 2.90 | **89.60** | 7.50 | 2.90 | **90.40** | 7.50 | 2.10 |
| BBS30013 | **90.20** | 7.00 | 2.80 | **74.70** | 21.70 | 3.60 | **93.40** | 2.70 | 3.90 | **93.40** | 2.90 | 3.70 | **93.40** | 2.90 | 3.70 |
| BBS30016 | **43.99** | 8.86 | 47.15 | **43.99** | 8.86 | 47.15 | **43.24** | 4.50 | 52.25 | **46.55** | 4.95 | 48.50 | **40.09** | 5.86 | 54.05 |
| BBS30030 | **93.60** | 1.40 | 5.00 | **65.90** | 29.20 | 4.90 | **93.80** | 0.80 | 5.40 | **93.80** | 0.90 | 5.30 | **93.90** | 0.90 | 5.20 |
| Average | **78.67** | 10.56 | 10.76 | **72.82** | 16.04 | 11.14 | **80.30** | 5.13 | 14.57 | **80.64** | 6.82 | 12.54 | **80.33** | 5.22 | 14.45 |

Supplementary Table 22: Best BLOSUM matrices for the previous table.

| Domain | $d_{cc}$ | $d_d$ | $d_{pos}$ | $d_{seq}$ | $d_{ssp}$ |
|---|---|---|---|---|---|
| BBS30001 | 62 | 45 | 45 | 45 | 45 |
| BBS30004 | 62 | 45 | 45 | 45 | 45 |
| BBS30005 | 45 | 45 | 45 | 45 | 45 |
| BBS30006 | 45 | 45 | 45 | 45 | 45 |
| BBS30008 | 50 | 50 | 50 | 62 | 50 |
| BBS30009 | 50 | 45 | 45 | 62 | 45 |
| BBS30010 | 80 | 80 | 45 | 45 | 45 |
| BBS30013 | 50 | 45 | 45 | 45 | 45 |
| BBS30016 | 50 | 50 | 50 | 50 | 45 |
| BBS30030 | 45 | 45 | 45 | 45 | 45 |
| BLOSUM | 45 | 50 | 62 | 80 | 90 |
| Total | 34 | 10 | 4 | 2 | 0 |

#### 4.5 CDD insert

Supplementary Table 23: Ankh-score vs best BLOSUM on CDD insert dataset.

| Domain | $d_{cc}$ | | | $d_{d}$ | | | $d_{pos}$ | | | $d_{seq}$ | | | $d_{ssp}$ | | |
|---|---|---|---|---|---|---|---|---|---|---|---|---|---|---|---|
| Average distances | | | | | | | | | | | | | | | |
| | Ankh | BLOSUM | P-value | Ankh | BLOSUM | P-value | Ankh | BLOSUM | P-value | Ankh | BLOSUM | P-value | Ankh | BLOSUM | P-value |
| cd00012 | **0.0448** | 0.1495 | 1.6E-273 | **0.0228** | 0.1500 | 1.1E-262 | **0.6829** | 0.9650 | 3.6E-283 | **0.6001** | 0.7612 | 7.8E-165 | **0.8051** | 0.9821 | 4.1E-283 |
| cd00083 | 0.1640 | 0.0312 | 4.2E-01 | 0.1546 | 0.0097 | 2.6E-01 | 0.3986 | 0.3636 | 5.3E-01 | 0.3775 | 0.3466 | 3.9E-01 | 0.4310 | 0.4474 | 3.0E-02 |
| cd00173 | **0.0354** | 0.1872 | 3.3E-25 | **0.0082** | 0.1546 | 5.4E-25 | **0.4242** | 0.6865 | 1.8E-31 | **0.4028** | 0.6497 | 2.0E-31 | **0.5465** | 0.7794 | 5.9E-32 |
| cd00637 | **0.0045** | 0.0204 | 0.0E+00 | **0.0010** | 0.0058 | 0.0E+00 | **0.1313** | 0.4187 | 0.0E+00 | **0.1063** | 0.3988 | 0.0E+00 | **0.2217** | 0.5802 | 0.0E+00 |
| cd01040 | **0.0041** | 0.0385 | 4.8E-261 | **0.0009** | 0.0111 | 6.7E-275 | **0.1162** | 0.4559 | 3.8E-285 | **0.0966** | 0.4405 | 3.9E-286 | **0.2028** | 0.5784 | 1.6E-290 |
| cd01068 | **0.0150** | 0.0369 | 2.2E-18 | **0.0010** | 0.0086 | 9.7E-20 | **0.1275** | 0.3953 | 1.1E-21 | **0.1257** | 0.3931 | 7.7E-22 | **0.2064** | 0.5174 | 4.6E-23 |
| cd10140 | **0.0657** | 0.0844 | 3.9E-56 | **0.0170** | 0.0358 | 2.7E-66 | **0.7565** | 0.8711 | 8.4E-45 | **0.7089** | 0.7865 | 6.6E-11 | **0.8518** | 0.9315 | 3.5E-45 |
| cd21116 | **0.0140** | 0.0476 | 1.9E-36 | **0.0039** | 0.0235 | 1.2E-40 | **0.3478** | 0.6351 | 1.0E-51 | **0.3200** | 0.6120 | 1.2E-50 | **0.4810** | 0.7596 | 3.5E-52 |
| Domains won | **7** | 0 | | **7** | 0 | | **7** | 0 | | **7** | 0 | | **7** | 0 | |
| Tests won | | | | | | | | | | | | | | | |
| | Ankh | BLOSUM | Equal | Ankh | BLOSUM | Equal | Ankh | BLOSUM | Equal | Ankh | BLOSUM | Equal | Ankh | BLOSUM | Equal |
| cd00012 | **1692** | 78 | 0 | **1647** | 123 | 0 | **1712** | 18 | 40 | **1377** | 385 | 8 | **1711** | 19 | 40 |
| cd00083 | 624 | 356 | 101 | 608 | 365 | 108 | 581 | 331 | 169 | 592 | 340 | 149 | 581 | 337 | 163 |
| cd00173 | **262** | 55 | 8 | **263** | 54 | 8 | **255** | 42 | 28 | **254** | 61 | 10 | **256** | 49 | 20 |
| cd00637 | **2993** | 10 | 0 | **2977** | 26 | 0 | **3001** | 1 | 1 | **3000** | 2 | 1 | **3000** | 2 | 1 |
| cd01040 | **1765** | 112 | 14 | **1793** | 82 | 16 | **1798** | 68 | 25 | **1802** | 72 | 17 | **1801** | 68 | 22 |
| cd01068 | **148** | 19 | 4 | **150** | 17 | 4 | **152** | 14 | 5 | **153** | 14 | 4 | **153** | 14 | 4 |
| cd10140 | **505** | 124 | 1 | **534** | 96 | 0 | **424** | 114 | 92 | **339** | 277 | 14 | **419** | 117 | 94 |
| cd21116 | **284** | 41 | 0 | **295** | 30 | 0 | **304** | 21 | 0 | **303** | 22 | 0 | **303** | 22 | 0 |
| Total | **7649** | 439 | 27 | **7659** | 428 | 28 | **7646** | 278 | 191 | **7228** | 833 | 54 | **7643** | 291 | 181 |
| Tests won (%) | | | | | | | | | | | | | | | |
| cd00012 | **95.59** | 4.41 | 0.00 | **93.05** | 6.95 | 0.00 | **96.72** | 1.02 | 2.26 | **77.80** | 21.75 | 0.45 | **96.67** | 1.07 | 2.26 |
| cd00083 | 57.72 | 32.93 | 9.34 | 56.24 | 33.77 | 9.99 | 53.75 | 30.62 | 15.63 | 54.76 | 31.45 | 13.78 | 53.75 | 31.17 | 15.08 |
| cd00173 | **80.62** | 16.92 | 2.46 | **80.92** | 16.62 | 2.46 | **78.46** | 12.92 | 8.62 | **78.15** | 18.77 | 3.08 | **78.77** | 15.08 | 6.15 |
| cd00637 | **99.67** | 0.33 | 0.00 | **99.13** | 0.87 | 0.00 | **99.93** | 0.03 | 0.03 | **99.90** | 0.07 | 0.03 | **99.90** | 0.07 | 0.03 |
| cd01040 | **93.34** | 5.92 | 0.74 | **94.82** | 4.34 | 0.85 | **95.08** | 3.60 | 1.32 | **95.29** | 3.81 | 0.90 | **95.24** | 3.60 | 1.16 |
| cd01068 | **86.55** | 11.11 | 2.34 | **87.72** | 9.94 | 2.34 | **88.89** | 8.19 | 2.92 | **89.47** | 8.19 | 2.34 | **89.47** | 8.19 | 2.34 |
| cd10140 | **80.16** | 19.68 | 0.16 | **84.76** | 15.24 | 0.00 | **67.30** | 18.10 | 14.60 | **53.81** | 43.97 | 2.22 | **66.51** | 18.57 | 14.92 |
| cd21116 | **87.38** | 12.62 | 0.00 | **90.77** | 9.23 | 0.00 | **93.54** | 6.46 | 0.00 | **93.23** | 6.77 | 0.00 | **93.23** | 6.77 | 0.00 |
| Average | **89.04** | 10.14 | 0.81 | **90.17** | 9.03 | 0.81 | **88.56** | 7.19 | 4.25 | **83.95** | 14.76 | 1.29 | **88.54** | 7.62 | 3.84 |

Supplementary Table 24: Best BLOSUM matrices for the previous table.

| Domain | $d_{cc}$ | $d_{d}$ | $d_{pos}$ | $d_{seq}$ | $d_{ssp}$ |
|---|---|---|---|---|---|
| cd00012 | 45 | 45 | 45 | 62 | 45 |
| cd00083 | 45 | 45 | 45 | 45 | 45 |
| cd00173 | 50 | 50 | 50 | 50 | 45 |
| cd00637 | 45 | 45 | 45 | 45 | 45 |
| cd01040 | 45 | 45 | 45 | 45 | 45 |
| cd01068 | 45 | 45 | 45 | 45 | 45 |
| cd10140 | 45 | 45 | 45 | 62 | 45 |
| cd21116 | 45 | 45 | 45 | 45 | 45 |
| BLOSUM | 45 | 50 | 62 | 80 | 90 |
| Total | 34 | 4 | 2 | 0 | 0 |

#### 4.6 BALiBASE RV911

Supplementary Table 25: Ankh-score vs best BLOSUM on BALiBASE RV911 dataset.

| Domain | $d_{cc}$ | | | $d_d$ | | | $d_{pos}$ | | | $d_{seq}$ | | | $d_{ssp}$ | | |
|---|---|---|---|---|---|---|---|---|---|---|---|---|---|---|---|
| Average distances | | | | | | | | | | | | | | | |
| | Ankh | BLOSUM | P-value | Ankh | BLOSUM | P-value | Ankh | BLOSUM | P-value | Ankh | BLOSUM | P-value | Ankh | BLOSUM | P-value |
| BOX001 | **0.0160** | 0.0269 | 5.4E-130 | **0.0076** | 0.0157 | 2.4E-88 | **0.5400** | 0.6674 | 1.4E-161 | **0.4800** | 0.6033 | 1.1E-152 | **0.6890** | 0.7893 | 3.6E-161 |
| BOX022 | **0.0393** | 0.0605 | 6.4E-25 | **0.0129** | 0.0245 | 1.1E-15 | **0.4944** | 0.6299 | 2.7E-60 | **0.5363** | 0.6029 | 5.7E-62 | **0.7076** | 0.7656 | 4.9E-69 |
| BOX032 | **0.0291** | 0.0316 | 3.5E-40 | 0.0120 | 0.0120 | 4.3E-01 | **0.5670** | 0.6147 | 8.4E-147 | **0.5513** | 0.5833 | 3.5E-72 | **0.7193** | 0.7573 | 3.0E-149 |
| BOX034 | **0.0145** | 0.0386 | 4.8E-156 | **0.0118** | 0.0339 | 1.4E-129 | **0.4642** | 0.5698 | 3.9E-155 | **0.4294** | 0.5187 | 3.1E-111 | **0.6301** | 0.7141 | 5.3E-155 |
| BOX035 | **0.0324** | 0.0575 | 1.2E-120 | **0.0482** | 0.0703 | 8.4E-48 | **0.6624** | 0.6939 | 4.4E-165 | 0.5767 | **0.4944** | 1.3E-130 | **0.7881** | 0.7945 | 4.7E-38 |
| BOX046 | **0.0568** | 0.1058 | 2.3E-12 | **0.0407** | 0.1002 | 1.7E-10 | **0.7165** | 0.7605 | 1.4E-24 | **0.6867** | 0.7190 | 2.3E-06 | **0.8109** | 0.8435 | 6.4E-28 |
| BOX060 | **0.1131** | 0.1198 | 3.0E-06 | **0.0752** | 0.0865 | 2.5E-06 | **0.7821** | 0.8251 | 4.1E-111 | 0.7055 | **0.6835** | 2.1E-04 | **0.8752** | 0.8985 | 6.8E-115 |
| BOX063 | **0.0082** | 0.0097 | 2.2E-83 | **0.0025** | 0.0029 | 8.3E-06 | **0.3600** | 0.4279 | 3.3E-165 | **0.3256** | 0.3962 | 3.3E-165 | **0.5161** | 0.5883 | 3.3E-165 |
| BOX076 | **0.0835** | 0.1197 | 1.5E-37 | **0.0653** | 0.1222 | 1.7E-46 | **0.7244** | 0.8044 | 8.2E-101 | **0.6282** | 0.6434 | 5.1E-05 | **0.8261** | 0.8868 | 3.9E-101 |
| BOX096 | **0.0297** | 0.0369 | 7.2E-85 | **0.0117** | 0.0174 | 3.3E-57 | **0.6267** | 0.7403 | 2.1E-162 | **0.5729** | 0.6703 | 5.3E-142 | **0.7623** | 0.8420 | 7.7E-162 |
| BOX115 | **0.0568** | 0.0747 | 3.3E-85 | **0.0375** | 0.0483 | 1.4E-58 | **0.8177** | 0.8228 | 3.0E-14 | 0.7787 | **0.7118** | 6.0E-102 | **0.8964** | 0.9008 | 4.1E-11 |
| BOX121 | **0.0223** | 0.0280 | 6.8E-98 | **0.0211** | 0.0263 | 4.9E-23 | **0.5563** | 0.6260 | 6.3E-152 | **0.5126** | 0.5477 | 1.5E-25 | **0.6936** | 0.7498 | 1.2E-150 |
| BOX122 | **0.0191** | 0.0196 | 1.7E-07 | 0.0048 | **0.0042** | 1.7E-07 | **0.3679** | 0.4328 | 4.8E-143 | **0.3615** | 0.4244 | 1.7E-141 | **0.5333** | 0.5979 | 1.0E-142 |
| BOX123 | **0.0469** | 0.0501 | 1.1E-34 | **0.0217** | 0.0266 | 2.6E-23 | 0.7220 | **0.7148** | 1.0E-70 | **0.6242** | 0.6486 | 9.6E-39 | **0.8319** | 0.8508 | 2.8E-93 |
| BOX142 | **0.0370** | 0.0503 | 1.2E-22 | **0.0231** | 0.0325 | 8.3E-08 | **0.6550** | 0.7381 | 1.2E-93 | **0.5862** | 0.6606 | 1.3E-90 | **0.7626** | 0.8345 | 8.2E-102 |
| BOX172 | **0.0890** | 0.1055 | 5.2E-47 | **0.1022** | 0.1208 | 3.3E-13 | **0.8343** | 0.8556 | 3.4E-18 | 0.6924 | **0.6259** | 8.8E-37 | **0.8929** | 0.9056 | 4.5E-15 |
| BOX175 | **0.0575** | 0.0587 | 3.5E-09 | 0.0273 | **0.0242** | 3.7E-03 | **0.6981** | 0.7272 | 3.0E-100 | **0.6615** | 0.6724 | 7.7E-09 | **0.8167** | 0.8365 | 3.1E-100 |
| BOX177 | **0.0353** | 0.0406 | 3.5E-45 | **0.0153** | 0.0182 | 7.0E-08 | **0.6708** | 0.7400 | 2.2E-132 | **0.6328** | 0.6770 | 5.9E-75 | **0.7954** | 0.8420 | 1.4E-129 |
| BOX180 | **0.0094** | 0.0260 | 3.9E-158 | **0.0056** | 0.0109 | 5.7E-117 | **0.2992** | 0.5299 | 9.9E-161 | **0.2771** | 0.5101 | 7.5E-161 | **0.4465** | 0.6709 | 3.5E-162 |
| BOX181 | **0.0683** | 0.0741 | 6.2E-14 | **0.0190** | 0.0236 | 2.6E-36 | **0.5716** | 0.6378 | 1.7E-118 | **0.5591** | 0.6174 | 6.5E-108 | **0.7250** | 0.7772 | 5.1E-127 |
| BOX192 | **0.0587** | 0.0736 | 6.1E-76 | **0.0422** | 0.0512 | 1.7E-44 | **0.6887** | 0.7563 | 9.3E-140 | 0.6005 | **0.5719** | 3.2E-18 | **0.8049** | 0.8511 | 6.5E-140 |
| BOX212 | **0.0562** | 0.0750 | 4.8E-55 | **0.0240** | 0.0388 | 1.2E-50 | **0.7332** | 0.7494 | 2.7E-11 | 0.6222 | **0.5825** | 3.9E-71 | **0.8346** | 0.8422 | 9.4E-07 |
| BOX214 | **0.0374** | 0.0596 | 1.9E-93 | **0.0392** | 0.0579 | 1.8E-55 | **0.5138** | 0.6411 | 5.9E-128 | **0.4244** | 0.4874 | 1.1E-68 | **0.6650** | 0.7606 | 2.6E-127 |
| BOX222 | **0.0232** | 0.0622 | 3.6E-105 | **0.0212** | 0.0639 | 4.6E-66 | **0.6460** | 0.7401 | 5.2E-159 | **0.4990** | 0.5387 | 2.8E-38 | **0.7655** | 0.8385 | 4.1E-159 |
| BOX240 | **0.0530** | 0.0677 | 5.6E-29 | **0.0204** | 0.0300 | 2.4E-32 | **0.6969** | 0.7321 | 6.9E-71 | 0.6455 | 0.6509 | 1.8E-02 | **0.8014** | 0.8297 | 2.7E-66 |
| BOX246 | **0.0387** | 0.0392 | 3.8E-11 | **0.0189** | 0.0206 | 5.4E-13 | **0.5935** | 0.6194 | 2.1E-118 | 0.5589 | **0.5338** | 4.1E-57 | **0.7349** | 0.7496 | 6.7E-55 |
| BOX258 | **0.0233** | 0.0312 | 2.0E-91 | **0.0181** | 0.0187 | 3.4E-24 | **0.4435** | 0.5267 | 7.1E-157 | **0.4168** | 0.4623 | 2.0E-24 | **0.6096** | 0.6923 | 4.5E-155 |
| BOX270 | **0.0451** | 0.0523 | 2.1E-25 | **0.0225** | 0.0292 | 4.4E-30 | **0.6211** | 0.6366 | 3.4E-106 | 0.5833 | **0.5514** | 4.6E-38 | **0.7564** | 0.7621 | 5.7E-62 |
| BOX284 | **0.0378** | 0.0766 | 1.6E-151 | **0.0190** | 0.0537 | 6.7E-161 | **0.5848** | 0.6971 | 3.2E-140 | **0.5032** | 0.6025 | 1.6E-114 | **0.7064** | 0.8005 | 1.4E-140 |
| Domains won | **29** | 0 | | **26** | 2 | | **28** | 1 | | **20** | 8 | | **29** | 0 | |

| Tests won | | | | | | | | | | | | | | | |
|---|---|---|---|---|---|---|---|---|---|---|---|---|---|---|---|
| | Ankh | BLOSUM | Equal | Ankh | BLOSUM | Equal | Ankh | BLOSUM | Equal | Ankh | BLOSUM | Equal | Ankh | BLOSUM | Equal |
| BOX001 | **876** | 124 | 0 | **762** | 238 | 0 | **964** | 36 | 0 | **924** | 76 | 0 | **963** | 37 | 0 |
| BOX022 | **653** | 300 | 47 | **594** | 359 | 47 | **775** | 156 | 69 | **761** | 186 | 53 | **780** | 162 | 58 |
| BOX032 | **674** | 325 | 1 | 435 | 565 | 0 | **885** | 109 | 6 | **769** | 207 | 24 | **888** | 107 | 5 |
| BOX034 | **944** | 56 | 0 | **868** | 132 | 0 | **965** | 35 | 0 | **821** | 153 | 26 | **940** | 60 | 0 |
| BOX035 | **871** | 129 | 0 | **649** | 351 | 0 | **995** | 5 | 0 | 68 | **927** | 5 | **598** | 402 | 0 |
| BOX046 | **649** | 351 | 0 | **607** | 393 | 0 | **545** | 224 | 231 | **508** | 486 | 6 | **543** | 226 | 231 |
| BOX060 | **597** | 403 | 0 | **578** | 422 | 0 | **846** | 88 | 66 | 473 | **522** | 5 | **839** | 159 | 2 |
| BOX063 | **775** | 225 | 0 | **597** | 403 | 0 | **1000** | 0 | 0 | **1000** | 0 | 0 | **1000** | 0 | 0 |
| BOX076 | **700** | 279 | 21 | **705** | 275 | 20 | **762** | 103 | 135 | **538** | 435 | 27 | **754** | 121 | 125 |
| BOX096 | **801** | 199 | 0 | **670** | 330 | 0 | **973** | 25 | 2 | **894** | 105 | 1 | **976** | 23 | 1 |
| BOX115 | **775** | 225 | 0 | **657** | 343 | 0 | **695** | 297 | 8 | 122 | **871** | 7 | **622** | 378 | 0 |
| BOX121 | **823** | 177 | 0 | **551** | 449 | 0 | **921** | 62 | 17 | **603** | 395 | 2 | **911** | 89 | 0 |
| BOX122 | **584** | 412 | 4 | 336 | **660** | 4 | **935** | 32 | 33 | **927** | 48 | 25 | **934** | 32 | 34 |
| BOX123 | **741** | 259 | 0 | **631** | 369 | 0 | **701** | 298 | 1 | **686** | 314 | 0 | **796** | 204 | 0 |
| BOX142 | **778** | 222 | 0 | **640** | 360 | 0 | **854** | 126 | 20 | **842** | 155 | 3 | **854** | 122 | 24 |
| BOX172 | **686** | 314 | 0 | **631** | 369 | 0 | **481** | 220 | 299 | 319 | **677** | 4 | **498** | 197 | 305 |
| BOX175 | **651** | 349 | 0 | 486 | **514** | 0 | **738** | 254 | 8 | **603** | 397 | 0 | **737** | 263 | 0 |
| BOX177 | **697** | 303 | 0 | **520** | 480 | 0 | **873** | 107 | 20 | **734** | 256 | 10 | **854** | 146 | 0 |
| BOX180 | **982** | 18 | 0 | **848** | 152 | 0 | **970** | 30 | 0 | **970** | 30 | 0 | **969** | 31 | 0 |
| BOX181 | **571** | 429 | 0 | **664** | 336 | 0 | **809** | 148 | 43 | **786** | 200 | 14 | **820** | 180 | 0 |
| BOX192 | **758** | 242 | 0 | **655** | 345 | 0 | **898** | 68 | 34 | 381 | **613** | 6 | **905** | 94 | 1 |
| BOX212 | **704** | 296 | 0 | **698** | 302 | 0 | **569** | 376 | 55 | 216 | **781** | 3 | **547** | 414 | 39 |
| BOX214 | **848** | 139 | 13 | **732** | 255 | 13 | **877** | 63 | 60 | **749** | 219 | 32 | **866** | 79 | 55 |
| BOX222 | **844** | 156 | 0 | **699** | 301 | 0 | **972** | 26 | 2 | **697** | 301 | 2 | **961** | 39 | 0 |
| BOX240 | **655** | 309 | 36 | **669** | 295 | 36 | **781** | 162 | 57 | 484 | 474 | 42 | **754** | 205 | 41 |
| BOX246 | **621** | 379 | 0 | **590** | 410 | 0 | **868** | 106 | 26 | 267 | **728** | 5 | **685** | 315 | 0 |
| BOX258 | **802** | 198 | 0 | **625** | 375 | 0 | **937** | 53 | 10 | **548** | 437 | 15 | **937** | 57 | 6 |
| BOX270 | **643** | 357 | 0 | **638** | 362 | 0 | **867** | 113 | 20 | 310 | **685** | 5 | **730** | 269 | 1 |
| BOX284 | **913** | 87 | 0 | **991** | 9 | 0 | **863** | 137 | 0 | **802** | 186 | 12 | **866** | 130 | 4 |
| Total | **21616** | 7262 | 122 | **18291** | 9589 | 120 | **24319** | 3459 | 1222 | **17318** | 10390 | 292 | **23527** | 4541 | 932 |

| Tests won (%) | | | | | | | | | | | | | | | |
|---|---|---|---|---|---|---|---|---|---|---|---|---|---|---|---|
| BOX001 | **87.60** | 12.40 | 0.00 | **76.20** | 23.80 | 0.00 | **96.40** | 3.60 | 0.00 | **92.40** | 7.60 | 0.00 | **96.30** | 3.70 | 0.00 |
| BOX022 | **65.30** | 30.00 | 4.70 | **59.40** | 35.90 | 4.70 | **77.50** | 15.60 | 6.90 | **76.10** | 18.60 | 5.30 | **78.00** | 16.20 | 5.80 |
| BOX032 | **67.40** | 32.50 | 0.10 | 43.50 | 56.50 | 0.00 | **88.50** | 10.90 | 0.60 | **76.90** | 20.70 | 2.40 | **88.80** | 10.70 | 0.50 |
| BOX034 | **94.40** | 5.60 | 0.00 | **86.80** | 13.20 | 0.00 | **96.50** | 3.50 | 0.00 | **82.10** | 15.30 | 2.60 | **94.00** | 6.00 | 0.00 |
| BOX035 | **87.10** | 12.90 | 0.00 | **64.90** | 35.10 | 0.00 | **99.50** | 0.50 | 0.00 | 6.80 | **92.70** | 0.50 | **59.80** | 40.20 | 0.00 |
| BOX046 | **64.90** | 35.10 | 0.00 | **60.70** | 39.30 | 0.00 | **54.50** | 22.40 | 23.10 | **50.80** | 48.60 | 0.60 | **54.30** | 22.60 | 23.10 |
| BOX060 | **59.70** | 40.30 | 0.00 | **57.80** | 42.20 | 0.00 | **84.60** | 8.80 | 6.60 | 47.30 | **52.20** | 0.50 | **83.90** | 15.90 | 0.20 |
| BOX063 | **77.50** | 22.50 | 0.00 | **59.70** | 40.30 | 0.00 | **100.00** | 0.00 | 0.00 | **100.00** | 0.00 | 0.00 | **100.00** | 0.00 | 0.00 |
| BOX076 | **70.00** | 27.90 | 2.10 | **70.50** | 27.50 | 2.00 | **76.20** | 10.30 | 13.50 | **53.80** | 43.50 | 2.70 | **75.40** | 12.10 | 12.50 |
| BOX096 | **80.10** | 19.90 | 0.00 | **67.00** | 33.00 | 0.00 | **97.30** | 2.50 | 0.20 | **89.40** | 10.50 | 0.10 | **97.60** | 2.30 | 0.10 |
| BOX115 | **77.50** | 22.50 | 0.00 | **65.70** | 34.30 | 0.00 | **69.50** | 29.70 | 0.80 | 12.20 | **87.10** | 0.70 | **62.20** | 37.80 | 0.00 |
| BOX121 | **82.30** | 17.70 | 0.00 | **55.10** | 44.90 | 0.00 | **92.10** | 6.20 | 1.70 | **60.30** | 39.50 | 0.20 | **91.10** | 8.90 | 0.00 |
| BOX122 | **58.40** | 41.20 | 0.40 | 33.60 | **66.00** | 0.40 | **93.50** | 3.20 | 3.30 | **92.70** | 4.80 | 2.50 | **93.40** | 3.20 | 3.40 |
| BOX123 | **74.10** | 25.90 | 0.00 | **63.10** | 36.90 | 0.00 | **70.10** | 29.80 | 0.10 | **68.60** | 31.40 | 0.00 | **79.60** | 20.40 | 0.00 |
| BOX142 | **77.80** | 22.20 | 0.00 | **64.00** | 36.00 | 0.00 | **85.40** | 12.60 | 2.00 | **84.20** | 15.50 | 0.30 | **85.40** | 12.20 | 2.40 |
| BOX172 | **68.60** | 31.40 | 0.00 | **63.10** | 36.90 | 0.00 | **48.10** | 22.00 | 29.90 | 31.90 | **67.70** | 0.40 | **49.80** | 19.70 | 30.50 |
| BOX175 | **65.10** | 34.90 | 0.00 | 48.60 | **51.40** | 0.00 | **73.80** | 25.40 | 0.80 | **60.30** | 39.70 | 0.00 | **73.70** | 26.30 | 0.00 |
| BOX177 | **69.70** | 30.30 | 0.00 | **52.00** | 48.00 | 0.00 | **87.30** | 10.70 | 2.00 | **73.40** | 25.60 | 1.00 | **85.40** | 14.60 | 0.00 |
| BOX180 | **98.20** | 1.80 | 0.00 | **84.80** | 15.20 | 0.00 | **97.00** | 3.00 | 0.00 | **97.00** | 3.00 | 0.00 | **96.90** | 3.10 | 0.00 |
| BOX181 | **57.10** | 42.90 | 0.00 | **66.40** | 33.60 | 0.00 | **80.90** | 14.80 | 4.30 | **78.60** | 20.00 | 1.40 | **82.00** | 18.00 | 0.00 |
| BOX192 | **75.80** | 24.20 | 0.00 | **65.50** | 34.50 | 0.00 | **89.80** | 6.80 | 3.40 | 38.10 | **61.30** | 0.60 | **90.50** | 9.40 | 0.10 |
| BOX212 | **70.40** | 29.60 | 0.00 | **69.80** | 30.20 | 0.00 | **56.90** | 37.60 | 5.50 | 21.60 | **78.10** | 0.30 | **54.70** | 41.40 | 3.90 |
| BOX214 | **84.80** | 13.90 | 1.30 | **73.20** | 25.50 | 1.30 | **87.70** | 6.30 | 6.00 | **74.90** | 21.90 | 3.20 | **86.60** | 7.90 | 5.50 |
| BOX222 | **84.40** | 15.60 | 0.00 | **69.90** | 30.10 | 0.00 | **97.20** | 2.60 | 0.20 | **69.70** | 30.10 | 0.20 | **96.10** | 3.90 | 0.00 |
| BOX240 | **65.50** | 30.90 | 3.60 | **66.90** | 29.50 | 3.60 | **78.10** | 16.20 | 5.70 | 48.40 | 47.40 | 4.20 | **75.40** | 20.50 | 4.10 |
| BOX246 | **62.10** | 37.90 | 0.00 | **59.00** | 41.00 | 0.00 | **86.80** | 10.60 | 2.60 | 26.70 | **72.80** | 0.50 | **68.50** | 31.50 | 0.00 |
| BOX258 | **80.20** | 19.80 | 0.00 | **62.50** | 37.50 | 0.00 | **93.70** | 5.30 | 1.00 | **54.80** | 43.70 | 1.50 | **93.70** | 5.70 | 0.60 |
| BOX270 | **64.30** | 35.70 | 0.00 | **63.80** | 36.20 | 0.00 | **86.70** | 11.30 | 2.00 | 31.00 | **68.50** | 0.50 | **73.00** | 26.90 | 0.10 |
| BOX284 | **91.30** | 8.70 | 0.00 | **99.10** | 0.90 | 0.00 | **86.30** | 13.70 | 0.00 | **80.20** | 18.60 | 1.20 | **86.60** | 13.00 | 0.40 |
| Average | **74.54** | 25.04 | 0.42 | **65.33** | 34.25 | 0.43 | **83.86** | 11.93 | 4.21 | **61.85** | 37.11 | 1.04 | **81.13** | 15.66 | 3.21 |

Supplementary Table 26: Best BLOSUM matrices for the previous table.

| Domain | $d_{cc}$ | $d_d$ | $d_{pos}$ | $d_{seq}$ | $d_{ssp}$ |
|---|---|---|---|---|---|
| BOX001 | 45 | 45 | 45 | 50 | 45 |
| BOX022 | 45 | 45 | 45 | 45 | 45 |
| BOX032 | 50 | 50 | 45 | 62 | 45 |
| BOX034 | 45 | 45 | 45 | 62 | 45 |
| BOX035 | 45 | 45 | 62 | 62 | 62 |
| BOX046 | 45 | 45 | 45 | 62 | 45 |
| BOX060 | 50 | 45 | 45 | 90 | 45 |
| BOX063 | 45 | 45 | 62 | 62 | 62 |
| BOX076 | 45 | 45 | 45 | 62 | 50 |
| BOX096 | 50 | 50 | 45 | 62 | 45 |
| BOX115 | 45 | 45 | 45 | 80 | 50 |
| BOX121 | 50 | 50 | 45 | 62 | 45 |
| BOX122 | 45 | 45 | 45 | 45 | 45 |
| BOX123 | 45 | 45 | 45 | 62 | 45 |
| BOX142 | 45 | 45 | 50 | 50 | 50 |
| BOX172 | 45 | 45 | 45 | 80 | 45 |
| BOX175 | 50 | 50 | 45 | 90 | 45 |
| BOX177 | 45 | 45 | 45 | 45 | 45 |
| BOX180 | 45 | 45 | 45 | 45 | 45 |
| BOX181 | 45 | 50 | 62 | 62 | 62 |
| BOX192 | 50 | 45 | 45 | 90 | 45 |
| BOX212 | 45 | 45 | 45 | 62 | 45 |
| BOX214 | 45 | 45 | 45 | 45 | 45 |
| BOX222 | 50 | 50 | 50 | 62 | 50 |
| BOX240 | 45 | 45 | 45 | 62 | 45 |
| BOX246 | 45 | 45 | 45 | 62 | 62 |
| BOX258 | 45 | 50 | 62 | 62 | 62 |
| BOX270 | 45 | 45 | 45 | 80 | 62 |
| BOX284 | 45 | 50 | 45 | 62 | 45 |
| BLOSUM | 45 | 50 | 62 | 80 | 90 |
| Total | 90 | 23 | 26 | 3 | 3 |

#### 4.7 BALiBASE RV912

Supplementary Table 27: Ankh-score vs best BLOSUM on BALiBASE RV912 dataset.

| Domain | $d_{cc}$ | | | $d_{d}$ | | | $d_{pos}$ | | | $d_{seq}$ | | | $d_{ssp}$ | | |
|---|---|---|---|---|---|---|---|---|---|---|---|---|---|---|---|
| Average distances | | | | | | | | | | | | | | | |
| | Ankh | BLOSUM | P-value | Ankh | BLOSUM | P-value | Ankh | BLOSUM | P-value | Ankh | BLOSUM | P-value | Ankh | BLOSUM | P-value |
| BOX011 | **0.0089** | 0.0244 | 1.8E-138 | **0.0070** | 0.0105 | 3.8E-44 | **0.2674** | 0.4070 | 1.1E-163 | **0.2340** | 0.3821 | 3.3E-161 | **0.4009** | 0.5598 | 7.1E-164 |
| BOX032 | **0.0032** | 0.0051 | 5.0E-122 | 0.0028 | **0.0024** | 1.5E-13 | **0.2445** | 0.2814 | 9.5E-122 | **0.2219** | 0.2628 | 3.0E-131 | **0.3748** | 0.4238 | 1.4E-132 |
| BOX045 | **0.0350** | 0.0481 | 1.4E-85 | **0.0254** | 0.0384 | 2.8E-74 | **0.4550** | 0.5465 | 6.3E-143 | **0.3448** | 0.4285 | 2.7E-98 | **0.5913** | 0.6819 | 1.4E-138 |
| BOX047 | 0.0142 | **0.0135** | 1.4E-26 | 0.0045 | **0.0037** | 4.3E-42 | 0.3206 | **0.3003** | 3.2E-35 | 0.2843 | **0.2622** | 1.7E-41 | 0.4555 | **0.4306** | 3.6E-49 |
| BOX049 | 0.0161 | **0.0144** | 6.6E-03 | 0.0078 | **0.0051** | 8.3E-17 | 0.2428 | **0.2121** | 1.8E-13 | 0.2287 | **0.1934** | 1.9E-12 | 0.3722 | **0.3384** | 2.1E-09 |
| BOX050 | **0.0150** | 0.0170 | 1.5E-04 | **0.0066** | 0.0115 | 5.8E-26 | **0.3417** | 0.3681 | 4.0E-11 | **0.3087** | 0.3296 | 4.9E-10 | **0.4963** | 0.5230 | 5.2E-16 |
| BOX054 | **0.0118** | 0.0165 | 1.7E-127 | **0.0041** | 0.0050 | 2.8E-31 | **0.3544** | 0.4361 | 9.0E-165 | **0.3006** | 0.3861 | 4.2E-165 | **0.5050** | 0.5910 | 6.5E-165 |
| BOX060 | **0.0097** | 0.0229 | 4.1E-157 | **0.0065** | 0.0093 | 2.2E-79 | **0.2025** | 0.3198 | 5.1E-160 | **0.1575** | 0.2851 | 3.9E-161 | **0.3086** | 0.4760 | 7.0E-163 |
| BOX075 | **0.0089** | 0.0134 | 5.5E-136 | **0.0018** | 0.0036 | 4.9E-80 | **0.2307** | 0.2998 | 6.1E-164 | **0.1833** | 0.2648 | 1.1E-164 | **0.3541** | 0.4473 | 8.7E-165 |
| BOX076 | **0.0091** | 0.0111 | 3.9E-65 | **0.0022** | 0.0031 | 3.5E-76 | **0.1807** | 0.2455 | 2.4E-163 | **0.1570** | 0.2179 | 2.5E-160 | **0.2965** | 0.3805 | 8.0E-163 |
| BOX096 | **0.0074** | 0.0081 | 6.4E-30 | **0.0007** | 0.0015 | 5.7E-32 | **0.1818** | 0.2256 | 2.5E-143 | **0.1533** | 0.2032 | 1.6E-158 | **0.2898** | 0.3610 | 9.4E-159 |
| BOX121 | **0.0093** | 0.0151 | 1.8E-97 | **0.0070** | 0.0113 | 4.7E-60 | **0.3123** | 0.3412 | 9.2E-144 | **0.2441** | 0.2659 | 1.4E-31 | **0.4694** | 0.4699 | 1.8E-126 |
| BOX122 | **0.0077** | 0.0119 | 9.2E-138 | **0.0010** | 0.0025 | 4.1E-139 | **0.1363** | 0.2414 | 6.9E-164 | **0.1303** | 0.2341 | 3.2E-163 | **0.2350** | 0.3813 | 8.5E-164 |
| BOX142 | **0.0063** | 0.0122 | 2.2E-146 | **0.0018** | 0.0026 | 8.6E-17 | **0.2768** | 0.3241 | 5.8E-67 | **0.2644** | 0.3038 | 3.9E-75 | **0.4196** | 0.4953 | 5.5E-117 |
| BOX149 | **0.0075** | 0.0091 | 1.0E-32 | 0.0021 | **0.0020** | 2.2E-07 | **0.2236** | 0.2503 | 2.1E-154 | **0.1896** | 0.2338 | 2.9E-148 | **0.3490** | 0.4125 | 3.9E-155 |
| BOX154 | **0.0193** | 0.0225 | 1.9E-36 | 0.0198 | **0.0151** | 1.1E-52 | **0.5219** | 0.5381 | 8.2E-34 | 0.4543 | **0.4338** | 4.6E-13 | **0.6634** | 0.6822 | 2.7E-28 |
| BOX175 | **0.0339** | 0.0375 | 4.4E-30 | **0.0116** | 0.0129 | 1.3E-11 | **0.5176** | 0.5583 | 2.0E-99 | **0.4948** | 0.5164 | 8.1E-48 | **0.6855** | 0.7097 | 2.2E-90 |
| BOX177 | 0.0101 | **0.0092** | 2.1E-21 | 0.0057 | **0.0032** | 6.1E-07 | **0.2933** | 0.3420 | 3.4E-141 | **0.2845** | 0.3197 | 3.2E-98 | **0.4310** | 0.4829 | 4.5E-118 |
| BOX187 | 0.0282 | **0.0280** | 1.8E-20 | **0.0061** | 0.0066 | 4.6E-05 | **0.3221** | 0.3698 | 2.2E-66 | **0.3090** | 0.3577 | 2.5E-66 | **0.4828** | 0.5339 | 1.2E-59 |
| BOX192 | **0.0493** | 0.0592 | 1.5E-85 | **0.0166** | 0.0298 | 3.8E-122 | **0.6134** | 0.6725 | 5.9E-114 | **0.5646** | 0.6040 | 1.4E-51 | **0.7553** | 0.7984 | 1.4E-112 |
| BOX202 | 0.0609 | **0.0605** | 1.6E-03 | **0.0259** | 0.0267 | 5.1E-03 | **0.4389** | 0.4994 | 7.6E-126 | **0.3975** | 0.4389 | 3.3E-36 | **0.5823** | 0.6512 | 1.8E-122 |
| BOX214 | **0.0069** | 0.0233 | 3.7E-104 | **0.0022** | 0.0060 | 8.1E-135 | **0.2039** | 0.2968 | 3.2E-151 | **0.1918** | 0.2787 | 9.1E-133 | **0.3119** | 0.4365 | 3.3E-151 |
| BOX240 | **0.0269** | 0.0295 | 3.6E-47 | **0.0072** | 0.0124 | 3.9E-127 | **0.4028** | 0.4297 | 1.8E-81 | **0.3819** | 0.3921 | 8.0E-18 | **0.5630** | 0.5903 | 6.1E-79 |
| BOX246 | **0.0223** | 0.0291 | 3.5E-97 | **0.0066** | 0.0122 | 3.1E-132 | **0.4035** | 0.4364 | 1.0E-91 | **0.3822** | 0.3936 | 1.4E-13 | **0.5591** | 0.5910 | 3.4E-100 |
| BOX258 | **0.0161** | 0.0194 | 6.0E-56 | **0.0145** | 0.0165 | 2.3E-17 | **0.3550** | 0.4220 | 1.3E-138 | **0.3420** | 0.3729 | 1.8E-14 | **0.5470** | 0.5940 | 3.0E-126 |
| BOX259 | **0.0050** | 0.0091 | 1.3E-116 | **0.0008** | 0.0015 | 1.9E-116 | **0.1041** | 0.1545 | 5.0E-142 | **0.0934** | 0.1425 | 1.4E-135 | **0.1868** | 0.2615 | 1.6E-140 |
| BOX281 | **0.0183** | 0.0264 | 5.2E-122 | **0.0071** | 0.0100 | 4.3E-74 | **0.3541** | 0.4355 | 2.2E-149 | **0.3250** | 0.4005 | 1.6E-154 | **0.5137** | 0.5977 | 3.2E-149 |
| BOX290 | **0.0246** | 0.0339 | 3.6E-53 | **0.0033** | 0.0089 | 7.7E-61 | **0.2475** | 0.2976 | 2.7E-157 | **0.2127** | 0.2834 | 1.4E-162 | **0.3844** | 0.4669 | 3.2E-165 |
| Domains won | **23** | 5 | | **22** | 6 | | **26** | 2 | | **25** | 3 | | **26** | 2 | |
| Tests won | | | | | | | | | | | | | | | |
| | Ankh | BLOSUM | Equal | Ankh | BLOSUM | Equal | Ankh | BLOSUM | Equal | Ankh | BLOSUM | Equal | Ankh | BLOSUM | Equal |
| BOX011 | **936** | 64 | 0 | **673** | 327 | 0 | **989** | 11 | 0 | **969** | 17 | 14 | **989** | 11 | 0 |
| BOX032 | **829** | 171 | 0 | 487 | **513** | 0 | **817** | 182 | 1 | **811** | 186 | 3 | **830** | 170 | 0 |
| BOX045 | **809** | 191 | 0 | **806** | 194 | 0 | **924** | 68 | 8 | **810** | 186 | 4 | **915** | 85 | 0 |
| BOX047 | 287 | **713** | 0 | 324 | **676** | 0 | 344 | **656** | 0 | 343 | **655** | 2 | 346 | **654** | 0 |
| BOX049 | **510** | 465 | 25 | 415 | **560** | 25 | 451 | **492** | 57 | 455 | **500** | 45 | 462 | **499** | 39 |
| BOX050 | **598** | 401 | 1 | **606** | 394 | 0 | **574** | 409 | 17 | **556** | 437 | 7 | **573** | 426 | 1 |
| BOX054 | **874** | 126 | 0 | **722** | 278 | 0 | **988** | 12 | 0 | **992** | 8 | 0 | **991** | 9 | 0 |
| BOX060 | **940** | 56 | 4 | **747** | 243 | 10 | **941** | 53 | 6 | **946** | 52 | 2 | **967** | 26 | 7 |
| BOX075 | **910** | 90 | 0 | **821** | 179 | 0 | **985** | 10 | 5 | **994** | 5 | 1 | **994** | 5 | 1 |
| BOX076 | **735** | 265 | 0 | **742** | 258 | 0 | **968** | 30 | 2 | **949** | 42 | 9 | **956** | 44 | 0 |
| BOX096 | **743** | 257 | 0 | **597** | 403 | 0 | **855** | 144 | 1 | **934** | 65 | 1 | **933** | 67 | 0 |
| BOX121 | **836** | 164 | 0 | **735** | 265 | 0 | **929** | 66 | 5 | **648** | 351 | 1 | **884** | 116 | 0 |
| BOX122 | **911** | 89 | 0 | **899** | 101 | 0 | **981** | 18 | 1 | **975** | 22 | 3 | **980** | 20 | 0 |
| BOX142 | **926** | 74 | 0 | **628** | 372 | 0 | **732** | 266 | 2 | **745** | 252 | 3 | **795** | 205 | 0 |
| BOX149 | **665** | 335 | 0 | **633** | 367 | 0 | **929** | 69 | 2 | **883** | 99 | 18 | **931** | 69 | 0 |
| BOX154 | **703** | 297 | 0 | 221 | **779** | 0 | **674** | 313 | 13 | **578** | 422 | 0 | **693** | 307 | 0 |
| BOX175 | **635** | 364 | 1 | **601** | 399 | 0 | **860** | 128 | 12 | **728** | 266 | 6 | **870** | 129 | 1 |
| BOX177 | **654** | 346 | 0 | **661** | 339 | 0 | **876** | 107 | 17 | **733** | 260 | 7 | **800** | 199 | 1 |
| BOX187 | **614** | 386 | 0 | **533** | 467 | 0 | **732** | 261 | 7 | **727** | 270 | 3 | **734** | 266 | 0 |
| BOX192 | **784** | 216 | 0 | **884** | 116 | 0 | **851** | 147 | 2 | **715** | 283 | 2 | **849** | 151 | 0 |
| BOX202 | **587** | 413 | 0 | **583** | 417 | 0 | **874** | 120 | 6 | **661** | 329 | 10 | **871** | 127 | 2 |
| BOX214 | **871** | 129 | 0 | **906** | 92 | 2 | **930** | 59 | 11 | **841** | 150 | 9 | **950** | 48 | 2 |
| BOX240 | **743** | 257 | 0 | **860** | 140 | 0 | **746** | 245 | 9 | **639** | 357 | 4 | **770** | 230 | 0 |
| BOX246 | **803** | 197 | 0 | **878** | 122 | 0 | **832** | 167 | 1 | **615** | 341 | 44 | **820** | 180 | 0 |
| BOX258 | **733** | 267 | 0 | **580** | 420 | 0 | **880** | 119 | 1 | **570** | 428 | 2 | **851** | 148 | 1 |
| BOX259 | **876** | 123 | 1 | **872** | 128 | 0 | **933** | 63 | 4 | **913** | 64 | 23 | **932** | 67 | 1 |
| BOX281 | **871** | 129 | 0 | **766** | 234 | 0 | **931** | 67 | 2 | **928** | 67 | 5 | **938** | 62 | 0 |
| BOX290 | **596** | 403 | 1 | **653** | 347 | 0 | **986** | 13 | 1 | **985** | 14 | 1 | **1000** | 0 | 0 |
| Total | **20979** | 6988 | 33 | **18833** | 9130 | 37 | **23512** | 4295 | 193 | **21643** | 6128 | 229 | **23624** | 4320 | 56 |
| Tests won (%) | | | | | | | | | | | | | | | |
| BOX011 | **93.60** | 6.40 | 0.00 | **67.30** | 32.70 | 0.00 | **98.90** | 1.10 | 0.00 | **96.90** | 1.70 | 1.40 | **98.90** | 1.10 | 0.00 |
| BOX032 | **82.90** | 17.10 | 0.00 | 48.70 | **51.30** | 0.00 | **81.70** | 18.20 | 0.10 | **81.10** | 18.60 | 0.30 | **83.00** | 17.00 | 0.00 |
| BOX045 | **80.90** | 19.10 | 0.00 | **80.60** | 19.40 | 0.00 | **92.40** | 6.80 | 0.80 | **81.00** | 18.60 | 0.40 | **91.50** | 8.50 | 0.00 |
| BOX047 | 28.70 | **71.30** | 0.00 | 32.40 | **67.60** | 0.00 | 34.40 | **65.60** | 0.00 | 34.30 | **65.50** | 0.20 | 34.60 | **65.40** | 0.00 |
| BOX049 | **51.00** | 46.50 | 2.50 | 41.50 | **56.00** | 2.50 | 45.10 | **49.20** | 5.70 | 45.50 | **50.00** | 4.50 | 46.20 | **49.90** | 3.90 |
| BOX050 | **59.80** | 40.10 | 0.10 | **60.60** | 39.40 | 0.00 | **57.40** | 40.90 | 1.70 | **55.60** | 43.70 | 0.70 | **57.30** | 42.60 | 0.10 |
| BOX054 | **87.40** | 12.60 | 0.00 | **72.20** | 27.80 | 0.00 | **98.80** | 1.20 | 0.00 | **99.20** | 0.80 | 0.00 | **99.10** | 0.90 | 0.00 |
| BOX060 | **94.00** | 5.60 | 0.40 | **74.70** | 24.30 | 1.00 | **94.10** | 5.30 | 0.60 | **94.60** | 5.20 | 0.20 | **96.70** | 2.60 | 0.70 |
| BOX075 | **91.00** | 9.00 | 0.00 | **82.10** | 17.90 | 0.00 | **98.50** | 1.00 | 0.50 | **99.40** | 0.50 | 0.10 | **99.40** | 0.50 | 0.10 |
| BOX076 | **73.50** | 26.50 | 0.00 | **74.20** | 25.80 | 0.00 | **96.80** | 3.00 | 0.20 | **94.90** | 4.20 | 0.90 | **95.60** | 4.40 | 0.00 |
| BOX096 | **74.30** | 25.70 | 0.00 | **59.70** | 40.30 | 0.00 | **85.50** | 14.40 | 0.10 | **93.40** | 6.50 | 0.10 | **93.30** | 6.70 | 0.00 |
| BOX121 | **83.60** | 16.40 | 0.00 | **73.50** | 26.50 | 0.00 | **92.90** | 6.60 | 0.50 | **64.80** | 35.10 | 0.10 | **88.40** | 11.60 | 0.00 |
| BOX122 | **91.10** | 8.90 | 0.00 | **89.90** | 10.10 | 0.00 | **98.10** | 1.80 | 0.10 | **97.50** | 2.20 | 0.30 | **98.00** | 2.00 | 0.00 |
| BOX142 | **92.60** | 7.40 | 0.00 | **62.80** | 37.20 | 0.00 | **73.20** | 26.60 | 0.20 | **74.50** | 25.20 | 0.30 | **79.50** | 20.50 | 0.00 |
| BOX149 | **66.50** | 33.50 | 0.00 | **63.30** | 36.70 | 0.00 | **92.90** | 6.90 | 0.20 | **88.30** | 9.90 | 1.80 | **93.10** | 6.90 | 0.00 |
| BOX154 | **70.30** | 29.70 | 0.00 | 22.10 | **77.90** | 0.00 | **67.40** | 31.30 | 1.30 | **57.80** | 42.20 | 0.00 | **69.30** | 30.70 | 0.00 |
| BOX175 | **63.50** | 36.40 | 0.10 | **60.10** | 39.90 | 0.00 | **86.00** | 12.80 | 1.20 | **72.80** | 26.60 | 0.60 | **87.00** | 12.90 | 0.10 |
| BOX177 | **65.40** | 34.60 | 0.00 | **66.10** | 33.90 | 0.00 | **87.60** | 10.70 | 1.70 | **73.30** | 26.00 | 0.70 | **80.00** | 19.90 | 0.10 |
| BOX187 | **61.40** | 38.60 | 0.00 | **53.30** | 46.70 | 0.00 | **73.20** | 26.10 | 0.70 | **72.70** | 27.00 | 0.30 | **73.40** | 26.60 | 0.00 |
| BOX192 | **78.40** | 21.60 | 0.00 | **88.40** | 11.60 | 0.00 | **85.10** | 14.70 | 0.20 | **71.50** | 28.30 | 0.20 | **84.90** | 15.10 | 0.00 |
| BOX202 | **58.70** | 41.30 | 0.00 | **58.30** | 41.70 | 0.00 | **87.40** | 12.00 | 0.60 | **66.10** | 32.90 | 1.00 | **87.10** | 12.70 | 0.20 |
| BOX214 | **87.10** | 12.90 | 0.00 | **90.60** | 9.20 | 0.20 | **93.00** | 5.90 | 1.10 | **84.10** | 15.00 | 0.90 | **95.00** | 4.80 | 0.20 |
| BOX240 | **74.30** | 25.70 | 0.00 | **86.00** | 14.00 | 0.00 | **74.60** | 24.50 | 0.90 | **63.90** | 35.70 | 0.40 | **77.00** | 23.00 | 0.00 |
| BOX246 | **80.30** | 19.70 | 0.00 | **87.80** | 12.20 | 0.00 | **83.20** | 16.70 | 0.10 | **61.50** | 34.10 | 4.40 | **82.00** | 18.00 | 0.00 |
| BOX258 | **73.30** | 26.70 | 0.00 | **58.00** | 42.00 | 0.00 | **88.00** | 11.90 | 0.10 | **57.00** | 42.80 | 0.20 | **85.10** | 14.80 | 0.10 |
| BOX259 | **87.60** | 12.30 | 0.10 | **87.20** | 12.80 | 0.00 | **93.30** | 6.30 | 0.40 | **91.30** | 6.40 | 2.30 | **93.20** | 6.70 | 0.10 |
| BOX281 | **87.10** | 12.90 | 0.00 | **76.60** | 23.40 | 0.00 | **93.10** | 6.70 | 0.20 | **92.80** | 6.70 | 0.50 | **93.80** | 6.20 | 0.00 |
| BOX290 | **59.60** | 40.30 | 0.10 | **65.30** | 34.70 | 0.00 | **98.60** | 1.30 | 0.10 | **98.50** | 1.40 | 0.10 | **100.00** | 0.00 | 0.00 |
| Average | **74.93** | 24.96 | 0.12 | **67.26** | 32.61 | 0.13 | **83.97** | 15.34 | 0.69 | **77.30** | 21.89 | 0.82 | **84.37** | 15.43 | 0.20 |

Supplementary Table 28: Best BLOSUM matrices for the previous table.

| Domain | $d_{cc}$ | $d_{d}$ | $d_{pos}$ | $d_{seq}$ | $d_{ssp}$ |
|---|---|---|---|---|---|
| BOX011 | 45 | 45 | 45 | 45 | 45 |
| BOX032 | 62 | 45 | 62 | 62 | 62 |
| BOX045 | 45 | 45 | 45 | 62 | 45 |
| BOX047 | 50 | 45 | 45 | 45 | 45 |
| BOX049 | 62 | 62 | 62 | 62 | 62 |
| BOX050 | 45 | 45 | 45 | 45 | 45 |
| BOX054 | 50 | 45 | 45 | 45 | 45 |
| BOX060 | 45 | 45 | 62 | 62 | 45 |
| BOX075 | 45 | 45 | 62 | 62 | 62 |
| BOX076 | 50 | 45 | 45 | 45 | 45 |
| BOX096 | 45 | 45 | 45 | 45 | 45 |
| BOX121 | 45 | 45 | 45 | 62 | 62 |
| BOX122 | 45 | 45 | 45 | 45 | 45 |
| BOX142 | 45 | 45 | 45 | 45 | 62 |
| BOX149 | 45 | 45 | 62 | 62 | 62 |
| BOX154 | 45 | 50 | 45 | 90 | 62 |
| BOX175 | 50 | 50 | 62 | 62 | 62 |
| BOX177 | 50 | 45 | 45 | 62 | 62 |
| BOX187 | 80 | 62 | 62 | 62 | 62 |
| BOX192 | 45 | 45 | 50 | 62 | 50 |
| BOX202 | 50 | 50 | 50 | 62 | 50 |
| BOX214 | 45 | 45 | 45 | 45 | 45 |
| BOX240 | 62 | 45 | 45 | 62 | 45 |
| BOX246 | 50 | 50 | 45 | 62 | 62 |
| BOX258 | 50 | 50 | 45 | 80 | 45 |
| BOX259 | 45 | 62 | 62 | 62 | 62 |
| BOX281 | 45 | 45 | 45 | 62 | 45 |
| BOX290 | 45 | 45 | 45 | 62 | 62 |
| BLOSUM | 45 | 50 | 62 | 80 | 90 |
| Total | 76 | 17 | 44 | 2 | 1 |

#### 4.8 BALiBASE RV913

Supplementary Table 29: Ankh-score vs best BLOSUM on BALiBASE RV913 dataset.

| Domain | $d_{cc}$ | | | $d_d$ | | | $d_{pos}$ | | | $d_{seq}$ | | | $d_{ssp}$ | | |
|---|---|---|---|---|---|---|---|---|---|---|---|---|---|---|---|
| Average distances | | | | | | | | | | | | | | | |
| | Ankh | BLOSUM | P-value | Ankh | BLOSUM | P-value | Ankh | BLOSUM | P-value | Ankh | BLOSUM | P-value | Ankh | BLOSUM | P-value |
| BOX012 | 0.0076 | **0.0069** | 1.1E-08 | **0.0020** | 0.0021 | 3.9E-09 | 0.2134 | **0.2097** | 6.7E-27 | 0.2193 | **0.1960** | 4.9E-17 | 0.3723 | **0.3326** | 4.8E-24 |
| BOX017 | **0.0154** | 0.0202 | 5.6E-28 | **0.0032** | 0.0056 | 4.3E-12 | **0.2872** | 0.3017 | 5.0E-13 | 0.2677 | 0.2688 | 7.2E-01 | **0.4249** | 0.4370 | 2.5E-06 |
| BOX032 | **0.0012** | 0.0022 | 9.2E-100 | 0.0016 | **0.0011** | 1.3E-32 | **0.1005** | 0.1082 | 6.2E-42 | **0.0944** | 0.1017 | 1.4E-34 | **0.1769** | 0.1887 | 4.3E-37 |
| BOX034 | **0.0015** | 0.0028 | 1.3E-143 | **0.0005** | 0.0010 | 3.1E-34 | **0.0938** | 0.1214 | 5.0E-122 | **0.0793** | 0.1079 | 3.4E-120 | **0.1630** | 0.2080 | 5.4E-122 |
| BOX036 | **0.0024** | 0.0032 | 2.9E-66 | **0.0021** | 0.0023 | 3.5E-50 | **0.1005** | 0.1127 | 1.1E-83 | 0.0927 | 0.0960 | 2.8E-01 | **0.1733** | 0.1878 | 1.4E-32 |
| BOX043 | **0.0007** | 0.0052 | 7.1E-116 | **0.0002** | 0.0008 | 1.3E-64 | **0.0583** | 0.0832 | 1.3E-111 | **0.0558** | 0.0818 | 4.5E-119 | **0.1091** | 0.1588 | 1.9E-109 |
| BOX049 | **0.0061** | 0.0064 | 4.3E-10 | 0.0027 | 0.0020 | 4.1E-02 | 0.1115 | 0.1064 | 2.2E-01 | 0.1082 | 0.1040 | 1.1E-01 | 0.1877 | 0.1835 | 1.0E-01 |
| BOX063 | 0.0049 | 0.0040 | 1.3E-01 | 0.0011 | **0.0011** | 1.6E-24 | **0.1259** | 0.1404 | 7.8E-88 | **0.1169** | 0.1306 | 5.0E-80 | **0.2174** | 0.2399 | 8.5E-95 |
| BOX075 | **0.0011** | 0.0024 | 9.3E-40 | **0.0001** | 0.0005 | 1.1E-115 | **0.0391** | 0.0438 | 8.3E-17 | **0.0377** | 0.0417 | 9.2E-13 | **0.0749** | 0.0831 | 2.1E-16 |
| BOX076 | 0.0068 | **0.0054** | 6.7E-30 | 0.0003 | 0.0003 | 8.1E-02 | **0.0786** | 0.0858 | 4.5E-11 | **0.0764** | 0.0827 | 6.7E-08 | **0.1440** | 0.1553 | 1.2E-08 |
| BOX079 | 0.0026 | 0.0026 | 1.7E-01 | 0.0013 | **0.0011** | 9.9E-12 | **0.1089** | 0.1121 | 3.2E-04 | 0.0897 | 0.0907 | 1.4E-01 | 0.1822 | 0.1844 | 2.8E-02 |
| BOX082 | **0.0008** | 0.0010 | 1.8E-34 | **0.0004** | 0.0007 | 1.0E-11 | **0.0568** | 0.0651 | 1.3E-21 | **0.0478** | 0.0535 | 2.6E-07 | **0.1052** | 0.1153 | 5.5E-12 |
| BOX121 | **0.0020** | 0.0054 | 1.3E-149 | **0.0009** | 0.0022 | 2.2E-123 | **0.1399** | 0.1818 | 2.3E-125 | **0.1383** | 0.1805 | 2.1E-124 | **0.2405** | 0.3045 | 2.2E-125 |
| BOX122 | **0.0038** | 0.0045 | 1.2E-76 | **0.0001** | 0.0006 | 1.5E-136 | **0.0453** | 0.0877 | 6.4E-125 | **0.0447** | 0.0866 | 7.1E-124 | **0.0860** | 0.1568 | 1.1E-125 |
| BOX126 | 0.0045 | **0.0041** | 9.9E-16 | 0.0004 | **0.0004** | 7.4E-18 | 0.0680 | **0.0646** | 1.8E-08 | 0.0645 | **0.0592** | 3.1E-09 | 0.1257 | **0.1203** | 6.1E-09 |
| BOX132 | 0.0222 | **0.0173** | 4.7E-26 | 0.0017 | **0.0017** | 2.4E-07 | 0.1504 | 0.1365 | 2.7E-01 | 0.1476 | **0.1341** | 1.1E-03 | 0.2530 | 0.2458 | 1.6E-01 |
| BOX133 | 0.0106 | **0.0097** | 3.9E-09 | **0.0009** | 0.0011 | 1.0E-60 | **0.1393** | 0.1637 | 8.1E-63 | **0.1329** | 0.1576 | 6.3E-51 | **0.2383** | 0.2816 | 2.0E-59 |
| BOX142 | 0.0016 | **0.0013** | 3.7E-04 | 0.0002 | **0.0002** | 1.0E-15 | 0.0631 | **0.0430** | 6.7E-126 | 0.0533 | **0.0359** | 1.1E-122 | 0.1115 | **0.0830** | 3.9E-122 |
| BOX146 | **0.0077** | 0.0083 | 2.0E-27 | 0.0010 | **0.0008** | 5.3E-80 | **0.0792** | 0.1086 | 9.2E-95 | **0.0696** | 0.0942 | 2.3E-79 | **0.1393** | 0.1843 | 2.2E-85 |
| BOX158 | 0.0058 | **0.0051** | 1.1E-09 | **0.0005** | 0.0006 | 2.6E-28 | **0.0619** | 0.0722 | 5.3E-101 | **0.0550** | 0.0628 | 5.0E-68 | **0.1116** | 0.1288 | 1.6E-93 |
| BOX180 | **0.0025** | 0.0026 | 3.7E-22 | **0.0011** | 0.0016 | 2.3E-23 | **0.1131** | 0.1166 | 2.9E-04 | 0.1340 | **0.1024** | 4.0E-11 | 0.2400 | 0.2013 | 9.7E-01 |
| BOX183 | **0.0025** | 0.0070 | 8.1E-118 | **0.0013** | 0.0020 | 4.6E-106 | **0.0989** | 0.1467 | 2.5E-120 | **0.0798** | 0.1374 | 7.3E-127 | **0.1783** | 0.2497 | 5.8E-129 |
| BOX214 | **0.0000** | 0.0006 | 6.8E-75 | 0.0000 | **0.0000** | 4.2E-37 | 0.0031 | **0.0018** | 1.0E-04 | 0.0031 | **0.0018** | 1.0E-04 | **0.0061** | 0.0068 | 1.8E-09 |
| BOX222 | 0.0093 | 0.0089 | 5.4E-02 | **0.0022** | 0.0024 | 4.2E-15 | **0.1939** | 0.2147 | 2.4E-74 | **0.1672** | 0.1827 | 9.5E-53 | **0.3111** | 0.3416 | 4.2E-78 |
| BOX246 | 0.0295 | **0.0229** | 2.2E-14 | 0.0059 | **0.0028** | 1.0E-08 | 0.2239 | **0.2224** | 2.5E-03 | 0.2131 | 0.2035 | 5.9E-02 | 0.3696 | **0.3623** | 4.8E-03 |
| BOX258 | 0.0129 | 0.0122 | 1.6E-01 | **0.0015** | 0.0025 | 3.2E-33 | **0.1704** | 0.1945 | 2.2E-33 | **0.1446** | 0.1618 | 4.8E-45 | **0.2860** | 0.3175 | 1.8E-64 |
| BOX290 | 0.0039 | 0.0041 | 1.1E-01 | 0.0002 | 0.0005 | 4.9E-02 | **0.0641** | 0.0770 | 1.2E-33 | **0.0445** | 0.0556 | 1.2E-20 | **0.1025** | 0.1221 | 2.2E-26 |
| Domains won | **14** | 8 | | **15** | 9 | | **20** | 5 | | **16** | 6 | | **19** | 4 | |
| Tests won | | | | | | | | | | | | | | | |
| | Ankh | BLOSUM | Equal | Ankh | BLOSUM | Equal | Ankh | BLOSUM | Equal | Ankh | BLOSUM | Equal | Ankh | BLOSUM | Equal |
| BOX012 | **503** | 464 | 33 | **581** | 384 | 35 | **576** | 386 | 38 | **560** | 399 | 41 | **589** | 377 | 34 |
| BOX017 | **601** | 399 | 0 | **537** | 463 | 0 | **604** | 362 | 34 | 472 | 479 | 49 | **606** | 393 | 1 |
| BOX032 | **763** | 237 | 0 | 287 | **713** | 0 | **697** | 292 | 11 | **681** | 309 | 10 | **697** | 303 | 0 |
| BOX034 | **925** | 75 | 0 | **757** | 243 | 0 | **853** | 132 | 15 | **841** | 111 | 48 | **842** | 134 | 24 |
| BOX036 | **725** | 258 | 17 | **733** | 255 | 12 | **762** | 215 | 23 | 458 | 482 | 60 | **588** | 397 | 15 |
| BOX043 | **789** | 211 | 0 | **733** | 267 | 0 | **781** | 204 | 15 | **821** | 171 | 8 | **798** | 198 | 4 |
| BOX049 | **549** | 278 | 173 | 520 | 332 | 148 | 524 | 261 | 215 | 519 | 260 | 221 | 524 | 261 | 215 |
| BOX063 | 621 | 379 | 0 | 339 | **661** | 0 | **854** | 142 | 4 | **820** | 158 | 22 | **868** | 131 | 1 |
| BOX075 | **747** | 251 | 2 | **863** | 135 | 2 | **659** | 314 | 27 | **645** | 331 | 24 | **661** | 328 | 11 |
| BOX076 | 430 | **570** | 0 | 535 | 463 | 2 | **543** | 431 | 26 | **527** | 443 | 30 | **542** | 434 | 24 |
| BOX079 | 434 | 496 | 70 | 389 | **543** | 68 | 428 | **441** | 131 | 462 | 436 | 102 | 462 | 442 | 96 |
| BOX082 | **654** | 346 | 0 | 498 | **501** | 1 | **534** | 421 | 45 | **525** | 458 | 17 | **553** | 434 | 13 |
| BOX121 | **947** | 52 | 1 | **853** | 147 | 0 | **876** | 119 | 5 | **879** | 111 | 10 | **881** | 115 | 4 |
| BOX122 | **810** | 156 | 34 | **837** | 118 | 45 | **780** | 147 | 73 | **773** | 169 | 58 | **781** | 168 | 51 |
| BOX126 | 306 | **684** | 10 | 350 | **640** | 10 | 341 | **591** | 68 | 340 | **593** | 67 | 339 | **597** | 64 |
| BOX132 | 379 | **589** | 32 | **525** | 440 | 35 | 471 | 414 | 115 | 450 | **461** | 89 | 480 | 447 | 73 |
| BOX133 | 451 | **547** | 2 | **730** | 268 | 2 | **674** | 285 | 41 | **663** | 314 | 23 | **683** | 306 | 11 |
| BOX142 | 431 | **562** | 7 | 384 | **604** | 12 | 85 | **865** | 50 | 98 | **852** | 50 | 93 | **862** | 45 |
| BOX146 | **675** | 325 | 0 | **812** | 188 | 0 | **887** | 111 | 2 | **881** | 114 | 5 | **889** | 110 | 1 |
| BOX158 | 455 | **545** | 0 | **615** | 385 | 0 | **816** | 170 | 14 | **742** | 239 | 19 | **800** | 193 | 7 |
| BOX180 | **602** | 334 | 64 | **575** | 359 | 66 | 407 | **482** | 111 | 286 | **602** | 112 | 415 | 515 | 70 |
| BOX183 | **797** | 121 | 82 | **799** | 119 | 82 | **785** | 127 | 88 | **803** | 101 | 96 | **805** | 100 | 95 |
| BOX214 | **460** | 179 | 361 | **324** | 243 | 433 | 312 | **323** | 365 | 312 | **323** | 365 | **324** | 313 | 363 |
| BOX222 | 501 | 498 | 1 | 398 | **602** | 0 | **682** | 316 | 2 | **660** | 337 | 3 | **668** | 332 | 0 |
| BOX246 | 483 | **517** | 0 | 498 | **502** | 0 | **682** | 284 | 34 | 682 | 315 | 3 | **703** | 295 | 2 |
| BOX258 | 561 | 438 | 1 | **685** | 315 | 0 | **664** | 326 | 10 | **725** | 270 | 5 | **734** | 264 | 2 |
| BOX290 | 410 | 424 | 166 | 434 | 335 | 231 | **480** | 269 | 251 | **404** | 350 | 246 | **467** | 301 | 232 |
| Total | **13482** | 7700 | 818 | **14102** | 9095 | 803 | **15762** | 7755 | 1483 | **13436** | 7216 | 1348 | **14911** | 7085 | 1004 |
| Tests won (%) | | | | | | | | | | | | | | | |
| BOX012 | **50.30** | 46.40 | 3.30 | **58.10** | 38.40 | 3.50 | **57.60** | 38.60 | 3.80 | **56.00** | 39.90 | 4.10 | **58.90** | 37.70 | 3.40 |
| BOX017 | **60.10** | 39.90 | 0.00 | **53.70** | 46.30 | 0.00 | **60.40** | 36.20 | 3.40 | 47.20 | 47.90 | 4.90 | **60.60** | 39.30 | 0.10 |
| BOX032 | **76.30** | 23.70 | 0.00 | 28.70 | **71.30** | 0.00 | **69.70** | 29.20 | 1.10 | **68.10** | 30.90 | 1.00 | **69.70** | 30.30 | 0.00 |
| BOX034 | **92.50** | 7.50 | 0.00 | **75.70** | 24.30 | 0.00 | **85.30** | 13.20 | 1.50 | **84.10** | 11.10 | 4.80 | **84.20** | 13.40 | 2.40 |
| BOX036 | **72.50** | 25.80 | 1.70 | **73.30** | 25.50 | 1.20 | **76.20** | 21.50 | 2.30 | 45.80 | 48.20 | 6.00 | **58.80** | 39.70 | 1.50 |
| BOX043 | **78.90** | 21.10 | 0.00 | **73.30** | 26.70 | 0.00 | **78.10** | 20.40 | 1.50 | **82.10** | 17.10 | 0.80 | **79.80** | 19.80 | 0.40 |
| BOX049 | **54.90** | 27.80 | 17.30 | 52.00 | 33.20 | 14.80 | 52.40 | 26.10 | 21.50 | 51.90 | 26.00 | 22.10 | 52.40 | 26.10 | 21.50 |
| BOX063 | 62.10 | 37.90 | 0.00 | 33.90 | **66.10** | 0.00 | **85.40** | 14.20 | 0.40 | **82.00** | 15.80 | 2.20 | **86.80** | 13.10 | 0.10 |
| BOX075 | **74.70** | 25.10 | 0.20 | **86.30** | 13.50 | 0.20 | **65.90** | 31.40 | 2.70 | **64.50** | 33.10 | 2.40 | **66.10** | 32.80 | 1.10 |
| BOX076 | 43.00 | **57.00** | 0.00 | 53.50 | 46.30 | 0.20 | **54.30** | 43.10 | 2.60 | **52.70** | 44.30 | 3.00 | **54.20** | 43.40 | 2.40 |
| BOX079 | 43.40 | 49.60 | 7.00 | 38.90 | **54.30** | 6.80 | 42.80 | **44.10** | 13.10 | 46.20 | 43.60 | 10.20 | 46.20 | 44.20 | 9.60 |
| BOX082 | **65.40** | 34.60 | 0.00 | 49.80 | **50.10** | 0.10 | **53.40** | 42.10 | 4.50 | **52.50** | 45.80 | 1.70 | **55.30** | 43.40 | 1.30 |
| BOX121 | **94.70** | 5.20 | 0.10 | **85.30** | 14.70 | 0.00 | **87.60** | 11.90 | 0.50 | **87.90** | 11.10 | 1.00 | **88.10** | 11.50 | 0.40 |
| BOX122 | **81.00** | 15.60 | 3.40 | **83.70** | 11.80 | 4.50 | **78.00** | 14.70 | 7.30 | **77.30** | 16.90 | 5.80 | **78.10** | 16.80 | 5.10 |
| BOX126 | 30.60 | **68.40** | 1.00 | 35.00 | **64.00** | 1.00 | 34.10 | **59.10** | 6.80 | 34.00 | **59.30** | 6.70 | 33.90 | **59.70** | 6.40 |
| BOX132 | 37.90 | **58.90** | 3.20 | **52.50** | 44.00 | 3.50 | 47.10 | 41.40 | 11.50 | 45.00 | **46.10** | 8.90 | 48.00 | 44.70 | 7.30 |
| BOX133 | 45.10 | **54.70** | 0.20 | **73.00** | 26.80 | 0.20 | **67.40** | 28.50 | 4.10 | **66.30** | 31.40 | 2.30 | **68.30** | 30.60 | 1.10 |
| BOX142 | 43.10 | **56.20** | 0.70 | 38.40 | **60.40** | 1.20 | 8.50 | **86.50** | 5.00 | 9.80 | **85.20** | 5.00 | 9.30 | **86.20** | 4.50 |
| BOX146 | **67.50** | 32.50 | 0.00 | **81.20** | 18.80 | 0.00 | **88.70** | 11.10 | 0.20 | **88.10** | 11.40 | 0.50 | **88.90** | 11.00 | 0.10 |
| BOX158 | 45.50 | **54.50** | 0.00 | **61.50** | 38.50 | 0.00 | **81.60** | 17.00 | 1.40 | **74.20** | 23.90 | 1.90 | **80.00** | 19.30 | 0.70 |
| BOX180 | **60.20** | 33.40 | 6.40 | **57.50** | 35.90 | 6.60 | 40.70 | **48.20** | 11.10 | 28.60 | **60.20** | 11.20 | 41.50 | 51.50 | 7.00 |
| BOX183 | **79.70** | 12.10 | 8.20 | **79.90** | 11.90 | 8.20 | **78.50** | 12.70 | 8.80 | **80.30** | 10.10 | 9.60 | **80.50** | 10.00 | 9.50 |
| BOX214 | **46.00** | 17.90 | 36.10 | **32.40** | 24.30 | 43.30 | 31.20 | **32.30** | 36.50 | 31.20 | **32.30** | 36.50 | **32.40** | 31.30 | 36.30 |
| BOX222 | 50.10 | 49.80 | 0.10 | 39.80 | **60.20** | 0.00 | **68.20** | 31.60 | 0.20 | **66.00** | 33.70 | 0.30 | **66.80** | 33.20 | 0.00 |
| BOX246 | 48.30 | **51.70** | 0.00 | 49.80 | **50.20** | 0.00 | **68.20** | 28.40 | 3.40 | 68.20 | 31.50 | 0.30 | **70.30** | 29.50 | 0.20 |
| BOX258 | 56.10 | 43.80 | 0.10 | **68.50** | 31.50 | 0.00 | **66.40** | 32.60 | 1.00 | **72.50** | 27.00 | 0.50 | **73.40** | 26.40 | 0.20 |
| BOX290 | 41.00 | 42.40 | 16.60 | 43.40 | 33.50 | 23.10 | **48.00** | 26.90 | 25.10 | **40.40** | 35.00 | 24.60 | **46.70** | 30.10 | 23.20 |
| Average | **61.28** | 35.00 | 3.72 | **58.76** | 37.90 | 3.35 | **63.05** | 31.02 | 5.93 | **61.07** | 32.80 | 6.13 | **64.83** | 30.80 | 4.37 |

Supplementary Table 30: Best BLOSUM matrices for the previous table.

| Domain | $d_{cc}$ | $d_{d}$ | $d_{pos}$ | $d_{seq}$ | $d_{ssp}$ |
|---|---|---|---|---|---|
| BOX012 | 50 | 62 | 62 | 62 | 62 |
| BOX017 | 90 | 45 | 45 | 45 | 45 |
| BOX032 | 45 | 90 | 45 | 45 | 45 |
| BOX034 | 62 | 62 | 62 | 62 | 62 |
| BOX036 | 50 | 50 | 45 | 62 | 45 |
| BOX043 | 62 | 45 | 45 | 45 | 62 |
| BOX049 | 62 | 50 | 62 | 62 | 62 |
| BOX063 | 50 | 45 | 62 | 62 | 62 |
| BOX075 | 45 | 45 | 45 | 45 | 45 |
| BOX076 | 50 | 45 | 62 | 62 | 62 |
| BOX079 | 62 | 50 | 80 | 80 | 80 |
| BOX082 | 62 | 62 | 62 | 62 | 62 |
| BOX121 | 45 | 45 | 62 | 62 | 62 |
| BOX122 | 62 | 45 | 62 | 62 | 62 |
| BOX126 | 62 | 62 | 62 | 62 | 62 |
| BOX132 | 50 | 50 | 50 | 50 | 50 |
| BOX133 | 50 | 50 | 45 | 45 | 45 |
| BOX142 | 90 | 62 | 90 | 90 | 90 |
| BOX146 | 45 | 62 | 62 | 62 | 62 |
| BOX158 | 50 | 45 | 45 | 62 | 45 |
| BOX180 | 50 | 45 | 62 | 62 | 45 |
| BOX183 | 90 | 50 | 50 | 50 | 50 |
| BOX214 | 50 | 50 | 62 | 62 | 50 |
| BOX222 | 45 | 45 | 45 | 62 | 62 |
| BOX246 | 50 | 50 | 50 | 50 | 50 |
| BOX258 | 45 | 45 | 62 | 62 | 62 |
| BOX290 | 45 | 45 | 45 | 45 | 45 |
| BLOSUM | 45 | 50 | 62 | 80 | 90 |
| Total | 42 | 28 | 55 | 3 | 7 |

### 5 Comparison against GPCRtm

#### 5.1 GPCRtm vs best BLOSUM

Supplementary Table 31: GPCRtm vs best BLOSUM.

| Domain | $d_{cc}$ | | | $d_{d}$ | | | $d_{pos}$ | | | $d_{seq}$ | | | $d_{ssp}$ | | |
|---|---|---|---|---|---|---|---|---|---|---|---|---|---|---|---|
| Average distances | | | | | | | | | | | | | | | |
| | GPCRtm | BLOSUM | P-value | GPCRtm | BLOSUM | P-value | GPCRtm | BLOSUM | P-value | GPCRtm | BLOSUM | P-value | GPCRtm | BLOSUM | P-value |
| Alicarboxylic | **0.0054** | 0.0082 | 1.0E-70 | 0.0024 | 0.0022 | 6.8E-01 | **0.0963** | 0.1293 | 1.0E-94 | **0.0916** | 0.1241 | 4.6E-91 | **0.1684** | 0.2150 | 3.1E-92 |
| Aminergic | **0.0027** | 0.0029 | 3.9E-06 | **0.0009** | 0.0012 | 6.4E-19 | **0.0780** | 0.0891 | 6.7E-24 | **0.0755** | 0.0864 | 5.0E-23 | **0.1431** | 0.1620 | 2.9E-25 |
| Lipid | **0.0219** | 0.0230 | 3.0E-03 | 0.0066 | **0.0059** | 7.8E-25 | **0.4026** | 0.4514 | 2.1E-23 | **0.4013** | 0.4389 | 3.4E-28 | **0.5542** | 0.6106 | 2.5E-30 |
| Nucleotide | 0.0171 | **0.0145** | 8.8E-05 | 0.0081 | **0.0044** | 2.4E-45 | **0.3026** | 0.3196 | 7.3E-19 | **0.2942** | 0.3112 | 5.1E-17 | **0.4169** | 0.4370 | 4.2E-22 |
| Orphan | 0.0668 | **0.0258** | 1.3E-35 | 0.0592 | **0.0074** | 1.0E-67 | 0.5300 | 0.5167 | 7.3E-01 | **0.4997** | 0.5016 | 5.2E-03 | **0.6479** | 0.6648 | 2.2E-03 |
| Peptide | **0.0099** | 0.0125 | 1.4E-41 | **0.0025** | 0.0034 | 3.9E-14 | **0.2126** | 0.2588 | 1.2E-59 | **0.2024** | 0.2484 | 1.1E-58 | **0.3446** | 0.3996 | 6.4E-62 |
| Protein | **0.0069** | 0.0088 | 4.6E-10 | **0.0018** | 0.0022 | 1.2E-03 | **0.1517** | 0.1979 | 1.0E-27 | **0.1427** | 0.1875 | 4.1E-28 | **0.2713** | 0.3181 | 3.8E-24 |
| Sensory | **0.0037** | 0.0054 | 5.7E-81 | **0.0015** | 0.0018 | 7.3E-48 | **0.0852** | 0.1067 | 9.1E-85 | **0.0821** | 0.1043 | 8.3E-86 | **0.1537** | 0.1881 | 3.5E-86 |
| Domains won | **6** | 2 | | **4** | 3 | | **7** | 0 | | **8** | 0 | | **8** | 0 | |
| Tests won | | | | | | | | | | | | | | | |
| | GPCRtm | BLOSUM | Equal | GPCRtm | BLOSUM | Equal | GPCRtm | BLOSUM | Equal | GPCRtm | BLOSUM | Equal | GPCRtm | BLOSUM | Equal |
| Alicarboxylic | **692** | 178 | 130 | 453 | 418 | 129 | **703** | 158 | 139 | **688** | 163 | 149 | **694** | 165 | 141 |
| Aminergic | **554** | 365 | 81 | **559** | 362 | 79 | **569** | 313 | 118 | **571** | 325 | 104 | **578** | 323 | 99 |
| Lipid | **524** | 467 | 9 | 402 | **589** | 9 | **654** | 327 | 19 | **665** | 313 | 22 | **662** | 322 | 16 |
| Nucleotide | **486** | 479 | 35 | 333 | **612** | 55 | **586** | 328 | 86 | **579** | 349 | 72 | **585** | 345 | 70 |
| Orphan | 422 | **575** | 3 | 303 | **694** | 3 | 529 | 442 | 29 | **565** | 426 | 9 | **540** | 438 | 22 |
| Peptide | **690** | 293 | 17 | **611** | 371 | 18 | **704** | 257 | 39 | **706** | 257 | 37 | **710** | 257 | 33 |
| Protein | **611** | 350 | 39 | **576** | 382 | 42 | **635** | 300 | 65 | **624** | 305 | 71 | **639** | 309 | 52 |
| Sensory | **765** | 145 | 90 | **701** | 201 | 98 | **710** | 130 | 160 | **710** | 142 | 148 | **711** | 153 | 136 |
| Total | **4744** | 2852 | 404 | **3485** | 3211 | 304 | **4561** | 1813 | 626 | **5108** | 2280 | 612 | **5119** | 2312 | 569 |
| Tests won (%) | | | | | | | | | | | | | | | |
| Alicarboxylic | **69.20** | 17.80 | 13.00 | 45.30 | 41.80 | 12.90 | **70.30** | 15.80 | 13.90 | **68.80** | 16.30 | 14.90 | **69.40** | 16.50 | 14.10 |
| Aminergic | **55.40** | 36.50 | 8.10 | **55.90** | 36.20 | 7.90 | **56.90** | 31.30 | 11.80 | **57.10** | 32.50 | 10.40 | **57.80** | 32.30 | 9.90 |
| Lipid | **52.40** | 46.70 | 0.90 | 40.20 | **58.90** | 0.90 | **65.40** | 32.70 | 1.90 | **66.50** | 31.30 | 2.20 | **66.20** | 32.20 | 1.60 |
| Nucleotide | **48.60** | 47.90 | 3.50 | 33.30 | **61.20** | 5.50 | **58.60** | 32.80 | 8.60 | **57.90** | 34.90 | 7.20 | **58.50** | 34.50 | 7.00 |
| Orphan | 42.20 | **57.50** | 0.30 | 30.30 | **69.40** | 0.30 | 52.90 | 44.20 | 2.90 | **56.50** | 42.60 | 0.90 | **54.00** | 43.80 | 2.20 |
| Peptide | **69.00** | 29.30 | 1.70 | **61.10** | 37.10 | 1.80 | **70.40** | 25.70 | 3.90 | **70.60** | 25.70 | 3.70 | **71.00** | 25.70 | 3.30 |
| Protein | **61.10** | 35.00 | 3.90 | **57.60** | 38.20 | 4.20 | **63.50** | 30.00 | 6.50 | **62.40** | 30.50 | 7.10 | **63.90** | 30.90 | 5.20 |
| Sensory | **76.50** | 14.50 | 9.00 | **70.10** | 20.10 | 9.80 | **71.00** | 13.00 | 16.00 | **71.00** | 14.20 | 14.80 | **71.10** | 15.30 | 13.60 |
| Average | **59.30** | 35.65 | 5.05 | **49.79** | 45.87 | 4.34 | **65.16** | 25.90 | 8.94 | **63.85** | 28.50 | 7.65 | **63.99** | 28.90 | 7.11 |

Supplementary Table 32: Best BLOSUM matrices for the previous table.

| Domain | $d_{cc}$ | $d_{d}$ | $d_{pos}$ | $d_{seq}$ | $d_{ssp}$ |
|---|---|---|---|---|---|
| cd00012 | 45 | 50 | 45 | 45 | 45 |
| cd00083 | 62 | 62 | 62 | 62 | 62 |
| cd00173 | 45 | 45 | 45 | 45 | 45 |
| cd00637 | 45 | 45 | 62 | 62 | 62 |
| cd01040 | 45 | 45 | 45 | 45 | 45 |
| cd01068 | 62 | 62 | 62 | 62 | 62 |
| cd10140 | 62 | 62 | 62 | 62 | 45 |
| cd21116 | 45 | 62 | 62 | 62 | 62 |
| BLOSUM | 45 | 50 | 62 | 80 | 90 |
| Total | 18 | 1 | 21 | 0 | 0 |

#### 5.2 GPCRtm vs BLOSUM62

Supplementary Table 33: GPCRtm vs BLOSUM62.

| Domain | $d_{cc}$ | | | $d_d$ | | | $d_{pos}$ | | | $d_{seq}$ | | | $d_{ssp}$ | | |
|---|---|---|---|---|---|---|---|---|---|---|---|---|---|---|---|
| Average distances | | | | | | | | | | | | | | | |
| | GPCRtm | BLOSUM62 | P-value | GPCRtm | BLOSUM62 | P-value | GPCRtm | BLOSUM62 | P-value | GPCRtm | BLOSUM62 | P-value | GPCRtm | BLOSUM62 | P-value |
| Alicarboxylic | **0.0054** | 0.0085 | 3.4E-70 | **0.0024** | 0.0027 | 2.6E-59 | **0.0963** | 0.1341 | 4.4E-93 | **0.0916** | 0.1295 | 6.1E-92 | **0.1684** | 0.2216 | 8.2E-91 |
| Aminergic | **0.0027** | 0.0029 | 3.9E-06 | **0.0009** | 0.0012 | 6.4E-19 | **0.0780** | 0.0891 | 6.7E-24 | **0.0755** | 0.0864 | 5.0E-23 | **0.1431** | 0.1620 | 2.9E-25 |
| Lipid | **0.0219** | 0.0287 | 7.3E-13 | **0.0066** | 0.0101 | 1.9E-05 | **0.4026** | 0.4748 | 6.5E-46 | **0.4013** | 0.4556 | 2.1E-46 | **0.5542** | 0.6271 | 1.4E-50 |
| Nucleotide | 0.0171 | **0.0159** | 7.4E-07 | 0.0081 | **0.0064** | 2.4E-04 | **0.3026** | 0.3196 | 7.3E-19 | **0.2942** | 0.3112 | 5.1E-17 | **0.4169** | 0.4370 | 4.2E-22 |
| Orphan | 0.0668 | 0.0635 | 1.3E-02 | 0.0592 | **0.0373** | 3.4E-12 | **0.5300** | 0.5566 | 5.0E-12 | **0.4997** | 0.5299 | 5.0E-15 | **0.6479** | 0.6841 | 1.9E-17 |
| Peptide | **0.0099** | 0.0125 | 1.4E-41 | **0.0025** | 0.0034 | 3.9E-14 | **0.2126** | 0.2588 | 1.2E-59 | **0.2024** | 0.2484 | 1.1E-58 | **0.3446** | 0.3996 | 6.4E-62 |
| Protein | **0.0069** | 0.0088 | 4.6E-10 | **0.0018** | 0.0022 | 1.2E-03 | **0.1517** | 0.1979 | 1.0E-27 | **0.1427** | 0.1875 | 4.1E-28 | **0.2713** | 0.3181 | 1.4E-30 |
| Sensory | **0.0037** | 0.0055 | 8.9E-52 | **0.0015** | 0.0018 | 7.3E-48 | **0.0852** | 0.1067 | 9.1E-85 | **0.0821** | 0.1043 | 8.3E-86 | **0.1537** | 0.1881 | 3.5E-86 |
| Domains won | **6** | 1 | | **6** | 2 | | **8** | 0 | | **8** | 0 | | **8** | 0 | |
| Tests won | | | | | | | | | | | | | | | |
| | GPCRtm | BLOSUM62 | Equal | GPCRtm | BLOSUM62 | Equal | GPCRtm | BLOSUM62 | Equal | GPCRtm | BLOSUM62 | Equal | GPCRtm | BLOSUM62 | Equal |
| Alicarboxylic | **723** | 158 | 119 | **702** | 179 | 119 | **732** | 131 | 137 | **726** | 138 | 136 | **724** | 140 | 136 |
| Aminergic | **554** | 365 | 81 | **559** | 362 | 79 | **569** | 313 | 118 | **571** | 325 | 104 | **578** | 323 | 99 |
| Lipid | **625** | 354 | 21 | **572** | 406 | 22 | **668** | 289 | 43 | **668** | 295 | 37 | **667** | 300 | 33 |
| Nucleotide | **559** | 384 | 57 | **467** | 466 | 67 | **586** | 328 | 86 | **579** | 349 | 72 | **585** | 345 | 70 |
| Orphan | 534 | 456 | 10 | 445 | **544** | 11 | **573** | 354 | 73 | **613** | 361 | 26 | **579** | 359 | 62 |
| Peptide | **690** | 293 | 17 | **611** | 371 | 18 | **704** | 257 | 39 | **706** | 257 | 37 | **710** | 257 | 33 |
| Protein | **611** | 350 | 39 | **576** | 382 | 42 | **635** | 300 | 65 | **624** | 305 | 71 | **638** | 300 | 62 |
| Sensory | **713** | 189 | 98 | **701** | 201 | 98 | **710** | 130 | 160 | **710** | 142 | 148 | **711** | 153 | 136 |
| Total | **4475** | 2093 | 432 | **4633** | 2911 | 456 | **5177** | 2102 | 721 | **5197** | 2172 | 631 | **5192** | 2177 | 631 |
| Tests won (%) | | | | | | | | | | | | | | | |
| Alicarboxylic | **72.30** | 15.80 | 11.90 | **70.20** | 17.90 | 11.90 | **73.20** | 13.10 | 13.70 | **72.60** | 13.80 | 13.60 | **72.40** | 14.00 | 13.60 |
| Aminergic | **55.40** | 36.50 | 8.10 | **55.90** | 36.20 | 7.90 | **56.90** | 31.30 | 11.80 | **57.10** | 32.50 | 10.40 | **57.80** | 32.30 | 9.90 |
| Lipid | **62.50** | 35.40 | 2.10 | **57.20** | 40.60 | 2.20 | **66.80** | 28.90 | 4.30 | **66.80** | 29.50 | 3.70 | **66.70** | 30.00 | 3.30 |
| Nucleotide | **55.90** | 38.40 | 5.70 | **46.70** | 46.60 | 6.70 | **58.60** | 32.80 | 8.60 | **57.90** | 34.90 | 7.20 | **58.50** | 34.50 | 7.00 |
| Orphan | 53.40 | 45.60 | 1.00 | 44.50 | **54.40** | 1.10 | **57.30** | 35.40 | 7.30 | **61.30** | 36.10 | 2.60 | **57.90** | 35.90 | 6.20 |
| Peptide | **69.00** | 29.30 | 1.70 | **61.10** | 37.10 | 1.80 | **70.40** | 25.70 | 3.90 | **70.60** | 25.70 | 3.70 | **71.00** | 25.70 | 3.30 |
| Protein | **61.10** | 35.00 | 3.90 | **57.60** | 38.20 | 4.20 | **63.50** | 30.00 | 6.50 | **62.40** | 30.50 | 7.10 | **63.80** | 30.00 | 6.20 |
| Sensory | **71.30** | 18.90 | 9.80 | **70.10** | 20.10 | 9.80 | **71.00** | 13.00 | 16.00 | **71.00** | 14.20 | 14.80 | **71.10** | 15.30 | 13.60 |
| Average | **63.93** | 29.90 | 6.17 | **57.91** | 36.39 | 5.70 | **64.71** | 26.28 | 9.01 | **64.96** | 27.15 | 7.89 | **64.90** | 27.21 | 7.89 |

#### 5.3 Ankh-score vs GPCRtm

Supplementary Table 34: Ankh-score vs GPCRtm.

| Domain | $d_{cc}$ | | | $d_d$ | | | $d_{pos}$ | | | $d_{seq}$ | | | $d_{ssp}$ | | |
|---|---|---|---|---|---|---|---|---|---|---|---|---|---|---|---|
| Average distances | | | | | | | | | | | | | | | |
| | Ankh | GPCRtm | P-value | Ankh | GPCRtm | P-value | Ankh | GPCRtm | P-value | Ankh | GPCRtm | P-value | Ankh | GPCRtm | P-value |
| Alicarboxylic | **0.0017** | 0.0054 | 2.5E-138 | **0.0024** | 0.0024 | 1.5E-22 | **0.0513** | 0.0963 | 7.9E-117 | **0.0434** | 0.0916 | 6.7E-129 | **0.0928** | 0.1684 | 3.6E-134 |
| Aminergic | **0.0007** | 0.0027 | 7.9E-152 | **0.0003** | 0.0009 | 6.7E-121 | **0.0302** | 0.0780 | 1.6E-157 | **0.0255** | 0.0755 | 5.6E-159 | **0.0598** | 0.1431 | 3.6E-159 |
| Lipid | **0.0053** | 0.0219 | 1.2E-163 | **0.0029** | 0.0066 | 3.1E-147 | **0.1694** | 0.4026 | 7.4E-164 | **0.1489** | 0.4013 | 4.3E-164 | **0.2767** | 0.5542 | 2.1E-164 |
| Nucleotide | **0.0040** | 0.0171 | 7.1E-162 | **0.0026** | 0.0081 | 7.4E-139 | **0.1266** | 0.3026 | 3.1E-163 | **0.1109** | 0.2942 | 1.9E-163 | **0.2074** | 0.4169 | 2.1E-163 |
| Orphan | **0.0060** | 0.0668 | 6.5E-164 | **0.0036** | 0.0592 | 1.2E-154 | **0.1920** | 0.5300 | 8.0E-164 | **0.1670** | 0.4997 | 5.9E-164 | **0.3121** | 0.6479 | 6.2E-164 |
| Peptide | **0.0029** | 0.0099 | 9.9E-163 | **0.0016** | 0.0025 | 4.0E-116 | **0.0995** | 0.2126 | 3.2E-163 | **0.0828** | 0.2024 | 2.3E-163 | **0.1711** | 0.3446 | 1.6E-163 |
| Protein | **0.0014** | 0.0069 | 5.1E-156 | **0.0013** | 0.0018 | 4.9E-133 | **0.0587** | 0.1517 | 1.2E-159 | **0.0423** | 0.1427 | 1.3E-160 | **0.1032** | 0.2713 | 6.2E-161 |
| Sensory | **0.0011** | 0.0037 | 2.2E-115 | **0.0013** | 0.0015 | 2.7E-86 | **0.0502** | 0.0852 | 2.6E-141 | **0.0451** | 0.0821 | 2.0E-142 | **0.0909** | 0.1537 | 8.7E-144 |
| Domains won | **8** | 0 | | **8** | 0 | | **8** | 0 | | **8** | 0 | | **8** | 0 | |
| Tests won | | | | | | | | | | | | | | | |
| | Ankh | GPCRtm | Equal | Ankh | GPCRtm | Equal | Ankh | GPCRtm | Equal | Ankh | GPCRtm | Equal | Ankh | GPCRtm | Equal |
| Alicarboxylic | **837** | 70 | 93 | **647** | 260 | 93 | **753** | 121 | 126 | **805** | 86 | 109 | **816** | 83 | 101 |
| Aminergic | **939** | 47 | 14 | **889** | 96 | 15 | **951** | 30 | 19 | **959** | 25 | 16 | **960** | 25 | 15 |
| Lipid | **980** | 17 | 3 | **914** | 83 | 3 | **980** | 16 | 4 | **983** | 13 | 4 | **985** | 13 | 2 |
| Nucleotide | **979** | 13 | 8 | **871** | 117 | 12 | **984** | 6 | 10 | **985** | 6 | 9 | **986** | 5 | 9 |
| Orphan | **986** | 12 | 2 | **939** | 59 | 2 | **978** | 20 | 2 | **979** | 19 | 2 | **979** | 19 | 2 |
| Peptide | **975** | 21 | 4 | **865** | 131 | 4 | **978** | 13 | 9 | **980** | 11 | 9 | **981** | 11 | 8 |
| Protein | **951** | 31 | 18 | **884** | 97 | 19 | **952** | 29 | 19 | **958** | 21 | 21 | **961** | 19 | 20 |
| Sensory | **779** | 120 | 101 | **710** | 189 | 101 | **830** | 68 | 102 | **833** | 62 | 105 | **846** | 53 | 101 |
| Total | **7426** | 331 | 243 | **6719** | 1032 | 249 | **7406** | 303 | 291 | **7482** | 243 | 275 | **7514** | 228 | 258 |
| Tests won (%) | | | | | | | | | | | | | | | |
| Alicarboxylic | **83.70** | 7.00 | 9.30 | **64.70** | 26.00 | 9.30 | **75.30** | 12.10 | 12.60 | **80.50** | 8.60 | 10.90 | **81.60** | 8.30 | 10.10 |
| Aminergic | **93.90** | 4.70 | 1.40 | **88.90** | 9.60 | 1.50 | **95.10** | 3.00 | 1.90 | **95.90** | 2.50 | 1.60 | **96.00** | 2.50 | 1.50 |
| Lipid | **98.00** | 1.70 | 0.30 | **91.40** | 8.30 | 0.30 | **98.00** | 1.60 | 0.40 | **98.30** | 1.30 | 0.40 | **98.50** | 1.30 | 0.20 |
| Nucleotide | **97.90** | 1.30 | 0.80 | **87.10** | 11.70 | 1.20 | **98.40** | 0.60 | 1.00 | **98.50** | 0.60 | 0.90 | **98.60** | 0.50 | 0.90 |
| Orphan | **98.60** | 1.20 | 0.20 | **93.90** | 5.90 | 0.20 | **97.80** | 2.00 | 0.20 | **97.90** | 1.90 | 0.20 | **97.90** | 1.90 | 0.20 |
| Peptide | **97.50** | 2.10 | 0.40 | **86.50** | 13.10 | 0.40 | **97.80** | 1.30 | 0.90 | **98.00** | 1.10 | 0.90 | **98.10** | 1.10 | 0.80 |
| Protein | **95.10** | 3.10 | 1.80 | **88.40** | 9.70 | 1.90 | **95.20** | 2.90 | 1.90 | **95.80** | 2.10 | 2.10 | **96.10** | 1.90 | 2.00 |
| Sensory | **77.90** | 12.00 | 10.10 | **71.00** | 18.90 | 10.10 | **83.00** | 6.80 | 10.20 | **83.30** | 6.20 | 10.50 | **84.60** | 5.30 | 10.10 |
| Average | **92.83** | 4.14 | 3.04 | **83.99** | 12.90 | 3.11 | **92.58** | 3.79 | 3.64 | **93.53** | 3.04 | 3.44 | **93.93** | 2.85 | 3.23 |

### 6 Comparison against ProtT5-score/PEbA

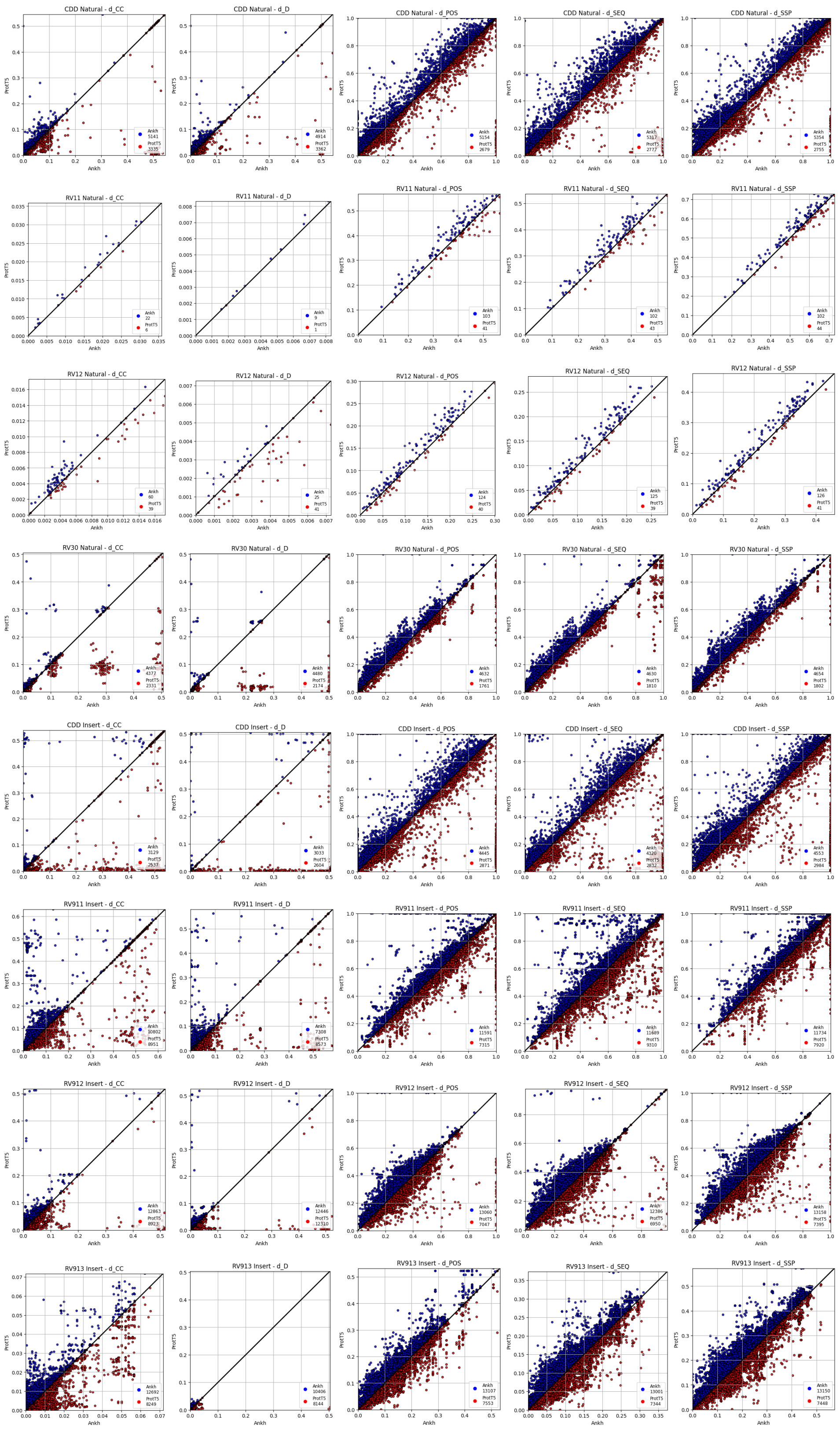


Supplementary Figure 1: Ankh-score vs ProtT5-score on all datasets.

#### 6.1 CDD natural

Supplementary Table 35: Ankh-score vs ProtT5-score on CDD natural dataset.

| Domain | $d_{cc}$ | | | $d_d$ | | | $d_{pos}$ | | | $d_{seq}$ | | | $d_{ssp}$ | | |
|---|---|---|---|---|---|---|---|---|---|---|---|---|---|---|---|
| Average distances | | | | | | | | | | | | | | | |
| | Ankh | ProtT5 | P-value | Ankh | ProtT5 | P-value | Ankh | ProtT5 | P-value | Ankh | ProtT5 | P-value | Ankh | ProtT5 | P-value |
| cd00012 | 0.0446 | **0.0441** | 6.0E-08 | **0.0235** | 0.0242 | 5.4E-07 | **0.6738** | 0.6956 | 4.4E-26 | **0.5926** | 0.6100 | 3.9E-19 | **0.7964** | 0.8139 | 3.3E-27 |
| cd00083 | **0.0026** | 0.0063 | 1.6E-60 | **0.0006** | 0.0020 | 1.4E-57 | **0.0430** | 0.0869 | 1.8E-59 | **0.0333** | 0.0765 | 8.0E-63 | **0.0885** | 0.1559 | 2.0E-61 |
| cd00173 | 0.0335 | 0.0346 | 2.1E-02 | 0.0081 | 0.0082 | 1.2E-02 | 0.4884 | 0.4589 | 7.7E-02 | 0.4576 | 0.4310 | 5.3E-02 | 0.5958 | 0.5767 | 5.1E-02 |
| cd00637 | 0.0044 | **0.0041** | 2.8E-20 | 0.0010 | **0.0009** | 6.9E-35 | **0.1315** | 0.1316 | 3.1E-09 | **0.1055** | 0.1083 | 5.6E-10 | **0.2211** | 0.2239 | 2.1E-12 |
| cd01040 | **0.0037** | 0.0052 | 3.7E-159 | **0.0007** | 0.0010 | 2.8E-151 | **0.1006** | 0.1208 | 1.2E-146 | **0.0795** | 0.1005 | 1.6E-161 | **0.1741** | 0.2036 | 1.2E-158 |
| cd01068 | **0.0027** | 0.0028 | 6.7E-03 | 0.0006 | 0.0007 | 5.3E-02 | **0.1046** | 0.1093 | 2.1E-04 | **0.1028** | 0.1068 | 2.0E-03 | **0.1720** | 0.1778 | 8.0E-04 |
| cd10140 | **0.0596** | 0.0611 | 3.5E-08 | **0.0120** | 0.0126 | 2.5E-06 | **0.7260** | 0.7372 | 3.9E-03 | **0.6739** | 0.6907 | 1.9E-03 | **0.8359** | 0.8435 | 3.7E-03 |
| cd21116 | **0.0084** | 0.0101 | 7.9E-04 | **0.0029** | 0.0037 | 5.8E-03 | **0.2944** | 0.3255 | 3.4E-09 | **0.2594** | 0.2846 | 3.7E-07 | **0.4129** | 0.4562 | 9.5E-10 |
| Domains won | **5** | 2 | | **5** | 1 | | **7** | 0 | | **7** | 0 | | **7** | 0 | |
| Tests won | | | | | | | | | | | | | | | |
| | Ankh | ProtT5 | Equal | Ankh | ProtT5 | Equal | Ankh | ProtT5 | Equal | Ankh | ProtT5 | Equal | Ankh | ProtT5 | Equal |
| cd00012 | **989** | 778 | 3 | **976** | 790 | 4 | **1005** | 639 | 126 | **1013** | 709 | 48 | **1042** | 678 | 50 |
| cd00083 | **724** | 146 | 211 | **714** | 114 | 253 | **713** | 112 | 256 | **714** | 112 | 255 | **714** | 113 | 254 |
| cd00173 | 162 | 100 | 63 | 161 | 102 | 62 | 148 | 98 | 79 | 153 | 99 | 73 | 152 | 100 | 73 |
| cd00637 | 1320 | **1676** | 7 | 1255 | **1742** | 6 | **1599** | 1273 | 131 | **1611** | 1279 | 113 | **1654** | 1291 | 58 |
| cd01040 | **1464** | 327 | 100 | **1423** | 344 | 124 | **1334** | 316 | 241 | **1410** | 291 | 190 | **1406** | 306 | 179 |
| cd01068 | **84** | 47 | 40 | 80 | 47 | 44 | **83** | 38 | 50 | **80** | 43 | 48 | **81** | 42 | 48 |
| cd10140 | **358** | 239 | 33 | **345** | 251 | 34 | **220** | 190 | 220 | **290** | 230 | 110 | **258** | 211 | 161 |
| cd21116 | **202** | 122 | 1 | **201** | 121 | 3 | **200** | 111 | 14 | **199** | 113 | 13 | **199** | 114 | 12 |
| Total | **5141** | 3335 | 395 | **4914** | 3362 | 424 | **5154** | 2679 | 1038 | **5317** | 2777 | 777 | **5354** | 2755 | 762 |
| Tests won (%) | | | | | | | | | | | | | | | |
| cd00012 | **55.88** | 43.95 | 0.17 | **55.14** | 44.63 | 0.23 | **56.78** | 36.10 | 7.12 | **57.23** | 40.06 | 2.71 | **58.87** | 38.31 | 2.82 |
| cd00083 | **66.98** | 13.51 | 19.52 | **66.05** | 10.55 | 23.40 | **65.96** | 10.36 | 23.68 | **66.05** | 10.36 | 23.59 | **66.05** | 10.45 | 23.50 |
| cd00173 | 49.85 | 30.77 | 19.38 | 49.54 | 31.38 | 19.08 | 45.54 | 30.15 | 24.31 | 47.08 | 30.46 | 22.46 | 46.77 | 30.77 | 22.46 |
| cd00637 | 43.96 | **55.81** | 0.23 | 41.79 | **58.01** | 0.20 | **53.25** | 42.39 | 4.36 | **53.65** | 42.59 | 3.76 | **55.08** | 42.99 | 1.93 |
| cd01040 | **77.42** | 17.29 | 5.29 | **75.25** | 18.19 | 6.56 | **70.54** | 16.71 | 12.74 | **74.56** | 15.39 | 10.05 | **74.35** | 16.18 | 9.47 |
| cd01068 | **49.12** | 27.49 | 23.39 | 46.78 | 27.49 | 25.73 | **48.54** | 22.22 | 29.24 | **46.78** | 25.15 | 28.07 | **47.37** | 24.56 | 28.07 |
| cd10140 | **56.83** | 37.94 | 5.24 | **54.76** | 39.84 | 5.40 | **34.92** | 30.16 | 34.92 | **46.03** | 36.51 | 17.46 | **40.95** | 33.49 | 25.56 |
| cd21116 | **62.15** | 37.54 | 0.31 | **61.85** | 37.23 | 0.92 | **61.54** | 34.15 | 4.31 | **61.23** | 34.77 | 4.00 | **61.23** | 35.08 | 3.69 |
| Average | **58.90** | 33.36 | 7.74 | **59.14** | 34.74 | 6.12 | **55.93** | 27.44 | 16.63 | **57.93** | 29.26 | 12.81 | **57.70** | 28.72 | 13.58 |

#### 6.2 BALiBASE RV11

Supplementary Table 36: Ankh-score vs ProtT5-score on BALiBASE RV11 dataset.

| Domain | $d_{cc}$ | | | $d_d$ | | | $d_{pos}$ | | | $d_{seq}$ | | | $d_{ssp}$ | | |
|---|---|---|---|---|---|---|---|---|---|---|---|---|---|---|---|
| Average distances | | | | | | | | | | | | | | | |
| | Ankh | ProtT5 | P-value | Ankh | ProtT5 | P-value | Ankh | ProtT5 | P-value | Ankh | ProtT5 | P-value | Ankh | ProtT5 | P-value |
| BBS11009 | 0.0207 | 0.0241 | 5.0E-01 | 0.0103 | 0.0105 | 2.2E-01 | 0.2317 | 0.2462 | 2.7E-01 | 0.2134 | 0.2344 | 2.2E-01 | 0.3654 | 0.3839 | 2.2E-01 |
| BBS11010 | 0.0232 | 0.0262 | 3.1E-01 | 0.0039 | 0.0046 | 9.4E-02 | 0.2844 | 0.3217 | 3.1E-01 | 0.2093 | 0.2351 | 3.1E-01 | 0.4258 | 0.4551 | 3.1E-01 |
| BBS11016 | 0.0111 | 0.0115 | 6.0E-02 | 0.0041 | 0.0044 | 1.3E-01 | 0.3578 | 0.3730 | 3.2E-02 | 0.3195 | 0.3322 | 3.4E-02 | 0.5197 | 0.5308 | 4.5E-02 |
| BBS11018 | 0.0190 | 0.0191 | 8.3E-01 | 0.0096 | 0.0090 | 1.2E-01 | **0.4264** | 0.4387 | 5.2E-03 | **0.3623** | 0.3707 | 3.4E-03 | **0.5856** | 0.5964 | 9.0E-03 |
| BBS11024 | 0.0244 | 0.0276 | 4.4E-01 | 0.0069 | 0.0136 | 6.9E-01 | 0.4017 | 0.4287 | 1.6E-01 | 0.3226 | 0.3596 | 2.2E-01 | 0.5443 | 0.5710 | 1.6E-01 |
| BBS11026 | 0.0873 | 0.0873 | 3.3E-01 | 0.0298 | 0.0293 | 4.1E-01 | 0.6577 | 0.6997 | 2.6E-02 | 0.6351 | 0.6757 | 3.1E-02 | 0.7841 | 0.8114 | 4.1E-02 |
| BBS11034 | 0.0196 | 0.0181 | 3.2E-02 | 0.0061 | 0.0061 | 8.1E-01 | **0.3908** | 0.4155 | 2.7E-04 | **0.3499** | 0.3767 | 3.4E-04 | **0.5500** | 0.5754 | 5.4E-04 |
| BBS11037 | 0.0229 | 0.0223 | 7.7E-01 | **0.0043** | 0.0045 | 3.9E-03 | 0.4064 | 0.4147 | 3.2E-01 | 0.3575 | 0.3695 | 1.9E-01 | 0.5548 | 0.5779 | 1.6E-01 |
| BBS11038 | **0.0167** | 0.0176 | 4.1E-03 | 0.0032 | 0.0032 | 6.0E-02 | **0.2270** | 0.2462 | 1.6E-03 | **0.1905** | 0.2074 | 1.6E-03 | **0.3564** | 0.3777 | 1.4E-03 |
| Domains won | **1** | 0 | | **1** | 0 | | **3** | 0 | | **3** | 0 | | **3** | 0 | |
| Tests won | | | | | | | | | | | | | | | |
| | Ankh | ProtT5 | Equal | Ankh | ProtT5 | Equal | Ankh | ProtT5 | Equal | Ankh | ProtT5 | Equal | Ankh | ProtT5 | Equal |
| BBS11009 | 4 | 1 | 1 | 4 | 1 | 1 | 3 | 1 | 2 | 4 | 1 | 1 | 4 | 1 | 1 |
| BBS11010 | 4 | 2 | 0 | 5 | 1 | 0 | 4 | 2 | 0 | 4 | 2 | 0 | 4 | 2 | 0 |
| BBS11016 | 18 | 10 | 0 | 18 | 10 | 0 | 19 | 8 | 1 | 19 | 8 | 1 | 19 | 9 | 0 |
| BBS11018 | 46 | 45 | 0 | 40 | 51 | 0 | **58** | 30 | 3 | **56** | 34 | 1 | **57** | 34 | 0 |
| BBS11024 | 4 | 2 | 0 | 3 | 3 | 0 | 5 | 1 | 0 | 5 | 1 | 0 | 5 | 1 | 0 |
| BBS11026 | 11 | 5 | 5 | 10 | 6 | 5 | 9 | 2 | 10 | 10 | 2 | 9 | 9 | 3 | 9 |
| BBS11034 | 21 | 7 | 0 | 13 | 15 | 0 | **23** | 5 | 0 | **23** | 4 | 1 | **22** | 5 | 1 |
| BBS11037 | 7 | 3 | 0 | **9** | 1 | 0 | 6 | 4 | 0 | 7 | 3 | 0 | 7 | 3 | 0 |
| BBS11038 | **22** | 6 | 0 | 21 | 7 | 0 | **22** | 6 | 0 | **23** | 5 | 0 | **23** | 5 | 0 |
| Total | **22** | 6 | 0 | **9** | 1 | 0 | **103** | 41 | 3 | **102** | 43 | 2 | **102** | 44 | 1 |
| Tests won (%) | | | | | | | | | | | | | | | |
| BBS11009 | 66.67 | 16.67 | 16.67 | 66.67 | 16.67 | 16.67 | 50.00 | 16.67 | 33.33 | 66.67 | 16.67 | 16.67 | 66.67 | 16.67 | 16.67 |
| BBS11010 | 66.67 | 33.33 | 0.00 | 83.33 | 16.67 | 0.00 | 66.67 | 33.33 | 0.00 | 66.67 | 33.33 | 0.00 | 66.67 | 33.33 | 0.00 |
| BBS11016 | 64.29 | 35.71 | 0.00 | 64.29 | 35.71 | 0.00 | 67.86 | 28.57 | 3.57 | 67.86 | 28.57 | 3.57 | 67.86 | 32.14 | 0.00 |
| BBS11018 | 50.55 | 49.45 | 0.00 | 43.96 | 56.04 | 0.00 | **63.74** | 32.97 | 3.30 | **61.54** | 37.36 | 1.10 | **62.64** | 37.36 | 0.00 |
| BBS11024 | 66.67 | 33.33 | 0.00 | 50.00 | 50.00 | 0.00 | 83.33 | 16.67 | 0.00 | 83.33 | 16.67 | 0.00 | 83.33 | 16.67 | 0.00 |
| BBS11026 | 52.38 | 23.81 | 23.81 | 47.62 | 28.57 | 23.81 | 42.86 | 9.52 | 47.62 | 47.62 | 9.52 | 42.86 | 42.86 | 14.29 | 42.86 |
| BBS11034 | 75.00 | 25.00 | 0.00 | 46.43 | 53.57 | 0.00 | **82.14** | 17.86 | 0.00 | **82.14** | 14.29 | 3.57 | **78.57** | 17.86 | 3.57 |
| BBS11037 | 70.00 | 30.00 | 0.00 | **90.00** | 10.00 | 0.00 | 60.00 | 40.00 | 0.00 | 70.00 | 30.00 | 0.00 | 70.00 | 30.00 | 0.00 |
| BBS11038 | **78.57** | 21.43 | 0.00 | 75.00 | 25.00 | 0.00 | **78.57** | 21.43 | 0.00 | **82.14** | 17.86 | 0.00 | **82.14** | 17.86 | 0.00 |
| Average | **78.57** | 21.43 | 0.00 | **90.00** | 10.00 | 0.00 | **74.82** | 24.08 | 1.10 | **75.27** | 23.17 | 1.56 | **74.45** | 24.36 | 1.19 |

#### 6.3 BALiBASE RV12

Supplementary Table 37: Ankh-score vs ProtT5-score on BALiBASE RV12 dataset.

| Domain | $d_{cc}$ | | | $d_d$ | | | $d_{pos}$ | | | $d_{seq}$ | | | $d_{ssp}$ | | |
|---|---|---|---|---|---|---|---|---|---|---|---|---|---|---|---|
| Average distances | | | | | | | | | | | | | | | |
| | Ankh | ProtT5 | P-value | Ankh | ProtT5 | P-value | Ankh | ProtT5 | P-value | Ankh | ProtT5 | P-value | Ankh | ProtT5 | P-value |
| BBS11009 | 0.0011 | 0.0011 | 1.9E-02 | 0.0003 | 0.0003 | 1.3E-02 | **0.0674** | 0.0718 | 1.1E-03 | **0.0635** | 0.0672 | 2.5E-03 | **0.1263** | 0.1336 | 4.6E-03 |
| BBS11026 | 0.0068 | 0.0069 | 7.8E-01 | 0.0031 | **0.0028** | 9.1E-03 | 0.3188 | 0.3264 | 6.8E-01 | 0.3062 | 0.3179 | 5.6E-01 | 0.4780 | 0.4897 | 8.5E-01 |
| BBS11034 | 0.0018 | 0.0025 | 3.0E-02 | 0.0002 | 0.0003 | 4.8E-02 | **0.0721** | 0.0889 | 2.3E-03 | **0.0701** | 0.0881 | 1.9E-03 | **0.1412** | 0.1701 | 1.9E-03 |
| BBS11037 | 0.0129 | **0.0118** | 2.9E-03 | 0.0032 | 0.0032 | 7.6E-01 | 0.2973 | 0.2983 | 6.1E-01 | 0.2360 | 0.2388 | 1.0E+00 | 0.4392 | 0.4380 | 1.0E+00 |
| BBS11038 | **0.0038** | 0.0043 | 2.8E-06 | 0.0022 | 0.0021 | 4.8E-01 | **0.1880** | 0.2059 | 5.8E-10 | **0.1726** | 0.1880 | 1.5E-09 | **0.3136** | 0.3372 | 9.0E-10 |
| Domains won | **1** | **1** | | 0 | **1** | | **3** | 0 | | **3** | 0 | | **3** | 0 | |
| Tests won | | | | | | | | | | | | | | | |
| | Ankh | ProtT5 | Equal | Ankh | ProtT5 | Equal | Ankh | ProtT5 | Equal | Ankh | ProtT5 | Equal | Ankh | ProtT5 | Equal |
| BBS11009 | 48 | 29 | 1 | 46 | 31 | 1 | **50** | 23 | 5 | **51** | 24 | 3 | **50** | 26 | 2 |
| BBS11026 | 37 | 29 | 0 | 25 | **41** | 0 | 35 | 31 | 0 | 36 | 30 | 0 | 33 | 33 | 0 |
| BBS11034 | 11 | 3 | 1 | 11 | 3 | 1 | **12** | 2 | 1 | **12** | 1 | 2 | **12** | 2 | 1 |
| BBS11037 | 5 | **16** | 0 | 11 | 10 | 0 | 11 | 10 | 0 | 11 | 10 | 0 | 11 | 10 | 0 |
| BBS11038 | **55** | 23 | 0 | 39 | 39 | 0 | **62** | 15 | 1 | **62** | 14 | 2 | **64** | 13 | 1 |
| Total | **60** | 39 | 0 | 25 | **41** | 0 | **124** | 40 | 7 | **125** | 39 | 7 | **126** | 41 | 4 |
| Tests won (%) | | | | | | | | | | | | | | | |
| BBS11009 | 61.54 | 37.18 | 1.28 | 58.97 | 39.74 | 1.28 | **64.10** | 29.49 | 6.41 | **65.38** | 30.77 | 3.85 | **64.10** | 33.33 | 2.56 |
| BBS11026 | 56.06 | 43.94 | 0.00 | 37.88 | **62.12** | 0.00 | 53.03 | 46.97 | 0.00 | 54.55 | 45.45 | 0.00 | 50.00 | 50.00 | 0.00 |
| BBS11034 | 73.33 | 20.00 | 6.67 | 73.33 | 20.00 | 6.67 | **80.00** | 13.33 | 6.67 | **80.00** | 6.67 | 13.33 | **80.00** | 13.33 | 6.67 |
| BBS11037 | 23.81 | **76.19** | 0.00 | 52.38 | 47.62 | 0.00 | 52.38 | 47.62 | 0.00 | 52.38 | 47.62 | 0.00 | 52.38 | 47.62 | 0.00 |
| BBS11038 | **70.51** | 29.49 | 0.00 | 50.00 | 50.00 | 0.00 | **79.49** | 19.23 | 1.28 | **79.49** | 17.95 | 2.56 | **82.05** | 16.67 | 1.28 |
| Average | 47.16 | **52.84** | 0.00 | 37.88 | **62.12** | 0.00 | **74.53** | 20.68 | 4.79 | **74.96** | 18.46 | 6.58 | **75.38** | 21.11 | 3.50 |

#### 6.4 BALiBASE RV30

Supplementary Table 38: Ankh-score vs ProtT5-score on BALiBASE RV30 dataset.

| Domain | $d_{cc}$ | | | $d_d$ | | | $d_{pos}$ | | | $d_{seq}$ | | | $d_{ssp}$ | | |
|---|---|---|---|---|---|---|---|---|---|---|---|---|---|---|---|
| Average distances | | | | | | | | | | | | | | | |
| | Ankh | ProtT5 | P-value | Ankh | ProtT5 | P-value | Ankh | ProtT5 | P-value | Ankh | ProtT5 | P-value | Ankh | ProtT5 | P-value |
| BBS30001 | **0.0157** | 0.0163 | 1.3E-25 | **0.0010** | 0.0013 | 9.9E-98 | **0.1513** | 0.1784 | 1.2E-87 | **0.1453** | 0.1739 | 5.4E-91 | **0.2360** | 0.2677 | 9.2E-89 |
| BBS30004 | **0.0019** | 0.0020 | 7.0E-17 | **0.0005** | 0.0005 | 8.0E-20 | **0.1171** | 0.1287 | 1.9E-43 | **0.1042** | 0.1114 | 1.4E-28 | **0.1935** | 0.2012 | 5.2E-24 |
| BBS30005 | **0.0078** | 0.0096 | 2.8E-98 | **0.0005** | 0.0008 | 6.8E-104 | **0.0817** | 0.1040 | 3.4E-103 | **0.0605** | 0.0803 | 1.4E-116 | **0.1422** | 0.1687 | 3.5E-114 |
| BBS30006 | **0.0566** | 0.0611 | 3.7E-11 | **0.0224** | 0.0246 | 1.8E-09 | **0.3454** | 0.3824 | 5.5E-15 | **0.3159** | 0.3524 | 1.9E-17 | **0.4445** | 0.4891 | 2.6E-18 |
| BBS30008 | **0.0200** | 0.0207 | 9.3E-09 | **0.0086** | 0.0089 | 3.3E-12 | **0.3952** | 0.4069 | 1.6E-36 | **0.3591** | 0.3666 | 3.8E-26 | **0.5062** | 0.5148 | 1.5E-38 |
| BBS30009 | 0.0694 | **0.0457** | 1.1E-32 | 0.0074 | **0.0070** | 3.9E-34 | 0.4691 | **0.4273** | 1.9E-19 | 0.4563 | **0.4180** | 3.1E-19 | 0.5083 | **0.4802** | 2.1E-21 |
| BBS30010 | **0.0013** | 0.0013 | 1.3E-15 | **0.0003** | 0.0004 | 4.7E-24 | **0.0845** | 0.0860 | 1.6E-21 | **0.0731** | 0.0783 | 1.0E-29 | **0.1404** | 0.1499 | 1.6E-26 |
| BBS30013 | 0.0135 | **0.0131** | 1.2E-03 | 0.0046 | **0.0043** | 1.8E-22 | **0.3144** | 0.3186 | 2.9E-08 | **0.2862** | 0.2880 | 2.7E-03 | **0.4238** | 0.4282 | 4.3E-08 |
| BBS30016 | 0.1917 | 0.1790 | 2.7E-02 | 0.1741 | 0.1585 | 2.1E-01 | 0.4616 | 0.4428 | 7.5E-02 | 0.4311 | 0.4189 | 7.7E-02 | 0.5044 | 0.4989 | 7.8E-02 |
| BBS30030 | **0.0043** | 0.0055 | 5.8E-76 | **0.0043** | 0.0043 | 8.3E-63 | **0.1262** | 0.1511 | 5.3E-92 | **0.1113** | 0.1357 | 5.2E-96 | **0.2148** | 0.2480 | 3.5E-95 |
| Domains won | **7** | 2 | | **7** | 2 | | **8** | 1 | | **8** | 1 | | **8** | 1 | |
| Tests won | | | | | | | | | | | | | | | |
| | Ankh | ProtT5 | Equal | Ankh | ProtT5 | Equal | Ankh | ProtT5 | Equal | Ankh | ProtT5 | Equal | Ankh | ProtT5 | Equal |
| BBS30001 | **546** | 219 | 235 | **679** | 81 | 240 | **644** | 103 | 253 | **657** | 102 | 241 | **657** | 103 | 240 |
| BBS30004 | **589** | 346 | 65 | **569** | 346 | 85 | **598** | 277 | 125 | **562** | 297 | 141 | **561** | 311 | 128 |
| BBS30005 | **758** | 135 | 107 | **784** | 99 | 117 | **746** | 110 | 144 | **769** | 89 | 142 | **768** | 95 | 137 |
| BBS30006 | **119** | 24 | 10 | **115** | 28 | 10 | **122** | 15 | 16 | **125** | 15 | 13 | **126** | 14 | 13 |
| BBS30008 | **387** | 218 | 25 | **398** | 206 | 26 | **434** | 154 | 42 | **423** | 163 | 44 | **446** | 147 | 37 |
| BBS30009 | 162 | **347** | 194 | 158 | **349** | 196 | 173 | **253** | 277 | 190 | **269** | 244 | 170 | **257** | 276 |
| BBS30010 | **578** | 395 | 27 | **622** | 352 | 26 | **575** | 360 | 65 | **600** | 349 | 51 | **592** | 365 | 43 |
| BBS30013 | **476** | 472 | 52 | 400 | **536** | 64 | **566** | 354 | 80 | **523** | 397 | 80 | **555** | 372 | 73 |
| BBS30016 | 134 | 92 | 440 | 134 | 87 | 445 | 116 | 88 | 462 | 118 | 90 | 458 | 118 | 91 | 457 |
| BBS30030 | **762** | 175 | 63 | **755** | 177 | 68 | **774** | 135 | 91 | **781** | 129 | 90 | **779** | 138 | 83 |
| Total | **4377** | 2331 | 778 | **4480** | 2174 | 832 | **4632** | 1761 | 1093 | **4630** | 1810 | 1046 | **4654** | 1802 | 1030 |
| Tests won (%) | | | | | | | | | | | | | | | |
| BBS30001 | **54.60** | 21.90 | 23.50 | **67.90** | 8.10 | 24.00 | **64.40** | 10.30 | 25.30 | **65.70** | 10.20 | 24.10 | **65.70** | 10.30 | 24.00 |
| BBS30004 | **58.90** | 34.60 | 6.50 | **56.90** | 34.60 | 8.50 | **59.80** | 27.70 | 12.50 | **56.20** | 29.70 | 14.10 | **56.10** | 31.10 | 12.80 |
| BBS30005 | **75.80** | 13.50 | 10.70 | **78.40** | 9.90 | 11.70 | **74.60** | 11.00 | 14.40 | **76.90** | 8.90 | 14.20 | **76.80** | 9.50 | 13.70 |
| BBS30006 | **77.78** | 15.69 | 6.54 | **75.16** | 18.30 | 6.54 | **79.74** | 9.80 | 10.46 | **81.70** | 9.80 | 8.50 | **82.35** | 9.15 | 8.50 |
| BBS30008 | **61.43** | 34.60 | 3.97 | **63.17** | 32.70 | 4.13 | **68.89** | 24.44 | 6.67 | **67.14** | 25.87 | 6.98 | **70.79** | 23.33 | 5.87 |
| BBS30009 | 23.04 | **49.36** | 27.60 | 22.48 | **49.64** | 27.88 | 24.61 | **35.99** | 39.40 | 27.03 | **38.26** | 34.71 | 24.18 | **36.56** | 39.26 |
| BBS30010 | **57.80** | 39.50 | 2.70 | **62.20** | 35.20 | 2.60 | **57.50** | 36.00 | 6.50 | **60.00** | 34.90 | 5.10 | **59.20** | 36.50 | 4.30 |
| BBS30013 | **47.60** | 47.20 | 5.20 | 40.00 | **53.60** | 6.40 | **56.60** | 35.40 | 8.00 | **52.30** | 39.70 | 8.00 | **55.50** | 37.20 | 7.30 |
| BBS30016 | 20.12 | 13.81 | 66.07 | 20.12 | 13.06 | 66.82 | 17.42 | 13.21 | 69.37 | 17.72 | 13.51 | 68.77 | 17.72 | 13.66 | 68.62 |
| BBS30030 | **76.20** | 17.50 | 6.30 | **75.50** | 17.70 | 6.80 | **77.40** | 13.50 | 9.10 | **78.10** | 12.90 | 9.00 | **77.90** | 13.80 | 8.30 |
| Average | **59.24** | 30.43 | 10.33 | **60.19** | 28.86 | 10.95 | **62.62** | 22.68 | 14.70 | **62.79** | 23.36 | 13.85 | **63.17** | 23.05 | 13.78 |

#### 6.5 CDD insert

Supplementary Table 39: Ankh-score vs ProtT5-score on CDD insert dataset.

| Domain | $d_{cc}$ | | | $d_{d}$ | | | $d_{pos}$ | | | $d_{seq}$ | | | $d_{ssp}$ | | |
|---|---|---|---|---|---|---|---|---|---|---|---|---|---|---|---|
| Average distances | | | | | | | | | | | | | | | |
| | Ankh | ProtT5 | P-value | Ankh | ProtT5 | P-value | Ankh | ProtT5 | P-value | Ankh | ProtT5 | P-value | Ankh | ProtT5 | P-value |
| cd00012 | 0.0448 | 0.0444 | 1.4E-01 | 0.0228 | 0.0228 | 4.3E-02 | **0.6829** | 0.7018 | 7.3E-21 | **0.6001** | 0.6171 | 2.0E-18 | **0.8051** | 0.8169 | 7.8E-20 |
| cd00083 | 0.1640 | **0.0069** | 4.6E-05 | 0.1546 | **0.0023** | 1.2E-05 | 0.3986 | **0.0958** | 2.4E-05 | 0.3775 | **0.0784** | 1.1E-04 | 0.4310 | **0.1610** | 7.0E-05 |
| cd00173 | 0.0354 | 0.0369 | 6.1E-01 | 0.0082 | 0.0080 | 7.7E-01 | 0.4242 | 0.3446 | 8.2E-01 | 0.4028 | 0.3305 | 7.3E-01 | 0.5465 | 0.5029 | 6.7E-01 |
| cd00637 | 0.0045 | **0.0041** | 2.1E-31 | 0.0010 | **0.0009** | 1.1E-39 | **0.1313** | 0.1322 | 2.0E-07 | **0.1063** | 0.1088 | 5.7E-07 | **0.2217** | 0.2246 | 2.8E-09 |
| cd01040 | **0.0041** | 0.0055 | 1.7E-71 | **0.0009** | 0.0012 | 5.4E-54 | **0.1162** | 0.1343 | 4.3E-64 | **0.0966** | 0.1111 | 3.1E-62 | **0.2028** | 0.2246 | 2.6E-58 |
| cd01068 | 0.0150 | 0.0035 | 5.1E-01 | 0.0010 | 0.0007 | 3.2E-01 | 0.1275 | 0.1184 | 2.6E-01 | 0.1257 | 0.1160 | 2.3E-01 | 0.2064 | 0.1921 | 2.5E-01 |
| cd10140 | 0.0657 | 0.0650 | 3.1E-01 | 0.0170 | 0.0165 | 1.8E-02 | 0.7565 | 0.7562 | 9.1E-01 | 0.7089 | 0.7163 | 3.9E-01 | 0.8518 | 0.8552 | 8.5E-01 |
| cd21116 | 0.0140 | 0.0148 | 5.9E-01 | 0.0039 | 0.0039 | 7.9E-01 | **0.3478** | 0.3672 | 1.8E-03 | 0.3200 | 0.3369 | 1.3E-02 | **0.4810** | 0.5060 | 1.7E-03 |
| Domains won | 1 | **2** | | 2 | **6** | | **4** | **4** | | **5** | 3 | | **5** | 3 | |
| Tests won | | | | | | | | | | | | | | | |
| | Ankh | ProtT5 | Equal | Ankh | ProtT5 | Equal | Ankh | ProtT5 | Equal | Ankh | ProtT5 | Equal | Ankh | ProtT5 | Equal |
| cd00012 | 940 | 829 | 1 | 922 | 847 | 1 | **990** | 667 | 113 | **1015** | 711 | 44 | **1019** | 700 | 51 |
| cd00083 | **519** | 349 | 213 | **513** | 336 | 232 | **490** | 313 | 278 | **492** | 317 | 272 | **490** | 315 | 276 |
| cd00173 | 179 | 139 | 7 | 186 | 130 | 9 | 184 | 118 | 23 | 186 | 115 | 24 | 183 | 119 | 23 |
| cd00637 | 1252 | **1745** | 6 | 1223 | **1779** | 1 | **1545** | 1306 | 152 | **1532** | 1348 | 123 | **1588** | 1355 | 60 |
| cd01040 | **1358** | 443 | 90 | **1297** | 489 | 105 | **1242** | 450 | 199 | **1281** | 456 | 154 | **1274** | 475 | 142 |
| cd01068 | 72 | 63 | 36 | 69 | 63 | 39 | 65 | 57 | 49 | 66 | 60 | 45 | 67 | 60 | 44 |
| cd10140 | 284 | 312 | 34 | 271 | 325 | 34 | 195 | 207 | 228 | 262 | 253 | 115 | 225 | 233 | 172 |
| cd21116 | 176 | 148 | 1 | 174 | 151 | 0 | **178** | 135 | 12 | 180 | 140 | 5 | **182** | 139 | 4 |
| Total | **3129** | 2537 | 309 | **3033** | 2604 | 338 | **4445** | 2871 | 754 | **4320** | 2832 | 593 | **4553** | 2984 | 533 |
| Tests won (%) | | | | | | | | | | | | | | | |
| cd00012 | 53.11 | 46.84 | 0.06 | 52.09 | 47.85 | 0.06 | **55.93** | 37.68 | 6.38 | **57.34** | 40.17 | 2.49 | **57.57** | 39.55 | 2.88 |
| cd00083 | **48.01** | 32.28 | 19.70 | **47.46** | 31.08 | 21.46 | **45.33** | 28.95 | 25.72 | **45.51** | 29.32 | 25.16 | **45.33** | 29.14 | 25.53 |
| cd00173 | 55.08 | 42.77 | 2.15 | 57.23 | 40.00 | 2.77 | 56.62 | 36.31 | 7.08 | 57.23 | 35.38 | 7.38 | 56.31 | 36.62 | 7.08 |
| cd00637 | 41.69 | **58.11** | 0.20 | 40.73 | **59.24** | 0.03 | **51.45** | 43.49 | 5.06 | **51.02** | 44.89 | 4.10 | **52.88** | 45.12 | 2.00 |
| cd01040 | **71.81** | 23.43 | 4.76 | **68.59** | 25.86 | 5.55 | **65.68** | 23.80 | 10.52 | **67.74** | 24.11 | 8.14 | **67.37** | 25.12 | 7.51 |
| cd01068 | 42.11 | 36.84 | 21.05 | 40.35 | 36.84 | 22.81 | 38.01 | 33.33 | 28.65 | 38.60 | 35.09 | 26.32 | 39.18 | 35.09 | 25.73 |
| cd10140 | 45.08 | 49.52 | 5.40 | 43.02 | 51.59 | 5.40 | 30.95 | 32.86 | 36.19 | 41.59 | 40.16 | 18.25 | 35.71 | 36.98 | 27.30 |
| cd21116 | 54.15 | 45.54 | 0.31 | 53.54 | 46.46 | 0.00 | **54.77** | 41.54 | 3.69 | 55.38 | 43.08 | 1.54 | **56.00** | 42.77 | 1.23 |
| Average | **53.84** | 37.94 | 8.22 | **52.26** | 38.73 | 9.02 | **54.63** | 35.09 | 10.28 | **55.40** | 34.62 | 9.97 | **55.83** | 36.34 | 7.83 |

#### 6.6 BALiBASE RV911

Supplementary Table 40: Ankh-score vs ProtT5-score on BALiBASE RV911 dataset.

**Average distances**

| Domain | $d_{cc}$ | | | $d_d$ | | | $d_{pos}$ | | | $d_{seq}$ | | | $d_{ssp}$ | | |
|---|---|---|---|---|---|---|---|---|---|---|---|---|---|---|---|
| | Ankh | ProtT5 | P-value | Ankh | ProtT5 | P-value | Ankh | ProtT5 | P-value | Ankh | ProtT5 | P-value | Ankh | ProtT5 | P-value |
| BOX001 | 0.0160 | **0.0153** | 1.9E-12 | 0.0076 | **0.0060** | 8.5E-26 | 0.5400 | 0.5389 | 1.2E-01 | 0.4800 | **0.4799** | 6.8E-04 | 0.6890 | 0.6869 | 2.1E-01 |
| BOX022 | **0.0393** | 0.0406 | 4.8E-03 | 0.0129 | 0.0132 | 6.1E-01 | **0.4944** | 0.4992 | 8.5E-04 | 0.5363 | 0.4809 | 1.2E-01 | 0.7076 | **0.7069** | 3.8E-03 |
| BOX032 | **0.0291** | 0.0294 | 2.3E-03 | 0.0120 | **0.0118** | 1.2E-20 | 0.5670 | 0.5678 | 2.3E-01 | 0.5513 | 0.5506 | 1.4E-01 | 0.7193 | 0.7197 | 6.1E-01 |
| BOX034 | 0.0145 | 0.0168 | 1.3E-02 | 0.0118 | **0.0113** | 3.0E-09 | **0.4642** | 0.4746 | 1.8E-28 | 0.4294 | **0.4271** | 2.0E-05 | **0.6301** | 0.6306 | 5.5E-07 |
| BOX035 | **0.0324** | 0.0364 | 2.0E-18 | 0.0482 | 0.0443 | 5.5E-01 | **0.6624** | 0.6687 | 9.7E-50 | 0.5767 | **0.5724** | 9.5E-07 | **0.7881** | 0.7919 | 3.5E-42 |
| BOX046 | 0.0568 | 0.0559 | 4.5E-01 | 0.0407 | 0.0402 | 3.0E-02 | 0.7165 | **0.7119** | 1.5E-12 | 0.6867 | **0.6834** | 4.4E-18 | 0.8109 | **0.8089** | 1.0E-14 |
| BOX060 | **0.1131** | 0.1161 | 2.3E-05 | 0.0752 | 0.0765 | 4.6E-02 | **0.7821** | 0.7929 | 3.0E-50 | **0.7055** | 0.7149 | 5.7E-10 | **0.8752** | 0.8820 | 1.7E-51 |
| BOX063 | **0.0082** | 0.0083 | 2.9E-10 | 0.0025 | **0.0024** | 6.0E-14 | **0.3600** | 0.3709 | 8.2E-55 | **0.3256** | 0.3370 | 7.1E-55 | **0.5161** | 0.5269 | 8.5E-46 |
| BOX076 | 0.0835 | 0.0833 | 1.8E-01 | 0.0653 | 0.0653 | 9.8E-02 | **0.7244** | 0.7295 | 2.8E-03 | **0.6282** | 0.6334 | 2.4E-05 | **0.8261** | 0.8305 | 9.4E-09 |
| BOX096 | 0.0297 | 0.0299 | 8.5E-01 | 0.0117 | **0.0106** | 1.6E-22 | **0.6267** | 0.6458 | 1.9E-77 | **0.5729** | 0.5876 | 7.8E-61 | **0.7623** | 0.7748 | 9.8E-77 |
| BOX115 | 0.0568 | 0.0582 | 1.2E-01 | **0.0375** | 0.0389 | 3.6E-04 | 0.8177 | 0.8167 | 4.7E-02 | 0.7787 | 0.7781 | 2.3E-01 | 0.8964 | 0.8966 | 3.6E-01 |
| BOX121 | **0.0223** | 0.0236 | 4.3E-14 | 0.0211 | 0.0210 | 8.9E-02 | **0.5563** | 0.5616 | 2.5E-25 | **0.5126** | 0.5187 | 1.0E-20 | **0.6936** | 0.6994 | 3.4E-35 |
| BOX122 | **0.0191** | 0.0195 | 2.2E-33 | 0.0048 | 0.0048 | 1.0E-01 | **0.3679** | 0.3719 | 2.3E-08 | **0.3615** | 0.3662 | 7.9E-09 | **0.5333** | 0.5370 | 3.3E-07 |
| BOX123 | **0.0469** | 0.0480 | 4.8E-04 | 0.0217 | **0.0217** | 9.6E-03 | 0.7220 | **0.6727** | 4.1E-58 | 0.6242 | **0.6097** | 5.2E-58 | 0.8319 | **0.8130** | 4.0E-65 |
| BOX142 | 0.0370 | 0.0402 | 3.0E-01 | **0.0231** | 0.0267 | 9.5E-22 | 0.6550 | **0.6543** | 1.2E-05 | 0.5862 | **0.5837** | 4.3E-03 | **0.7626** | 0.7650 | 6.6E-07 |
| BOX172 | 0.0890 | 0.0910 | 8.9E-02 | 0.1022 | 0.1113 | 4.8E-02 | **0.8343** | 0.8378 | 5.1E-04 | **0.6924** | 0.6973 | 9.1E-04 | **0.8929** | 0.8953 | 7.3E-04 |
| BOX175 | 0.0575 | **0.0537** | 5.3E-04 | 0.0273 | **0.0245** | 4.2E-04 | 0.6981 | **0.6930** | 4.6E-05 | 0.6615 | **0.6526** | 3.0E-12 | 0.8167 | **0.8133** | 8.3E-06 |
| BOX177 | 0.0353 | **0.0314** | 6.9E-09 | 0.0153 | 0.0148 | 2.2E-01 | 0.6708 | 0.6712 | 2.5E-02 | 0.6328 | 0.6296 | 6.3E-01 | 0.7954 | 0.7987 | 6.9E-02 |
| BOX180 | **0.0094** | 0.0097 | 4.9E-11 | **0.0056** | 0.0058 | 9.4E-11 | **0.2992** | 0.3061 | 3.0E-11 | **0.2771** | 0.2839 | 2.5E-12 | **0.4465** | 0.4536 | 9.4E-11 |
| BOX181 | **0.0683** | 0.0693 | 1.4E-13 | **0.0190** | 0.0196 | 2.3E-17 | **0.5716** | 0.5832 | 6.9E-13 | **0.5591** | 0.5717 | 8.6E-15 | **0.7250** | 0.7370 | 1.0E-19 |
| BOX192 | 0.0587 | **0.0569** | 6.8E-05 | 0.0422 | **0.0392** | 1.0E-09 | 0.6887 | 0.6851 | 8.6E-01 | 0.6005 | **0.5923** | 4.1E-03 | 0.8049 | 0.8015 | 3.7E-01 |
| BOX212 | 0.0562 | 0.0560 | 1.4E-01 | 0.0240 | 0.0257 | 4.9E-01 | 0.7332 | 0.7309 | 3.9E-01 | 0.6222 | 0.6186 | 4.0E-02 | 0.8346 | 0.8316 | 4.2E-01 |
| BOX214 | **0.0374** | 0.0383 | 2.9E-08 | 0.0392 | 0.0405 | 9.2E-01 | **0.5138** | 0.5199 | 3.4E-16 | **0.4244** | 0.4313 | 7.1E-15 | **0.6650** | 0.6983 | 1.1E-20 |
| BOX222 | **0.0232** | 0.0237 | 2.6E-07 | 0.0212 | 0.0215 | 6.6E-01 | **0.6460** | 0.6498 | 1.3E-26 | 0.4990 | **0.4946** | 7.5E-12 | 0.7655 | **0.7634** | 9.8E-29 |
| BOX240 | **0.0530** | 0.0532 | 5.8E-03 | 0.0204 | **0.0193** | 2.6E-04 | 0.6969 | 0.6847 | 3.2E-01 | 0.6455 | 0.6350 | 4.5E-01 | 0.8014 | 0.8016 | 8.7E-01 |
| BOX246 | 0.0387 | 0.0380 | 3.9E-01 | 0.0189 | 0.0196 | 1.2E-01 | 0.5935 | 0.5933 | 7.3E-01 | 0.5589 | **0.5535** | 6.2E-07 | 0.7349 | **0.7317** | 9.0E-03 |
| BOX258 | 0.0233 | **0.0225** | 8.6E-03 | 0.0181 | **0.0143** | 2.6E-26 | **0.4435** | 0.4462 | 1.5E-04 | **0.4168** | 0.4169 | 9.2E-15 | **0.6096** | 0.6188 | 3.1E-11 |
| BOX270 | **0.0451** | 0.0470 | 2.0E-05 | **0.0225** | 0.0239 | 1.9E-03 | 0.6211 | **0.6156** | 5.0E-15 | 0.5833 | 0.5775 | 4.3E-02 | 0.7564 | **0.7533** | 4.5E-09 |
| BOX284 | **0.0378** | 0.0382 | 4.2E-14 | 0.0190 | **0.0179** | 3.4E-23 | 0.5848 | **0.5783** | 3.4E-05 | 0.5032 | **0.4972** | 3.7E-03 | 0.7064 | 0.7043 | 1.4E-02 |
| Domains won | **15** | 5 | | 5 | **11** | | **15** | 6 | | **11** | **11** | | **14** | 7 | |

**Tests won**

| Domain | $d_{cc}$ Ankh | ProtT5 | Equal | $d_d$ Ankh | ProtT5 | Equal | $d_{pos}$ Ankh | ProtT5 | Equal | $d_{seq}$ Ankh | ProtT5 | Equal | $d_{ssp}$ Ankh | ProtT5 | Equal |
|---|---|---|---|---|---|---|---|---|---|---|---|---|---|---|---|
| BOX001 | 415 | **585** | 0 | 373 | **627** | 0 | 479 | 515 | 6 | 468 | **524** | 8 | 484 | 515 | 1 |
| BOX022 | **492** | 389 | 119 | 445 | 437 | 118 | **444** | 366 | 190 | 427 | 418 | 155 | **453** | 387 | 160 |
| BOX032 | **504** | 496 | 0 | 348 | **652** | 0 | 536 | 443 | 21 | 479 | 500 | 21 | 535 | 464 | 1 |
| BOX034 | 500 | 499 | 1 | 382 | **617** | 1 | **685** | 255 | 60 | **545** | 401 | 54 | **556** | 406 | 38 |
| BOX035 | **634** | 366 | 0 | 486 | 514 | 0 | **720** | 239 | 41 | **628** | 361 | 11 | **721** | 277 | 2 |
| BOX046 | 522 | 466 | 12 | 483 | 505 | 12 | **517** | 215 | 268 | **603** | 317 | 80 | **527** | 216 | 257 |
| BOX060 | **583** | 416 | 1 | 534 | 465 | 1 | **709** | 177 | 114 | **596** | 323 | 81 | **713** | 253 | 34 |
| BOX063 | **643** | 357 | 0 | 426 | **574** | 0 | **693** | 303 | 4 | **677** | 314 | 9 | **682** | 315 | 3 |
| BOX076 | 467 | 500 | 33 | 445 | 520 | 35 | **471** | 346 | 183 | **524** | 380 | 96 | **518** | 336 | 146 |
| BOX096 | 534 | 466 | 0 | 389 | **611** | 0 | **800** | 192 | 8 | **769** | 220 | 11 | **808** | 191 | 1 |
| BOX115 | 514 | 486 | 0 | 450 | **550** | 0 | 357 | 329 | 314 | 495 | 488 | 17 | 501 | 489 | 10 |
| BOX121 | **637** | 363 | 0 | 450 | 550 | 0 | **696** | 270 | 34 | **676** | 313 | 11 | **699** | 298 | 3 |
| BOX122 | **660** | 312 | 28 | 464 | 508 | 28 | **542** | 401 | 57 | **531** | 415 | 54 | **540** | 409 | 51 |
| BOX123 | 421 | **579** | 0 | 481 | **519** | 0 | 361 | **580** | 59 | 338 | **648** | 14 | 365 | **634** | 1 |
| BOX142 | 437 | 508 | 55 | 290 | **655** | 55 | **486** | 423 | 91 | **488** | 431 | 81 | **491** | 427 | 82 |
| BOX172 | 517 | 483 | 0 | 429 | 571 | 0 | **324** | 272 | 404 | **503** | 420 | 77 | **389** | 318 | 293 |
| BOX175 | 421 | **579** | 0 | 443 | **557** | 0 | 318 | **545** | 137 | 335 | **608** | 57 | 361 | **589** | 50 |
| BOX177 | 418 | **579** | 3 | 492 | 508 | 0 | 532 | 432 | 36 | 512 | 460 | 28 | 525 | 461 | 14 |
| BOX180 | **608** | 389 | 3 | **644** | 353 | 3 | **585** | 358 | 57 | **600** | 358 | 42 | **606** | 366 | 28 |
| BOX181 | **603** | 393 | 4 | **662** | 334 | 4 | **560** | 367 | 73 | **560** | 378 | 62 | **581** | 373 | 46 |
| BOX192 | 436 | **556** | 8 | 419 | **573** | 8 | 390 | 363 | 247 | 427 | **505** | 68 | 446 | 495 | 59 |
| BOX212 | 482 | 517 | 1 | 481 | 519 | 0 | 482 | 440 | 78 | 481 | 495 | 24 | 494 | 438 | 68 |
| BOX214 | **580** | 390 | 30 | 481 | 491 | 28 | **564** | 307 | 129 | **580** | 341 | 79 | **592** | 322 | 86 |
| BOX222 | **622** | 377 | 1 | 536 | 463 | 1 | **680** | 293 | 27 | **620** | 366 | 14 | **707** | 290 | 3 |
| BOX240 | 464 | **500** | 36 | 471 | **494** | 35 | 459 | 431 | 110 | 469 | 473 | 58 | 483 | 455 | 62 |
| BOX246 | 527 | 472 | 1 | 516 | 484 | 0 | 478 | 469 | 53 | 424 | **549** | 27 | 462 | **527** | 11 |
| BOX258 | 455 | **538** | 7 | 341 | **652** | 7 | 411 | **538** | 51 | 334 | **617** | 49 | 361 | **607** | 32 |
| BOX270 | **568** | 426 | 6 | **531** | 463 | 6 | **591** | 341 | 68 | 529 | 433 | 38 | **602** | 379 | 19 |
| BOX284 | **638** | 361 | 1 | **658** | 342 | 0 | 434 | **527** | 39 | 463 | **521** | 16 | 465 | 522 | 13 |
| Total | **10802** | 8951 | 247 | 7308 | **8573** | 119 | **11591** | 7315 | 2094 | **11689** | 9310 | 1001 | **11734** | 7920 | 1346 |

**Tests won (%)**

| Domain | $d_{cc}$ Ankh | ProtT5 | Equal | $d_d$ Ankh | ProtT5 | Equal | $d_{pos}$ Ankh | ProtT5 | Equal | $d_{seq}$ Ankh | ProtT5 | Equal | $d_{ssp}$ Ankh | ProtT5 | Equal |
|---|---|---|---|---|---|---|---|---|---|---|---|---|---|---|---|
| BOX001 | 41.50 | **58.50** | 0.00 | 37.30 | **62.70** | 0.00 | 47.90 | 51.50 | 0.60 | 46.80 | **52.40** | 0.80 | 48.40 | 51.50 | 0.10 |
| BOX022 | **49.20** | 38.90 | 11.90 | 44.50 | 43.70 | 11.80 | **44.40** | 36.60 | 19.00 | 42.70 | 41.80 | 15.50 | **45.30** | 38.70 | 16.00 |
| BOX032 | **50.40** | 49.60 | 0.00 | 34.80 | **65.20** | 0.00 | 53.60 | 44.30 | 2.10 | 47.90 | 50.00 | 2.10 | 53.50 | 46.40 | 0.10 |
| BOX034 | 50.00 | 49.90 | 0.10 | 38.20 | **61.70** | 0.10 | **68.50** | 25.50 | 6.00 | **54.50** | 40.10 | 5.40 | **55.60** | 40.60 | 3.80 |
| BOX035 | **63.40** | 36.60 | 0.00 | 48.60 | 51.40 | 0.00 | **72.00** | 23.90 | 4.10 | **62.80** | 36.10 | 1.10 | **72.10** | 27.70 | 0.20 |
| BOX046 | 52.20 | 46.60 | 1.20 | 48.30 | 50.50 | 1.20 | **51.70** | 21.50 | 26.80 | **60.30** | 31.70 | 8.00 | **52.70** | 21.60 | 25.70 |
| BOX060 | **58.30** | 41.60 | 0.10 | 53.40 | 46.50 | 0.10 | **70.90** | 17.70 | 11.40 | **59.60** | 32.30 | 8.10 | **71.30** | 25.30 | 3.40 |
| BOX063 | **64.30** | 35.70 | 0.00 | 42.60 | **57.40** | 0.00 | **69.30** | 30.30 | 0.40 | **67.70** | 31.40 | 0.90 | **68.20** | 31.50 | 0.30 |
| BOX076 | 46.70 | 50.00 | 3.30 | 44.50 | 52.00 | 3.50 | **47.10** | 34.60 | 18.30 | **52.40** | 38.00 | 9.60 | **51.80** | 33.60 | 14.60 |
| BOX096 | 53.40 | 46.60 | 0.00 | 38.90 | **61.10** | 0.00 | **80.00** | 19.20 | 0.80 | **76.90** | 22.00 | 1.10 | **80.80** | 19.10 | 0.10 |
| BOX115 | 51.40 | 48.60 | 0.00 | 45.00 | **55.00** | 0.00 | 35.70 | 32.90 | 31.40 | 49.50 | 48.80 | 1.70 | 50.10 | 48.90 | 1.00 |
| BOX121 | **63.70** | 36.30 | 0.00 | 45.00 | 55.00 | 0.00 | **69.60** | 27.00 | 3.40 | **67.60** | 31.30 | 1.10 | **69.90** | 29.80 | 0.30 |
| BOX122 | **66.00** | 31.20 | 2.80 | 46.40 | 50.80 | 2.80 | **54.20** | 40.10 | 5.70 | **53.10** | 41.50 | 5.40 | **54.00** | 40.90 | 5.10 |
| BOX123 | 42.10 | **57.90** | 0.00 | 48.10 | **51.90** | 0.00 | 36.10 | **58.00** | 5.90 | 33.80 | **64.80** | 1.40 | 36.50 | **63.40** | 0.10 |
| BOX142 | 43.70 | 50.80 | 5.50 | 29.00 | **65.50** | 5.50 | **48.60** | 42.30 | 9.10 | **48.80** | 43.10 | 8.10 | **49.10** | 42.70 | 8.20 |
| BOX172 | 51.70 | 48.30 | 0.00 | 42.90 | 57.10 | 0.00 | **32.40** | 27.20 | 40.40 | **50.30** | 42.00 | 7.70 | **38.90** | 31.80 | 29.30 |
| BOX175 | 42.10 | **57.90** | 0.00 | 44.30 | **55.70** | 0.00 | 31.80 | **54.50** | 13.70 | 33.50 | **60.80** | 5.70 | 36.10 | **58.90** | 5.00 |
| BOX177 | 41.80 | **57.90** | 0.30 | 49.20 | 50.80 | 0.00 | 53.20 | 43.20 | 3.60 | 51.20 | 46.00 | 2.80 | 52.50 | 46.10 | 1.40 |
| BOX180 | **60.80** | 38.90 | 0.30 | **64.40** | 35.30 | 0.30 | **58.50** | 35.80 | 5.70 | **60.00** | 35.80 | 4.20 | **60.60** | 36.60 | 2.80 |
| BOX181 | **60.30** | 39.30 | 0.40 | **66.20** | 33.40 | 0.40 | **56.00** | 36.70 | 7.30 | **56.00** | 37.80 | 6.20 | **58.10** | 37.30 | 4.60 |
| BOX192 | 43.60 | **55.60** | 0.80 | 41.90 | **57.30** | 0.80 | 39.00 | 36.30 | 24.70 | 42.70 | **50.50** | 6.80 | 44.60 | 49.50 | 5.90 |
| BOX212 | 48.20 | 51.70 | 0.10 | 48.10 | 51.90 | 0.00 | 48.20 | 44.00 | 7.80 | 48.10 | 49.50 | 2.40 | 49.40 | 43.80 | 6.80 |
| BOX214 | **58.00** | 39.00 | 3.00 | 48.10 | 49.10 | 2.80 | **56.40** | 30.70 | 12.90 | **58.00** | 34.10 | 7.90 | **59.20** | 32.20 | 8.60 |
| BOX222 | **62.20** | 37.70 | 0.10 | 53.60 | 46.30 | 0.10 | **68.00** | 29.30 | 2.70 | **62.00** | 36.60 | 1.40 | **70.70** | 29.00 | 0.30 |
| BOX240 | 46.40 | **50.00** | 3.60 | 47.10 | **49.40** | 3.50 | 45.90 | 43.10 | 11.00 | 46.90 | 47.30 | 5.80 | 48.30 | 45.50 | 6.20 |
| BOX246 | 52.70 | 47.20 | 0.10 | 51.60 | 48.40 | 0.00 | 47.80 | 46.90 | 5.30 | 42.40 | **54.90** | 2.70 | 46.20 | **52.70** | 1.10 |
| BOX258 | 45.50 | **53.80** | 0.70 | 34.10 | **65.20** | 0.70 | 41.10 | **53.80** | 5.10 | 33.40 | **61.70** | 4.90 | 36.10 | **60.70** | 3.20 |
| BOX270 | **56.80** | 42.60 | 0.60 | **53.10** | 46.30 | 0.60 | **59.10** | 34.10 | 6.80 | 52.90 | 43.30 | 3.80 | **60.20** | 37.90 | 1.90 |
| BOX284 | **63.80** | 36.10 | 0.10 | **65.80** | 34.20 | 0.00 | 43.40 | **52.70** | 3.90 | 46.30 | **52.10** | 1.60 | 46.50 | 52.20 | 1.30 |
| Average | **54.01** | 44.76 | 1.24 | 45.68 | **53.58** | 0.74 | **55.20** | 34.83 | 9.97 | **53.13** | 42.32 | 4.55 | **55.88** | 37.71 | 6.41 |

#### 6.7 BALiBASE RV912

Supplementary Table 41: Ankh-score vs ProtT5-score on BALiBASE RV912 dataset.

| Domain | $d_{cc}$ | | | $d_{d}$ | | | $d_{pos}$ | | | $d_{seq}$ | | | $d_{ssp}$ | | |
|---|---|---|---|---|---|---|---|---|---|---|---|---|---|---|---|
| Average distances | | | | | | | | | | | | | | | |
| | Ankh | ProtT5 | P-value | Ankh | ProtT5 | P-value | Ankh | ProtT5 | P-value | Ankh | ProtT5 | P-value | Ankh | ProtT5 | P-value |
| BOX011 | **0.0089** | 0.0109 | 7.4E-44 | 0.0070 | **0.0059** | 3.1E-139 | **0.2674** | 0.2899 | 6.2E-102 | **0.2340** | 0.2797 | 2.4E-99 | **0.4009** | 0.4594 | 7.8E-107 |
| BOX032 | **0.0032** | 0.0034 | 1.6E-12 | 0.0028 | **0.0027** | 1.4E-15 | **0.2445** | 0.2474 | 5.0E-12 | **0.2219** | 0.2246 | 9.3E-13 | **0.3748** | 0.3788 | 3.0E-21 |
| BOX045 | 0.0350 | 0.0332 | 8.2E-02 | 0.0254 | 0.0201 | 1.3E-02 | 0.4550 | 0.4505 | 2.2E-01 | 0.3448 | 0.3382 | 7.3E-01 | 0.5913 | 0.5849 | 9.8E-01 |
| BOX047 | 0.0142 | **0.0136** | 7.6E-11 | 0.0045 | **0.0039** | 1.3E-47 | 0.3206 | **0.3110** | 5.5E-04 | 0.2843 | **0.2721** | 1.5E-08 | 0.4555 | **0.4386** | 2.0E-11 |
| BOX049 | 0.0161 | **0.0142** | 5.5E-14 | 0.0078 | **0.0062** | 1.0E-22 | 0.2428 | **0.2253** | 7.2E-13 | 0.2287 | **0.2061** | 2.3E-16 | 0.3722 | **0.3452** | 1.4E-14 |
| BOX050 | 0.0150 | **0.0149** | 9.8E-04 | 0.0066 | **0.0041** | 1.0E-07 | 0.3417 | 0.3405 | 2.4E-01 | 0.3087 | 0.3000 | 3.2E-01 | 0.4963 | 0.4890 | 8.9E-01 |
| BOX054 | **0.0118** | 0.0141 | 1.3E-83 | **0.0041** | 0.0045 | 8.4E-94 | **0.3544** | 0.4071 | 8.1E-107 | **0.3006** | 0.3326 | 8.6E-98 | **0.5050** | 0.5597 | 2.5E-92 |
| BOX060 | **0.0097** | 0.0104 | 2.7E-04 | 0.0065 | **0.0061** | 9.7E-31 | 0.2025 | 0.2045 | 8.9E-01 | 0.1575 | 0.1565 | 7.2E-02 | 0.3086 | 0.3083 | 1.2E-01 |
| BOX075 | **0.0089** | 0.0093 | 1.8E-10 | **0.0018** | 0.0020 | 7.1E-48 | **0.2307** | 0.2481 | 3.7E-74 | **0.1833** | 0.1971 | 1.4E-89 | **0.3541** | 0.3768 | 4.5E-81 |
| BOX076 | **0.0091** | 0.0095 | 3.3E-37 | **0.0022** | 0.0023 | 3.1E-06 | **0.1807** | 0.1967 | 1.7E-123 | **0.1570** | 0.1712 | 1.7E-110 | **0.2965** | 0.3156 | 5.2E-119 |
| BOX096 | **0.0074** | 0.0076 | 4.7E-36 | **0.0007** | 0.0008 | 1.7E-03 | 0.1818 | 0.1848 | 6.0E-01 | **0.1533** | 0.1582 | 5.2E-46 | **0.2898** | 0.3130 | 8.0E-57 |
| BOX121 | 0.0093 | **0.0087** | 1.4E-03 | 0.0070 | **0.0066** | 1.7E-19 | 0.3123 | **0.2782** | 1.2E-05 | 0.2441 | 0.2388 | 3.4E-01 | 0.4694 | 0.4501 | 4.2E-01 |
| BOX122 | **0.0077** | 0.0085 | 7.4E-81 | **0.0010** | 0.0011 | 2.2E-44 | **0.1363** | 0.1587 | 6.6E-133 | **0.1303** | 0.1526 | 9.6E-132 | **0.2350** | 0.2672 | 2.4E-130 |
| BOX142 | **0.0063** | 0.0066 | 1.7E-28 | 0.0018 | **0.0017** | 8.2E-10 | 0.2768 | **0.2663** | 5.5E-22 | 0.2644 | **0.2514** | 5.0E-35 | 0.4196 | **0.4055** | 1.6E-30 |
| BOX149 | **0.0075** | 0.0076 | 3.5E-08 | 0.0021 | **0.0019** | 7.7E-12 | **0.2236** | 0.2335 | 4.8E-21 | **0.1896** | 0.1982 | 6.4E-17 | **0.3490** | 0.3620 | 1.1E-19 |
| BOX154 | 0.0193 | 0.0202 | 1.9E-02 | **0.0198** | 0.0202 | 2.1E-17 | 0.5219 | 0.5258 | 8.5E-01 | 0.4543 | 0.4537 | 9.5E-02 | **0.6634** | 0.6675 | 9.2E-03 |
| BOX175 | 0.0339 | 0.0332 | 7.4E-02 | 0.0116 | **0.0112** | 3.1E-22 | **0.5176** | 0.5254 | 3.4E-21 | **0.4948** | 0.4996 | 1.3E-09 | 0.6855 | **0.6839** | 1.2E-14 |
| BOX177 | **0.0101** | 0.0103 | 3.1E-03 | **0.0057** | 0.0064 | 3.1E-25 | **0.2933** | 0.2967 | 9.9E-08 | **0.2845** | 0.2872 | 1.0E-05 | **0.4310** | 0.4353 | 2.4E-07 |
| BOX187 | **0.0282** | 0.0293 | 1.2E-13 | **0.0061** | 0.0067 | 6.0E-17 | **0.3221** | 0.3282 | 6.9E-11 | **0.3090** | 0.3143 | 5.0E-12 | **0.4828** | 0.4886 | 3.2E-13 |
| BOX192 | **0.0493** | 0.0500 | 5.1E-03 | **0.0166** | 0.0166 | 7.3E-03 | 0.6134 | 0.6069 | 4.0E-01 | 0.5646 | 0.5476 | 2.2E-01 | 0.7553 | 0.7468 | 7.8E-02 |
| BOX202 | 0.0609 | **0.0578** | 3.0E-06 | 0.0259 | 0.0240 | 3.3E-02 | **0.4389** | 0.4539 | 1.3E-14 | **0.3975** | 0.4122 | 2.2E-10 | **0.5823** | 0.6027 | 1.3E-19 |
| BOX214 | **0.0069** | 0.0072 | 5.5E-06 | **0.0022** | 0.0023 | 1.4E-13 | 0.2039 | 0.2105 | 2.5E-02 | 0.1918 | 0.1978 | 5.2E-01 | 0.3119 | 0.3219 | 8.3E-02 |
| BOX240 | 0.0269 | 0.0283 | 3.1E-02 | 0.0072 | 0.0074 | 2.9E-02 | 0.4028 | **0.3976** | 2.4E-25 | 0.3819 | **0.3768** | 3.9E-18 | 0.5630 | **0.5598** | 4.8E-09 |
| BOX246 | 0.0223 | 0.0227 | 3.0E-02 | 0.0066 | **0.0057** | 4.9E-03 | **0.4035** | 0.4095 | 7.4E-33 | 0.3822 | **0.3821** | 4.8E-04 | **0.5591** | 0.5640 | 9.7E-19 |
| BOX258 | 0.0161 | 0.0147 | 1.8E-01 | 0.0145 | **0.0118** | 2.6E-04 | **0.3550** | 0.3595 | 8.4E-04 | 0.3420 | 0.3338 | 7.0E-01 | 0.5470 | 0.5470 | 8.1E-02 |
| BOX259 | **0.0050** | 0.0061 | 6.2E-57 | **0.0008** | 0.0014 | 2.4E-65 | **0.1041** | 0.1294 | 1.1E-75 | **0.0934** | 0.1184 | 3.3E-82 | **0.1868** | 0.2205 | 3.2E-79 |
| BOX281 | **0.0183** | 0.0217 | 6.3E-57 | 0.0071 | **0.0060** | 7.5E-04 | **0.3541** | 0.3971 | 2.2E-107 | **0.3250** | 0.3725 | 1.9E-98 | **0.5137** | 0.5588 | 2.6E-104 |
| BOX290 | **0.0246** | 0.0255 | 2.0E-09 | **0.0033** | 0.0035 | 2.8E-10 | **0.2475** | 0.2497 | 6.4E-22 | **0.2127** | 0.2268 | 5.0E-28 | 0.3844 | **0.3823** | 6.8E-21 |
| Domains won | **17** | 5 | | 12 | **13** | | **16** | 5 | | **15** | 5 | | **15** | 6 | |
| Tests won | | | | | | | | | | | | | | | |
| | Ankh | ProtT5 | Equal | Ankh | ProtT5 | Equal | Ankh | ProtT5 | Equal | Ankh | ProtT5 | Equal | Ankh | ProtT5 | Equal |
| BOX011 | **697** | 303 | 0 | 128 | **872** | 0 | **833** | 133 | 34 | **831** | 164 | 5 | **841** | 155 | 4 |
| BOX032 | **589** | 411 | 0 | 378 | **622** | 0 | **558** | 415 | 27 | **590** | 383 | 27 | **616** | 370 | 14 |
| BOX045 | 537 | 447 | 16 | 571 | 412 | 17 | 449 | 466 | 85 | 470 | 477 | 53 | 477 | 477 | 46 |
| BOX047 | 426 | **574** | 0 | 238 | **762** | 0 | 490 | **494** | 16 | 390 | **596** | 14 | 423 | **575** | 2 |
| BOX049 | 338 | **543** | 119 | 319 | **557** | 124 | 324 | **495** | 181 | 304 | **519** | 177 | 315 | **510** | 175 |
| BOX050 | 476 | **520** | 4 | 440 | **556** | 4 | 520 | 426 | 54 | 505 | 455 | 40 | 527 | 448 | 25 |
| BOX054 | **801** | 199 | 0 | **799** | 201 | 0 | **800** | 197 | 3 | **766** | 227 | 7 | **740** | 256 | 4 |
| BOX060 | 434 | **536** | 30 | 253 | **715** | 32 | 431 | 486 | 83 | 392 | 514 | 94 | 391 | 527 | 82 |
| BOX075 | **646** | 354 | 0 | **725** | 275 | 0 | **776** | 194 | 30 | **802** | 177 | 21 | **790** | 206 | 4 |
| BOX076 | **732** | 268 | 0 | **592** | 408 | 0 | **859** | 127 | 14 | **819** | 156 | 25 | **844** | 145 | 11 |
| BOX096 | **732** | 265 | 3 | **508** | 492 | 0 | 524 | 455 | 21 | **634** | 324 | 42 | **673** | 310 | 17 |
| BOX121 | 482 | **516** | 2 | 404 | **596** | 0 | **568** | 356 | 76 | 526 | 428 | 46 | 541 | 427 | 32 |
| BOX122 | **821** | 179 | 0 | **733** | 267 | 0 | **885** | 102 | 13 | **893** | 96 | 11 | **886** | 109 | 5 |
| BOX142 | 316 | **681** | 3 | 403 | **597** | 0 | 378 | **605** | 17 | 327 | **639** | 34 | 355 | **627** | 18 |
| BOX149 | **630** | 369 | 1 | **705** | 295 | 0 | **627** | 364 | 9 | **596** | 383 | 21 | **612** | 378 | 10 |
| BOX154 | 484 | 516 | 0 | 353 | **647** | 0 | 421 | 567 | 12 | 480 | 519 | 1 | 487 | **513** | 0 |
| BOX175 | 542 | 457 | 1 | 392 | **608** | 0 | **645** | 326 | 29 | **622** | 366 | 12 | **643** | 355 | 2 |
| BOX177 | 489 | **511** | 0 | **592** | 408 | 0 | **559** | 428 | 13 | **536** | 449 | 15 | **547** | 451 | 2 |
| BOX187 | **590** | 403 | 7 | **610** | 384 | 6 | **565** | 330 | 105 | **595** | 362 | 43 | **601** | 363 | 36 |
| BOX192 | **549** | 442 | 9 | **556** | 435 | 9 | 526 | 389 | 85 | 495 | 469 | 36 | 556 | 420 | 24 |
| BOX202 | **553** | 445 | 2 | 494 | 504 | 2 | **510** | 338 | 152 | **537** | 363 | 100 | **571** | 341 | 88 |
| BOX214 | **558** | 424 | 18 | **605** | 377 | 18 | 349 | 419 | 232 | 439 | 474 | 87 | 455 | 486 | 59 |
| BOX240 | 453 | 547 | 0 | 505 | 495 | 0 | 291 | **697** | 12 | 369 | **614** | 17 | 378 | **616** | 6 |
| BOX246 | 531 | 469 | 0 | 464 | **536** | 0 | **641** | 307 | 52 | **568** | 410 | 22 | **627** | 371 | 2 |
| BOX258 | 546 | 454 | 0 | 461 | **539** | 0 | **545** | 428 | 27 | 491 | 487 | 22 | 531 | 460 | 9 |
| BOX259 | **704** | 280 | 16 | **684** | 265 | 51 | **735** | 220 | 45 | **740** | 217 | 43 | **737** | 227 | 36 |
| BOX281 | **753** | 247 | 0 | 488 | **512** | 0 | **843** | 138 | 19 | **828** | 156 | 16 | **844** | 147 | 9 |
| BOX290 | **547** | 453 | 0 | **616** | 384 | 0 | **628** | 353 | 19 | **639** | 349 | 12 | **628** | 370 | 2 |
| Total | **12863** | 8923 | 214 | **12446** | 12310 | 244 | **13060** | 7047 | 893 | **12386** | 6950 | 664 | **13158** | 7395 | 447 |
| Tests won (%) | | | | | | | | | | | | | | | |
| BOX011 | **69.70** | 30.30 | 0.00 | 12.80 | **87.20** | 0.00 | **83.30** | 13.30 | 3.40 | **83.10** | 16.40 | 0.50 | **84.10** | 15.50 | 0.40 |
| BOX032 | **58.90** | 41.10 | 0.00 | 37.80 | **62.20** | 0.00 | **55.80** | 41.50 | 2.70 | **59.00** | 38.30 | 2.70 | **61.60** | 37.00 | 1.40 |
| BOX045 | 53.70 | 44.70 | 1.60 | 57.10 | 41.20 | 1.70 | 44.90 | 46.60 | 8.50 | 47.00 | 47.70 | 5.30 | 47.70 | 47.70 | 4.60 |
| BOX047 | 42.60 | **57.40** | 0.00 | 23.80 | **76.20** | 0.00 | 49.00 | **49.40** | 1.60 | 39.00 | **59.60** | 1.40 | 42.30 | **57.50** | 0.20 |
| BOX049 | 33.80 | **54.30** | 11.90 | 31.90 | **55.70** | 12.40 | 32.40 | **49.50** | 18.10 | 30.40 | **51.90** | 17.70 | 31.50 | **51.00** | 17.50 |
| BOX050 | 47.60 | **52.00** | 0.40 | 44.00 | **55.60** | 0.40 | 52.00 | 42.60 | 5.40 | 50.50 | 45.50 | 4.00 | 52.70 | 44.80 | 2.50 |
| BOX054 | **80.10** | 19.90 | 0.00 | **79.90** | 20.10 | 0.00 | **80.00** | 19.70 | 0.30 | **76.60** | 22.70 | 0.70 | **74.00** | 25.60 | 0.40 |
| BOX060 | 43.40 | **53.60** | 3.00 | 25.30 | **71.50** | 3.20 | 43.10 | 48.60 | 8.30 | 39.20 | 51.40 | 9.40 | 39.10 | 52.70 | 8.20 |
| BOX075 | **64.60** | 35.40 | 0.00 | **72.50** | 27.50 | 0.00 | **77.60** | 19.40 | 3.00 | **80.20** | 17.70 | 2.10 | **79.00** | 20.60 | 0.40 |
| BOX076 | **73.20** | 26.80 | 0.00 | **59.20** | 40.80 | 0.00 | **85.90** | 12.70 | 1.40 | **81.90** | 15.60 | 2.50 | **84.40** | 14.50 | 1.10 |
| BOX096 | **73.20** | 26.50 | 0.30 | **50.80** | 49.20 | 0.00 | 52.40 | 45.50 | 2.10 | **63.40** | 32.40 | 4.20 | **67.30** | 31.00 | 1.70 |
| BOX121 | 48.20 | **51.60** | 0.20 | 40.40 | **59.60** | 0.00 | **56.80** | 35.60 | 7.60 | 52.60 | 42.80 | 4.60 | 54.10 | 42.70 | 3.20 |
| BOX122 | **82.10** | 17.90 | 0.00 | **73.30** | 26.70 | 0.00 | **88.50** | 10.20 | 1.30 | **89.30** | 9.60 | 1.10 | **88.60** | 10.90 | 0.50 |
| BOX142 | 31.60 | **68.10** | 0.30 | 40.30 | **59.70** | 0.00 | 37.80 | **60.50** | 1.70 | 32.70 | **63.90** | 3.40 | 35.50 | **62.70** | 1.80 |
| BOX149 | **63.00** | 36.90 | 0.10 | **70.50** | 29.50 | 0.00 | **62.70** | 36.40 | 0.90 | **59.60** | 38.30 | 2.10 | **61.20** | 37.80 | 1.00 |
| BOX154 | 48.40 | 51.60 | 0.00 | 35.30 | **64.70** | 0.00 | 42.10 | 56.70 | 1.20 | 48.00 | 51.90 | 0.10 | 48.70 | **51.30** | 0.00 |
| BOX175 | 54.20 | 45.70 | 0.10 | 39.20 | **60.80** | 0.00 | **64.50** | 32.60 | 2.90 | **62.20** | 36.60 | 1.20 | **64.30** | 35.50 | 0.20 |
| BOX177 | 48.90 | **51.10** | 0.00 | **59.20** | 40.80 | 0.00 | **55.90** | 42.80 | 1.30 | **53.60** | 44.90 | 1.50 | **54.70** | 45.10 | 0.20 |
| BOX187 | **59.00** | 40.30 | 0.70 | **61.00** | 38.40 | 0.60 | **56.50** | 33.00 | 10.50 | **59.50** | 36.20 | 4.30 | **60.10** | 36.30 | 3.60 |
| BOX192 | **54.90** | 44.20 | 0.90 | **55.60** | 43.50 | 0.90 | 52.60 | 38.90 | 8.50 | 49.50 | 46.90 | 3.60 | 55.60 | 42.00 | 2.40 |
| BOX202 | **55.30** | 44.50 | 0.20 | 49.40 | 50.40 | 0.20 | **51.00** | 33.80 | 15.20 | **53.70** | 36.30 | 10.00 | **57.10** | 34.10 | 8.80 |
| BOX214 | **55.80** | 42.40 | 1.80 | **60.50** | 37.70 | 1.80 | 34.90 | 41.90 | 23.20 | 43.90 | 47.40 | 8.70 | 45.50 | 48.60 | 5.90 |
| BOX240 | 45.30 | 54.70 | 0.00 | 50.50 | 49.50 | 0.00 | 29.10 | **69.70** | 1.20 | 36.90 | **61.40** | 1.70 | 37.80 | **61.60** | 0.60 |
| BOX246 | 53.10 | 46.90 | 0.00 | 46.40 | **53.60** | 0.00 | **64.10** | 30.70 | 5.20 | **56.80** | 41.00 | 2.20 | **62.70** | 37.10 | 0.20 |
| BOX258 | 54.60 | 45.40 | 0.00 | 46.10 | **53.90** | 0.00 | **54.50** | 42.80 | 2.70 | 49.10 | 48.70 | 2.20 | 53.10 | 46.00 | 0.90 |
| BOX259 | **70.40** | 28.00 | 1.60 | **68.40** | 26.50 | 5.10 | **73.50** | 22.00 | 4.50 | **74.00** | 21.70 | 4.30 | **73.70** | 22.70 | 3.60 |
| BOX281 | **75.30** | 24.70 | 0.00 | 48.80 | **51.20** | 0.00 | **84.30** | 13.80 | 1.90 | **82.80** | 15.60 | 1.60 | **84.40** | 14.70 | 0.90 |
| BOX290 | **54.70** | 45.30 | 0.00 | **61.60** | 38.40 | 0.00 | **62.80** | 35.30 | 1.90 | **63.90** | 34.90 | 1.20 | **62.80** | 37.00 | 0.20 |
| Average | **58.47** | 40.56 | 0.97 | **49.78** | 49.24 | 0.98 | **62.19** | 33.56 | 4.25 | **61.93** | 34.75 | 3.32 | **62.66** | 35.21 | 2.13 |

#### 6.8 BALiBASE RV913

Supplementary Table 42: Ankh-score vs ProtT5-score on BALiBASE RV913 dataset.

| Domain | $d_{cc}$ | | | $d_d$ | | | $d_{pos}$ | | | $d_{seq}$ | | | $d_{ssp}$ | | |
|---|---|---|---|---|---|---|---|---|---|---|---|---|---|---|---|
| Average distances | | | | | | | | | | | | | | | |
| | Ankh | ProtT5 | P-value | Ankh | ProtT5 | P-value | Ankh | ProtT5 | P-value | Ankh | ProtT5 | P-value | Ankh | ProtT5 | P-value |
| BOX012 | **0.0076** | 0.0078 | 1.3E-15 | **0.0020** | 0.0020 | 1.4E-08 | 0.2134 | **0.2050** | 1.7E-24 | 0.2193 | **0.1946** | 1.6E-21 | **0.3723** | 0.3802 | 1.4E-19 |
| BOX017 | 0.0154 | 0.0164 | 9.2E-01 | **0.0032** | 0.0039 | 8.7E-06 | **0.2872** | 0.2888 | 3.6E-03 | 0.2677 | 0.2657 | 7.5E-01 | 0.4249 | 0.4237 | 9.0E-02 |
| BOX032 | 0.0012 | **0.0010** | 2.9E-38 | 0.0016 | **0.0015** | 6.6E-156 | 0.1005 | **0.0989** | 1.4E-07 | 0.0944 | **0.0927** | 1.6E-09 | 0.1769 | **0.1744** | 1.1E-06 |
| BOX034 | **0.0015** | 0.0018 | 1.3E-74 | **0.0005** | 0.0009 | 2.0E-79 | **0.0938** | 0.0989 | 3.4E-32 | **0.0793** | 0.0857 | 1.4E-45 | **0.1630** | 0.1711 | 2.2E-35 |
| BOX036 | 0.0024 | **0.0023** | 3.9E-07 | 0.0021 | 0.0015 | 7.7E-01 | **0.1005** | 0.1011 | 9.6E-06 | 0.0927 | **0.0919** | 4.2E-04 | 0.1733 | **0.1726** | 3.6E-04 |
| BOX043 | **0.0007** | 0.0007 | 6.4E-14 | 0.0002 | 0.0003 | 1.4E-02 | **0.0583** | 0.0610 | 3.2E-04 | **0.0558** | 0.0578 | 5.7E-06 | **0.1091** | 0.1136 | 1.1E-07 |
| BOX049 | **0.0061** | 0.0066 | 7.2E-05 | **0.0027** | 0.0029 | 1.1E-03 | 0.1115 | 0.1147 | 2.1E-01 | 0.1082 | 0.1129 | 1.3E-01 | 0.1877 | 0.1944 | 8.7E-02 |
| BOX063 | 0.0049 | **0.0047** | 6.5E-05 | **0.0011** | 0.0012 | 5.0E-17 | **0.1259** | 0.1301 | 8.7E-24 | **0.1169** | 0.1217 | 1.0E-28 | **0.2174** | 0.2248 | 2.3E-29 |
| BOX075 | **0.0011** | 0.0013 | 4.8E-28 | **0.0001** | 0.0002 | 4.2E-09 | **0.0391** | 0.0487 | 1.6E-57 | **0.0377** | 0.0435 | 7.9E-63 | **0.0749** | 0.0923 | 4.4E-62 |
| BOX076 | **0.0068** | 0.0068 | 4.4E-14 | 0.0003 | 0.0003 | 8.5E-01 | **0.0786** | 0.0814 | 1.4E-14 | **0.0764** | 0.0787 | 3.0E-12 | **0.1440** | 0.1480 | 1.2E-13 |
| BOX079 | **0.0026** | 0.0027 | 2.9E-19 | **0.0013** | 0.0013 | 2.2E-11 | **0.1089** | 0.1157 | 4.7E-49 | **0.0897** | 0.0951 | 2.8E-50 | **0.1822** | 0.1900 | 1.7E-49 |
| BOX082 | **0.0008** | 0.0009 | 9.8E-16 | 0.0004 | **0.0003** | 2.0E-08 | **0.0568** | 0.0572 | 5.2E-11 | **0.0478** | 0.0482 | 4.9E-10 | 0.1052 | **0.1049** | 1.5E-11 |
| BOX121 | **0.0020** | 0.0025 | 8.8E-46 | **0.0009** | 0.0009 | 5.5E-06 | **0.1399** | 0.1493 | 1.5E-45 | **0.1383** | 0.1479 | 1.9E-44 | **0.2405** | 0.2541 | 3.5E-44 |
| BOX122 | **0.0038** | 0.0039 | 2.6E-17 | **0.0001** | 0.0001 | 7.7E-16 | **0.0453** | 0.0525 | 1.5E-37 | **0.0447** | 0.0520 | 1.6E-38 | **0.0860** | 0.0985 | 5.8E-38 |
| BOX126 | 0.0045 | **0.0044** | 5.9E-12 | **0.0004** | 0.0004 | 9.5E-12 | **0.0680** | 0.0766 | 7.1E-52 | **0.0645** | 0.0738 | 1.7E-58 | **0.1257** | 0.1420 | 2.6E-59 |
| BOX132 | 0.0222 | 0.0216 | 9.3E-02 | 0.0017 | **0.0016** | 1.3E-08 | 0.1504 | 0.1518 | 5.5E-02 | 0.1476 | 0.1482 | 2.1E-01 | 0.2530 | 0.2551 | 6.8E-02 |
| BOX133 | 0.0106 | **0.0105** | 3.3E-06 | 0.0009 | 0.0009 | 2.0E-02 | 0.1393 | 0.1374 | 2.4E-01 | 0.1329 | 0.1308 | 1.8E-01 | 0.2383 | 0.2357 | 2.2E-01 |
| BOX142 | 0.0016 | **0.0015** | 1.6E-03 | **0.0002** | 0.0003 | 7.7E-17 | 0.0631 | **0.0577** | 6.0E-43 | 0.0533 | **0.0477** | 8.8E-38 | 0.1115 | **0.1016** | 2.0E-45 |
| BOX146 | **0.0077** | 0.0078 | 6.6E-22 | 0.0010 | **0.0009** | 2.6E-05 | **0.0792** | 0.0922 | 1.2E-78 | **0.0696** | 0.0838 | 2.0E-73 | **0.1393** | 0.1613 | 1.1E-75 |
| BOX158 | 0.0058 | 0.0057 | 8.3E-01 | 0.0005 | 0.0005 | 4.2E-02 | **0.0619** | 0.0632 | 1.0E-22 | **0.0550** | 0.0570 | 9.4E-22 | **0.1116** | 0.1148 | 4.9E-26 |
| BOX180 | **0.0025** | 0.0033 | 4.0E-115 | **0.0011** | 0.0016 | 4.3E-113 | **0.1131** | 0.1345 | 1.4E-106 | 0.1340 | **0.1278** | 8.1E-120 | **0.2400** | 0.2748 | 1.5E-115 |
| BOX183 | **0.0025** | 0.0033 | 6.8E-45 | **0.0013** | 0.0013 | 1.7E-39 | **0.0989** | 0.1081 | 5.4E-46 | **0.0798** | 0.0954 | 6.2E-46 | **0.1783** | 0.2008 | 3.4E-43 |
| BOX214 | 0.0000 | **0.0000** | 1.9E-15 | 0.0000 | **0.0000** | 1.2E-26 | 0.0031 | **0.0000** | 2.7E-47 | 0.0031 | **0.0000** | 2.7E-47 | 0.0061 | **0.0008** | 2.2E-45 |
| BOX222 | 0.0093 | 0.0091 | 2.1E-01 | 0.0022 | 0.0021 | 2.1E-01 | **0.1939** | 0.2042 | 3.8E-53 | **0.1672** | 0.1744 | 3.4E-33 | **0.3111** | 0.3200 | 5.2E-32 |
| BOX246 | 0.0295 | **0.0263** | 2.1E-16 | 0.0059 | **0.0038** | 9.8E-33 | 0.2239 | **0.2134** | 1.0E-16 | 0.2131 | **0.2040** | 3.0E-05 | 0.3696 | **0.3555** | 1.7E-06 |
| BOX258 | 0.0129 | **0.0126** | 4.3E-14 | **0.0015** | 0.0017 | 3.1E-45 | **0.1704** | 0.1908 | 1.4E-66 | **0.1446** | 0.1520 | 1.0E-31 | **0.2860** | 0.2948 | 1.1E-32 |
| BOX290 | 0.0039 | **0.0037** | 7.4E-12 | **0.0002** | 0.0002 | 6.4E-43 | 0.0641 | **0.0590** | 9.0E-56 | 0.0445 | **0.0375** | 2.0E-59 | 0.1025 | **0.0920** | 2.4E-56 |
| Domains won | **13** | 10 | | **15** | 6 | | **18** | 6 | | **15** | 8 | | **16** | 7 | |
| Tests won | | | | | | | | | | | | | | | |
| | Ankh | ProtT5 | Equal | Ankh | ProtT5 | Equal | Ankh | ProtT5 | Equal | Ankh | ProtT5 | Equal | Ankh | ProtT5 | Equal |
| BOX012 | **605** | 296 | 99 | **556** | 345 | 99 | **595** | 290 | 115 | **598** | 286 | 116 | **604** | 294 | 102 |
| BOX017 | 425 | 482 | 93 | 387 | **520** | 93 | **409** | 290 | 301 | 390 | 404 | 206 | 423 | 377 | 200 |
| BOX032 | 319 | **681** | 0 | 45 | **955** | 0 | 419 | **555** | 26 | 381 | **567** | 52 | 433 | **561** | 6 |
| BOX034 | **778** | 212 | 10 | **738** | 252 | 10 | **629** | 254 | 117 | **698** | 247 | 55 | **682** | 273 | 45 |
| BOX036 | **597** | 394 | 9 | 620 | 371 | 9 | **583** | 364 | 53 | **574** | 390 | 36 | **590** | 383 | 27 |
| BOX043 | **656** | 337 | 7 | 573 | 418 | 9 | **550** | 403 | 47 | **565** | 407 | 28 | **576** | 401 | 23 |
| BOX049 | **418** | 306 | 276 | **409** | 304 | 287 | 316 | 339 | 345 | 324 | 316 | 360 | 323 | 334 | 343 |
| BOX063 | **576** | 424 | 0 | **658** | 342 | 0 | **640** | 355 | 5 | **658** | 320 | 22 | **666** | 333 | 1 |
| BOX075 | **641** | 226 | 133 | **528** | 337 | 135 | **578** | 243 | 179 | **634** | 193 | 173 | **633** | 194 | 173 |
| BOX076 | **637** | 339 | 24 | 525 | 452 | 23 | **591** | 311 | 98 | **588** | 334 | 78 | **601** | 330 | 69 |
| BOX079 | **610** | 349 | 41 | **634** | 326 | 40 | **617** | 265 | 118 | **623** | 255 | 122 | **615** | 267 | 118 |
| BOX082 | **616** | 355 | 29 | **556** | 414 | 30 | **548** | 356 | 96 | **585** | 364 | 51 | **596** | 356 | 48 |
| BOX121 | **681** | 280 | 39 | **581** | 381 | 38 | **678** | 221 | 101 | **682** | 230 | 88 | **685** | 243 | 72 |
| BOX122 | **509** | 325 | 166 | **527** | 297 | 176 | **545** | 219 | 236 | **554** | 231 | 215 | **552** | 235 | 213 |
| BOX126 | **622** | 351 | 27 | **601** | 371 | 28 | **690** | 191 | 119 | **697** | 213 | 90 | **697** | 217 | 86 |
| BOX132 | 358 | 387 | 255 | 311 | **434** | 255 | 321 | 227 | 452 | 330 | 267 | 403 | 337 | 262 | 401 |
| BOX133 | 444 | **550** | 6 | 488 | 507 | 5 | 509 | 466 | 25 | 494 | 477 | 29 | 509 | 472 | 19 |
| BOX142 | 375 | **514** | 111 | **496** | 393 | 111 | 300 | **578** | 122 | 294 | **552** | 154 | 285 | **581** | 134 |
| BOX146 | **667** | 325 | 8 | **620** | 370 | 10 | **794** | 174 | 32 | **776** | 180 | 44 | **792** | 184 | 24 |
| BOX158 | 531 | 453 | 16 | 516 | 470 | 14 | **622** | 312 | 66 | **613** | 328 | 59 | **631** | 326 | 43 |
| BOX180 | **785** | 65 | 150 | **772** | 78 | 150 | **734** | 53 | 213 | **796** | 47 | 157 | **794** | 49 | 157 |
| BOX183 | **648** | 269 | 83 | **691** | 228 | 81 | **616** | 274 | 110 | **625** | 260 | 115 | **627** | 263 | 110 |
| BOX214 | 49 | **295** | 656 | 37 | **300** | 663 | 35 | **310** | 655 | 35 | **310** | 655 | 35 | **310** | 655 |
| BOX222 | 576 | 419 | 5 | 552 | 444 | 4 | **685** | 297 | 18 | **611** | 362 | 27 | **612** | 372 | 16 |
| BOX246 | 415 | **584** | 1 | 343 | **656** | 1 | 390 | **419** | 191 | **543** | 439 | 18 | **555** | 429 | 16 |
| BOX258 | **663** | 337 | 0 | **684** | 316 | 0 | **692** | 289 | 19 | **704** | 277 | 19 | **718** | 277 | 5 |
| BOX290 | 381 | **435** | 184 | 232 | **525** | 243 | 167 | **530** | 303 | 167 | **552** | 281 | 171 | **570** | 259 |
| Total | **12692** | 8249 | 2059 | **10406** | 8144 | 2450 | **13107** | 7553 | 3340 | **13001** | 7344 | 2655 | **13150** | 7448 | 2402 |
| Tests won (%) | | | | | | | | | | | | | | | |
| BOX012 | **60.50** | 29.60 | 9.90 | **55.60** | 34.50 | 9.90 | **59.50** | 29.00 | 11.50 | **59.80** | 28.60 | 11.60 | **60.40** | 29.40 | 10.20 |
| BOX017 | 42.50 | 48.20 | 9.30 | 38.70 | **52.00** | 9.30 | **40.90** | 29.00 | 30.10 | 39.00 | 40.40 | 20.60 | 42.30 | 37.70 | 20.00 |
| BOX032 | 31.90 | **68.10** | 0.00 | 4.50 | **95.50** | 0.00 | 41.90 | **55.50** | 2.60 | 38.10 | **56.70** | 5.20 | 43.30 | **56.10** | 0.60 |
| BOX034 | **77.80** | 21.20 | 1.00 | **73.80** | 25.20 | 1.00 | **62.90** | 25.40 | 11.70 | **69.80** | 24.70 | 5.50 | **68.20** | 27.30 | 4.50 |
| BOX036 | **59.70** | 39.40 | 0.90 | 62.00 | 37.10 | 0.90 | **58.30** | 36.40 | 5.30 | **57.40** | 39.00 | 3.60 | **59.00** | 38.30 | 2.70 |
| BOX043 | **65.60** | 33.70 | 0.70 | 57.30 | 41.80 | 0.90 | **55.00** | 40.30 | 4.70 | **56.50** | 40.70 | 2.80 | **57.60** | 40.10 | 2.30 |
| BOX049 | **41.80** | 30.60 | 27.60 | **40.90** | 30.40 | 28.70 | 31.60 | 33.90 | 34.50 | 32.40 | 31.60 | 36.00 | 32.30 | 33.40 | 34.30 |
| BOX063 | **57.60** | 42.40 | 0.00 | **65.80** | 34.20 | 0.00 | **64.00** | 35.50 | 0.50 | **65.80** | 32.00 | 2.20 | **66.60** | 33.30 | 0.10 |
| BOX075 | **64.10** | 22.60 | 13.30 | **52.80** | 33.70 | 13.50 | **57.80** | 24.30 | 17.90 | **63.40** | 19.30 | 17.30 | **63.30** | 19.40 | 17.30 |
| BOX076 | **63.70** | 33.90 | 2.40 | 52.50 | 45.20 | 2.30 | **59.10** | 31.10 | 9.80 | **58.80** | 33.40 | 7.80 | **60.10** | 33.00 | 6.90 |
| BOX079 | **61.00** | 34.90 | 4.10 | **63.40** | 32.60 | 4.00 | **61.70** | 26.50 | 11.80 | **62.30** | 25.50 | 12.20 | **61.50** | 26.70 | 11.80 |
| BOX082 | **61.60** | 35.50 | 2.90 | **55.60** | 41.40 | 3.00 | **54.80** | 35.60 | 9.60 | **58.50** | 36.40 | 5.10 | **59.60** | 35.60 | 4.80 |
| BOX121 | **68.10** | 28.00 | 3.90 | **58.10** | 38.10 | 3.80 | **67.80** | 22.10 | 10.10 | **68.20** | 23.00 | 8.80 | **68.50** | 24.30 | 7.20 |
| BOX122 | **50.90** | 32.50 | 16.60 | **52.70** | 29.70 | 17.60 | **54.50** | 21.90 | 23.60 | **55.40** | 23.10 | 21.50 | **55.20** | 23.50 | 21.30 |
| BOX126 | **62.20** | 35.10 | 2.70 | **60.10** | 37.10 | 2.80 | **69.00** | 19.10 | 11.90 | **69.70** | 21.30 | 9.00 | **69.70** | 21.70 | 8.60 |
| BOX132 | 35.80 | 38.70 | 25.50 | 31.10 | **43.40** | 25.50 | 32.10 | 22.70 | 45.20 | 33.00 | 26.70 | 40.30 | 33.70 | 26.20 | 40.10 |
| BOX133 | 44.40 | **55.00** | 0.60 | 48.80 | 50.70 | 0.50 | 50.90 | 46.60 | 2.50 | 49.40 | 47.70 | 2.90 | 50.90 | 47.20 | 1.90 |
| BOX142 | 37.50 | **51.40** | 11.10 | **49.60** | 39.30 | 11.10 | 30.00 | **57.80** | 12.20 | 29.40 | **55.20** | 15.40 | 28.50 | **58.10** | 13.40 |
| BOX146 | **66.70** | 32.50 | 0.80 | **62.00** | 37.00 | 1.00 | **79.40** | 17.40 | 3.20 | **77.60** | 18.00 | 4.40 | **79.20** | 18.40 | 2.40 |
| BOX158 | 53.10 | 45.30 | 1.60 | 51.60 | 47.00 | 1.40 | **62.20** | 31.20 | 6.60 | **61.30** | 32.80 | 5.90 | **63.10** | 32.60 | 4.30 |
| BOX180 | **78.50** | 6.50 | 15.00 | **77.20** | 7.80 | 15.00 | **73.40** | 5.30 | 21.30 | **79.60** | 4.70 | 15.70 | **79.40** | 4.90 | 15.70 |
| BOX183 | **64.80** | 26.90 | 8.30 | **69.10** | 22.80 | 8.10 | **61.60** | 27.40 | 11.00 | **62.50** | 26.00 | 11.50 | **62.70** | 26.30 | 11.00 |
| BOX214 | 4.90 | **29.50** | 65.60 | 3.70 | **30.00** | 66.30 | 3.50 | **31.00** | 65.50 | 3.50 | **31.00** | 65.50 | 3.50 | **31.00** | 65.50 |
| BOX222 | 57.60 | 41.90 | 0.50 | 55.20 | 44.40 | 0.40 | **68.50** | 29.70 | 1.80 | **61.10** | 36.20 | 2.70 | **61.20** | 37.20 | 1.60 |
| BOX246 | 41.50 | **58.40** | 0.10 | 34.30 | **65.60** | 0.10 | 39.00 | **41.90** | 19.10 | **54.30** | 43.90 | 1.80 | **55.50** | 42.90 | 1.60 |
| BOX258 | **66.30** | 33.70 | 0.00 | **68.40** | 31.60 | 0.00 | **69.20** | 28.90 | 1.90 | **70.40** | 27.70 | 1.90 | **71.80** | 27.70 | 0.50 |
| BOX290 | 38.10 | **43.50** | 18.40 | 23.20 | **52.50** | 24.30 | 16.70 | **53.00** | 30.30 | 16.70 | **55.20** | 28.10 | 17.10 | **57.00** | 25.90 |
| Average | **55.18** | 35.87 | 8.95 | **49.55** | 38.78 | 11.67 | **54.61** | 31.47 | 13.92 | **56.53** | 31.93 | 11.54 | **57.17** | 32.38 | 10.44 |

### 7 Comparison against DEDAL

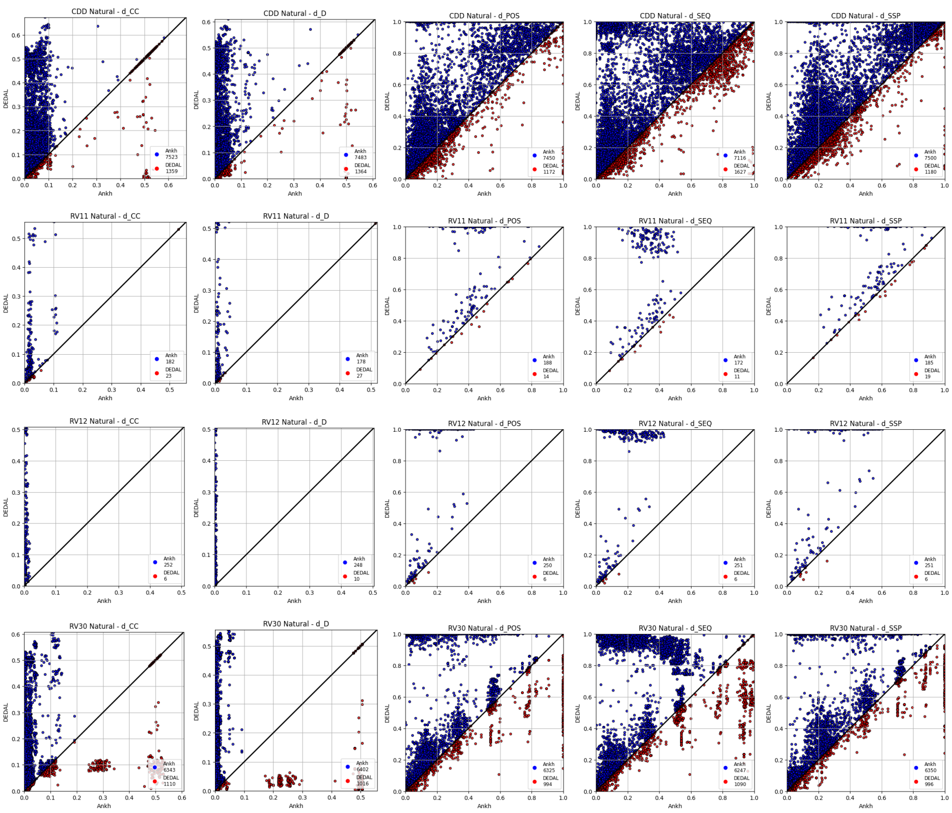


Supplementary Figure 2: Ankh-score vs DEDAL on all natural datasets.

#### 7.1 CDD natural

Supplementary Table 43: Ankh-score vs DEDAL on CDD natural dataset.

| Domain | $d_{cc}$ | | | $d_d$ | | | $d_{pos}$ | | | $d_{seq}$ | | | $d_{ssp}$ | | |
|---|---|---|---|---|---|---|---|---|---|---|---|---|---|---|---|
| Average distances | | | | | | | | | | | | | | | |
| | Ankh | DEDAL | P-value | Ankh | DEDAL | P-value | Ankh | DEDAL | P-value | Ankh | DEDAL | P-value | Ankh | DEDAL | P-value |
| cd00012 | **0.0446** | 0.1971 | 2.3E-264 | **0.0235** | 0.1782 | 6.9E-252 | **0.6738** | 0.9042 | 8.5E-268 | **0.5926** | 0.7265 | 4.3E-127 | **0.7964** | 0.9477 | 8.0E-264 |
| cd00083 | **0.0026** | 0.2618 | 8.3E-120 | **0.0006** | 0.2527 | 2.9E-116 | **0.0430** | 0.5823 | 8.4E-121 | **0.0333** | 0.5545 | 5.0E-121 | **0.0885** | 0.6096 | 1.5E-121 |
| cd00173 | **0.0335** | 0.4757 | 4.1E-41 | **0.0081** | 0.4768 | 1.1E-40 | **0.4884** | 1.0000 | 2.2E-41 | **0.4576** | 0.9304 | 6.2E-42 | **0.5958** | 1.0000 | 3.2E-41 |
| cd00637 | **0.0044** | 0.0063 | 7.7E-229 | **0.0010** | 0.0016 | 2.1E-228 | **0.1315** | 0.1803 | 2.1E-255 | **0.1055** | 0.1554 | 2.0E-270 | **0.2211** | 0.2864 | 5.1E-268 |
| cd01040 | **0.0037** | 0.0149 | 2.5E-236 | **0.0007** | 0.0027 | 3.3E-234 | **0.1006** | 0.2332 | 2.2E-247 | **0.0795** | 0.2153 | 1.9E-249 | **0.1741** | 0.3576 | 3.7E-250 |
| cd01068 | **0.0027** | 0.0354 | 8.7E-23 | **0.0006** | 0.0069 | 3.4E-21 | **0.1046** | 0.2197 | 6.5E-25 | **0.1028** | 0.2170 | 8.6E-25 | **0.1720** | 0.4720 | 5.2E-25 |
| cd10140 | **0.0596** | 0.1274 | 1.6E-93 | **0.0120** | 0.1033 | 9.0E-98 | **0.7260** | 0.9017 | 5.5E-59 | **0.6739** | 0.8067 | 4.3E-25 | **0.8359** | 0.9481 | 2.2E-57 |
| cd21116 | **0.0084** | 0.0570 | 1.4E-48 | **0.0029** | 0.0311 | 7.4E-49 | **0.2944** | 0.5625 | 3.8E-48 | **0.2594** | 0.5186 | 3.5E-45 | **0.4129** | 0.6666 | 6.4E-48 |
| Domains won | **8** | 0 | | **8** | 0 | | **8** | 0 | | **8** | 0 | | **8** | 0 | |
| Tests won | | | | | | | | | | | | | | | |
| | Ankh | DEDAL | Equal | Ankh | DEDAL | Equal | Ankh | DEDAL | Equal | Ankh | DEDAL | Equal | Ankh | DEDAL | Equal |
| cd00012 | **1646** | 123 | 1 | **1593** | 176 | 1 | **1632** | 99 | 39 | **1320** | 442 | 8 | **1623** | 110 | 37 |
| cd00083 | **803** | 109 | 169 | **788** | 100 | 193 | **788** | 93 | 200 | **791** | 93 | 197 | **790** | 93 | 198 |
| cd00173 | **243** | 8 | 74 | **241** | 10 | 74 | **238** | 6 | 81 | **242** | 7 | 76 | **237** | 7 | 81 |
| cd00637 | **2215** | 787 | 1 | **2249** | 753 | 1 | **2279** | 670 | 54 | **2332** | 626 | 45 | **2331** | 652 | 20 |
| cd01040 | **1614** | 222 | 55 | **1603** | 223 | 65 | **1611** | 181 | 99 | **1620** | 177 | 94 | **1625** | 180 | 86 |
| cd01068 | **147** | 15 | 9 | **140** | 22 | 9 | **145** | 15 | 11 | **143** | 15 | 13 | **144** | 15 | 12 |
| cd10140 | **562** | 65 | 3 | **577** | 49 | 4 | **464** | 82 | 84 | **382** | 233 | 15 | **459** | 93 | 78 |
| cd21116 | **293** | 30 | 2 | **292** | 31 | 2 | **293** | 26 | 6 | **286** | 34 | 5 | **291** | 30 | 4 |
| Total | **7523** | 1359 | 314 | **7483** | 1364 | 349 | **7450** | 1172 | 574 | **7116** | 1627 | 453 | **7500** | 1180 | 516 |
| Tests won (%) | | | | | | | | | | | | | | | |
| cd00012 | **92.99** | 6.95 | 0.06 | **90.00** | 9.94 | 0.06 | **92.20** | 5.59 | 2.20 | **74.58** | 24.97 | 0.45 | **91.69** | 6.21 | 2.09 |
| cd00083 | **74.28** | 10.08 | 15.63 | **72.90** | 9.25 | 17.85 | **72.90** | 8.60 | 18.50 | **73.17** | 8.60 | 18.22 | **73.08** | 8.60 | 18.32 |
| cd00173 | **74.77** | 2.46 | 22.77 | **74.15** | 3.08 | 22.77 | **73.23** | 1.85 | 24.92 | **74.46** | 2.15 | 23.38 | **72.92** | 2.15 | 24.92 |
| cd00637 | **73.76** | 26.21 | 0.03 | **74.89** | 25.07 | 0.03 | **75.89** | 22.31 | 1.80 | **77.66** | 20.85 | 1.50 | **77.62** | 21.71 | 0.67 |
| cd01040 | **85.35** | 11.74 | 2.91 | **84.77** | 11.79 | 3.44 | **85.19** | 9.57 | 5.24 | **85.67** | 9.36 | 4.97 | **85.93** | 9.52 | 4.55 |
| cd01068 | **85.96** | 8.77 | 5.26 | **81.87** | 12.87 | 5.26 | **84.80** | 8.77 | 6.43 | **83.63** | 8.77 | 7.60 | **84.21** | 8.77 | 7.02 |
| cd10140 | **89.21** | 10.32 | 0.48 | **91.59** | 7.78 | 0.63 | **73.65** | 13.02 | 13.33 | **60.63** | 36.98 | 2.38 | **72.86** | 14.76 | 12.38 |
| cd21116 | **90.15** | 9.23 | 0.62 | **89.85** | 9.54 | 0.62 | **90.15** | 8.00 | 1.85 | **88.00** | 10.46 | 1.54 | **89.54** | 9.23 | 1.23 |
| Average | **83.31** | 10.72 | 5.97 | **82.50** | 11.17 | 6.33 | **81.00** | 9.71 | 9.28 | **77.22** | 15.27 | 7.51 | **80.98** | 10.12 | 8.90 |

#### 7.2 BALiBASE RV11

Supplementary Table 44: Ankh-score vs DEDAL on BALiBASE RV11 dataset.

| Domain | $d_{cc}$ | | | $d_d$ | | | $d_{pos}$ | | | $d_{seq}$ | | | $d_{ssp}$ | | |
|---|---|---|---|---|---|---|---|---|---|---|---|---|---|---|---|
| Average distances | | | | | | | | | | | | | | | |
| | Ankh | DEDAL | P-value | Ankh | DEDAL | P-value | Ankh | DEDAL | P-value | Ankh | DEDAL | P-value | Ankh | DEDAL | P-value |
| BBS11009 | 0.0207 | 0.0337 | 6.9E-01 | 0.0103 | 0.0127 | 5.6E-01 | 0.2317 | 0.3992 | 4.4E-01 | 0.2134 | 0.3692 | 4.4E-01 | 0.3654 | 0.5348 | 4.4E-01 |
| BBS11010 | 0.0232 | 0.0438 | 3.1E-02 | 0.0039 | 0.0287 | 3.1E-02 | 0.2844 | 0.5066 | 9.4E-02 | 0.2093 | 0.4269 | 6.3E-02 | 0.4258 | 0.6366 | 6.3E-02 |
| BBS11016 | **0.0111** | 0.0379 | 1.1E-04 | **0.0041** | 0.0065 | 7.2E-05 | **0.3578** | 0.6666 | 9.8E-06 | **0.3195** | 0.6094 | 2.5E-07 | **0.5197** | 0.7567 | 2.5E-07 |
| BBS11018 | **0.0190** | 0.0862 | 1.8E-16 | **0.0096** | 0.0689 | 9.4E-16 | **0.4264** | 0.9978 | 2.0E-16 | **0.3623** | 0.9177 | 2.0E-16 | **0.5856** | 0.9994 | 2.0E-16 |
| BBS11024 | 0.0244 | 0.2655 | 3.1E-02 | 0.0069 | 0.2986 | 3.1E-02 | 0.4017 | 0.8322 | 3.1E-02 | 0.3226 | 0.7386 | 3.1E-02 | 0.5443 | 0.9345 | 3.1E-02 |
| BBS11026 | **0.0873** | 0.2011 | 1.9E-04 | **0.0298** | 0.1951 | 3.4E-04 | **0.6577** | 0.7783 | 5.4E-04 | 0.6351 | 0.6894 | 1.6E-01 | **0.7841** | 0.8584 | 4.0E-03 |
| BBS11034 | **0.0196** | 0.0323 | 1.1E-03 | **0.0061** | 0.0071 | 1.1E-04 | **0.3908** | 0.6882 | 1.0E-07 | **0.3499** | 0.6283 | 7.5E-08 | **0.5500** | 0.7772 | 1.4E-07 |
| BBS11037 | **0.0229** | 0.2192 | 3.9E-03 | **0.0043** | 0.2037 | 5.9E-03 | **0.4064** | 0.6920 | 3.9E-03 | **0.3575** | 0.6180 | 3.9E-03 | **0.5548** | 0.7874 | 3.9E-03 |
| BBS11038 | **0.0167** | 0.0267 | 1.1E-05 | **0.0032** | 0.0062 | 7.2E-05 | **0.2270** | 0.2953 | 4.0E-06 | **0.1905** | 0.2583 | 7.0E-05 | **0.3564** | 0.4387 | 9.9E-05 |
| Domains won | **6** | 0 | | **6** | 0 | | **6** | 0 | | **5** | 0 | | **6** | 0 | |
| Tests won | | | | | | | | | | | | | | | |
| | Ankh | DEDAL | Equal | Ankh | DEDAL | Equal | Ankh | DEDAL | Equal | Ankh | DEDAL | Equal | Ankh | DEDAL | Equal |
| BBS11009 | 4 | 2 | 0 | 4 | 2 | 0 | 5 | 1 | 0 | 5 | 1 | 0 | 5 | 1 | 0 |
| BBS11010 | 6 | 0 | 0 | 6 | 0 | 0 | 5 | 1 | 0 | 5 | 1 | 0 | 5 | 1 | 0 |
| BBS11016 | **23** | 5 | 0 | **23** | 5 | 0 | **26** | 1 | 1 | **26** | 2 | 0 | **26** | 2 | 0 |
| BBS11018 | **88** | 3 | 0 | **84** | 7 | 0 | **88** | 3 | 0 | **88** | 3 | 0 | **88** | 3 | 0 |
| BBS11024 | 6 | 0 | 0 | 6 | 0 | 0 | 6 | 0 | 0 | 6 | 0 | 0 | 6 | 0 | 0 |
| BBS11026 | **19** | 1 | 1 | **17** | 3 | 1 | **15** | 3 | 3 | 13 | 7 | 1 | **14** | 6 | 1 |
| BBS11034 | **21** | 7 | 0 | **22** | 6 | 0 | **27** | 1 | 0 | **27** | 1 | 0 | **27** | 1 | 0 |
| BBS11037 | **9** | 1 | 0 | **9** | 1 | 0 | **9** | 1 | 0 | **9** | 1 | 0 | **9** | 1 | 0 |
| BBS11038 | **22** | 6 | 0 | **23** | 5 | 0 | **23** | 5 | 0 | **22** | 4 | 2 | **21** | 6 | 1 |
| Total | **182** | 23 | 1 | **178** | 27 | 1 | **188** | 14 | 4 | **172** | 11 | 2 | **185** | 19 | 2 |
| Tests won (%) | | | | | | | | | | | | | | | |
| BBS11009 | 66.67 | 33.33 | 0.00 | 66.67 | 33.33 | 0.00 | 83.33 | 16.67 | 0.00 | 83.33 | 16.67 | 0.00 | 83.33 | 16.67 | 0.00 |
| BBS11010 | 100.00 | 0.00 | 0.00 | 100.00 | 0.00 | 0.00 | 83.33 | 16.67 | 0.00 | 83.33 | 16.67 | 0.00 | 83.33 | 16.67 | 0.00 |
| BBS11016 | **82.14** | 17.86 | 0.00 | **82.14** | 17.86 | 0.00 | **92.86** | 3.57 | 3.57 | **92.86** | 7.14 | 0.00 | **92.86** | 7.14 | 0.00 |
| BBS11018 | **96.70** | 3.30 | 0.00 | **92.31** | 7.69 | 0.00 | **96.70** | 3.30 | 0.00 | **96.70** | 3.30 | 0.00 | **96.70** | 3.30 | 0.00 |
| BBS11024 | 100.00 | 0.00 | 0.00 | 100.00 | 0.00 | 0.00 | 100.00 | 0.00 | 0.00 | 100.00 | 0.00 | 0.00 | 100.00 | 0.00 | 0.00 |
| BBS11026 | **90.48** | 4.76 | 4.76 | **80.95** | 14.29 | 4.76 | **71.43** | 14.29 | 14.29 | 61.90 | 33.33 | 4.76 | **66.67** | 28.57 | 4.76 |
| BBS11034 | **75.00** | 25.00 | 0.00 | **78.57** | 21.43 | 0.00 | **96.43** | 3.57 | 0.00 | **96.43** | 3.57 | 0.00 | **96.43** | 3.57 | 0.00 |
| BBS11037 | **90.00** | 10.00 | 0.00 | **90.00** | 10.00 | 0.00 | **90.00** | 10.00 | 0.00 | **90.00** | 10.00 | 0.00 | **90.00** | 10.00 | 0.00 |
| BBS11038 | **78.57** | 21.43 | 0.00 | **82.14** | 17.86 | 0.00 | **82.14** | 17.86 | 0.00 | **78.57** | 14.29 | 7.14 | **75.00** | 21.43 | 3.57 |
| Average | **85.48** | 13.72 | 0.79 | **84.35** | 14.85 | 0.79 | **88.26** | 8.76 | 2.98 | **90.91** | 7.66 | 1.43 | **86.28** | 12.34 | 1.39 |

#### 7.3 BALiBASE RV12

Supplementary Table 45: Ankh-score vs DEDAL on BALiBASE RV12 dataset.

| Domain | $d_{cc}$ | | | $d_d$ | | | $d_{pos}$ | | | $d_{seq}$ | | | $d_{ssp}$ | | |
|---|---|---|---|---|---|---|---|---|---|---|---|---|---|---|---|
| Average distances | | | | | | | | | | | | | | | |
| | Ankh | DEDAL | P-value | Ankh | DEDAL | P-value | Ankh | DEDAL | P-value | Ankh | DEDAL | P-value | Ankh | DEDAL | P-value |
| BBS11009 | **0.0011** | 0.0259 | 1.1E-13 | **0.0003** | 0.0047 | 1.0E-11 | **0.0674** | 0.3656 | 3.6E-12 | **0.0635** | 0.3604 | 4.0E-12 | **0.1263** | 0.4203 | 3.2E-12 |
| BBS11026 | **0.0068** | 0.2249 | 1.6E-12 | **0.0031** | 0.2170 | 1.6E-12 | **0.3188** | 1.0000 | 1.6E-12 | **0.3062** | 0.9684 | 1.6E-12 | **0.4780** | 1.0000 | 1.6E-12 |
| BBS11034 | **0.0018** | 0.3030 | 6.1E-05 | **0.0002** | 0.2974 | 6.1E-05 | **0.0721** | 1.0000 | 6.1E-05 | **0.0701** | 0.9907 | 6.1E-05 | **0.1412** | 1.0000 | 6.1E-05 |
| BBS11037 | **0.0129** | 0.0630 | 9.5E-07 | **0.0032** | 0.0250 | 9.5E-07 | **0.2973** | 0.6950 | 8.9E-05 | **0.2360** | 0.6599 | 9.5E-07 | **0.4392** | 0.7613 | 9.5E-07 |
| BBS11038 | **0.0038** | 0.0607 | 1.7E-14 | **0.0022** | 0.0391 | 1.8E-14 | **0.1880** | 1.0000 | 1.7E-14 | **0.1726** | 0.9782 | 1.7E-14 | **0.3136** | 1.0000 | 1.7E-14 |
| Domains won | **5** | 0 | | **5** | 0 | | **5** | 0 | | **5** | 0 | | **5** | 0 | |
| Tests won | | | | | | | | | | | | | | | |
| | Ankh | DEDAL | Equal | Ankh | DEDAL | Equal | Ankh | DEDAL | Equal | Ankh | DEDAL | Equal | Ankh | DEDAL | Equal |
| BBS11009 | **72** | 6 | 0 | **69** | 9 | 0 | **71** | 6 | 1 | **71** | 6 | 1 | **71** | 6 | 1 |
| BBS11026 | **66** | 0 | 0 | **66** | 0 | 0 | **66** | 0 | 0 | **66** | 0 | 0 | **66** | 0 | 0 |
| BBS11034 | **15** | 0 | 0 | **15** | 0 | 0 | **15** | 0 | 0 | **15** | 0 | 0 | **15** | 0 | 0 |
| BBS11037 | **21** | 0 | 0 | **21** | 0 | 0 | **20** | 0 | 1 | **21** | 0 | 0 | **21** | 0 | 0 |
| BBS11038 | **78** | 0 | 0 | **77** | 1 | 0 | **78** | 0 | 0 | **78** | 0 | 0 | **78** | 0 | 0 |
| Total | **252** | 6 | 0 | **248** | 10 | 0 | **250** | 6 | 2 | **251** | 6 | 1 | **251** | 6 | 1 |
| Tests won (%) | | | | | | | | | | | | | | | |
| BBS11009 | **92.31** | 7.69 | 0.00 | **88.46** | 11.54 | 0.00 | **91.03** | 7.69 | 1.28 | **91.03** | 7.69 | 1.28 | **91.03** | 7.69 | 1.28 |
| BBS11026 | **100.00** | 0.00 | 0.00 | **100.00** | 0.00 | 0.00 | **100.00** | 0.00 | 0.00 | **100.00** | 0.00 | 0.00 | **100.00** | 0.00 | 0.00 |
| BBS11034 | **100.00** | 0.00 | 0.00 | **100.00** | 0.00 | 0.00 | **100.00** | 0.00 | 0.00 | **100.00** | 0.00 | 0.00 | **100.00** | 0.00 | 0.00 |
| BBS11037 | **100.00** | 0.00 | 0.00 | **100.00** | 0.00 | 0.00 | **95.24** | 0.00 | 4.76 | **100.00** | 0.00 | 0.00 | **100.00** | 0.00 | 0.00 |
| BBS11038 | **100.00** | 0.00 | 0.00 | **98.72** | 1.28 | 0.00 | **100.00** | 0.00 | 0.00 | **100.00** | 0.00 | 0.00 | **100.00** | 0.00 | 0.00 |
| Average | **98.46** | 1.54 | 0.00 | **97.44** | 2.56 | 0.00 | **97.25** | 1.54 | 1.21 | **98.21** | 1.54 | 0.26 | **98.21** | 1.54 | 0.26 |

#### 7.4 BALiBASE RV30

Supplementary Table 46: Ankh-score vs DEDAL on BALiBASE RV30 dataset.

| Domain | $d_{cc}$ | | | $d_d$ | | | $d_{pos}$ | | | $d_{seq}$ | | | $d_{ssp}$ | | |
|---|---|---|---|---|---|---|---|---|---|---|---|---|---|---|---|
| Average distances | | | | | | | | | | | | | | | |
| | Ankh | DEDAL | P-value | Ankh | DEDAL | P-value | Ankh | DEDAL | P-value | Ankh | DEDAL | P-value | Ankh | DEDAL | P-value |
| BBS30001 | **0.0157** | 0.0235 | 1.6E-78 | **0.0010** | 0.0019 | 1.3E-93 | **0.1513** | 0.3943 | 7.7E-92 | **0.1453** | 0.3849 | 2.8E-93 | **0.2360** | 0.4468 | 3.1E-94 |
| BBS30004 | **0.0019** | 0.0779 | 1.4E-154 | **0.0005** | 0.0656 | 2.3E-151 | **0.1171** | 0.5837 | 1.0E-157 | **0.1042** | 0.5650 | 1.4E-157 | **0.1935** | 0.6003 | 5.7E-159 |
| BBS30005 | **0.0078** | 0.0265 | 3.8E-119 | **0.0005** | 0.0024 | 3.7E-128 | **0.0817** | 0.3873 | 1.2E-134 | **0.0605** | 0.3682 | 1.9E-137 | **0.1422** | 0.4492 | 9.3E-138 |
| BBS30006 | **0.0566** | 0.2577 | 1.3E-22 | **0.0224** | 0.2490 | 5.3E-22 | **0.3454** | 0.6154 | 2.7E-22 | **0.3159** | 0.5353 | 1.6E-22 | **0.4445** | 0.6603 | 1.3E-22 |
| BBS30008 | **0.0200** | 0.1649 | 8.4E-90 | **0.0086** | 0.1469 | 5.9E-82 | **0.3952** | 0.5729 | 2.4E-75 | **0.3591** | 0.4785 | 2.4E-41 | **0.5062** | 0.6317 | 4.1E-75 |
| BBS30009 | 0.0694 | **0.0511** | 4.6E-30 | **0.0074** | 0.0175 | 8.2E-05 | 0.4691 | **0.3788** | 4.4E-32 | 0.4563 | **0.3650** | 4.0E-38 | 0.5083 | **0.4553** | 1.9E-26 |
| BBS30010 | **0.0013** | 0.0981 | 3.3E-165 | **0.0003** | 0.0667 | 3.4E-165 | **0.0845** | 1.0000 | 3.3E-165 | **0.0731** | 0.9937 | 3.3E-165 | **0.1404** | 1.0000 | 3.3E-165 |
| BBS30013 | **0.0135** | 0.1675 | 5.1E-165 | **0.0046** | 0.1438 | 5.8E-165 | **0.3144** | 1.0000 | 5.1E-165 | **0.2862** | 0.9495 | 5.2E-165 | **0.4238** | 1.0000 | 5.2E-165 |
| BBS30016 | 0.1917 | **0.0546** | 2.5E-10 | 0.1741 | **0.0182** | 4.3E-08 | 0.4616 | **0.3952** | 2.7E-06 | 0.4311 | **0.3695** | 1.4E-06 | 0.5044 | **0.4812** | 3.4E-05 |
| BBS30030 | **0.0043** | 0.0682 | 9.9E-146 | **0.0043** | 0.0433 | 2.5E-113 | **0.1262** | 0.6078 | 1.0E-143 | **0.1113** | 0.5820 | 2.8E-143 | **0.2148** | 0.6544 | 2.1E-145 |
| Domains won | **8** | 2 | | **9** | 1 | | **8** | 2 | | **8** | 2 | | **8** | 2 | |
| Tests won | | | | | | | | | | | | | | | |
| | Ankh | DEDAL | Equal | Ankh | DEDAL | Equal | Ankh | DEDAL | Equal | Ankh | DEDAL | Equal | Ankh | DEDAL | Equal |
| BBS30001 | **658** | 171 | 171 | **682** | 124 | 194 | **674** | 134 | 192 | **680** | 137 | 183 | **682** | 136 | 182 |
| BBS30004 | **930** | 52 | 18 | **945** | 37 | 18 | **950** | 30 | 20 | **951** | 26 | 23 | **957** | 25 | 18 |
| BBS30005 | **801** | 102 | 97 | **814** | 82 | 104 | **823** | 69 | 108 | **829** | 62 | 109 | **829** | 63 | 108 |
| BBS30006 | **135** | 11 | 7 | **135** | 10 | 8 | **135** | 6 | 12 | **137** | 6 | 10 | **137** | 7 | 9 |
| BBS30008 | **545** | 74 | 11 | **520** | 99 | 11 | **501** | 101 | 28 | **444** | 166 | 20 | **503** | 110 | 17 |
| BBS30009 | 193 | **343** | 167 | **294** | 242 | 167 | 178 | **317** | 208 | 144 | **352** | 207 | 177 | **319** | 207 |
| BBS30010 | **999** | 1 | 0 | **998** | 2 | 0 | **999** | 1 | 0 | **999** | 1 | 0 | **999** | 1 | 0 |
| BBS30013 | **997** | 2 | 1 | **993** | 6 | 1 | **996** | 3 | 1 | **995** | 4 | 1 | **995** | 4 | 1 |
| BBS30016 | 189 | **283** | 194 | 230 | **242** | 194 | 192 | **254** | 220 | 197 | **257** | 212 | 193 | **254** | 219 |
| BBS30030 | **896** | 71 | 33 | **791** | 172 | 37 | **877** | 79 | 44 | **871** | 79 | 50 | **878** | 77 | 45 |
| Total | **6343** | 1110 | 699 | **6402** | 1016 | 734 | **6325** | 994 | 833 | **6247** | 1090 | 815 | **6350** | 996 | 806 |
| Tests won (%) | | | | | | | | | | | | | | | |
| BBS30001 | **65.80** | 17.10 | 17.10 | **68.20** | 12.40 | 19.40 | **67.40** | 13.40 | 19.20 | **68.00** | 13.70 | 18.30 | **68.20** | 13.60 | 18.20 |
| BBS30004 | **93.00** | 5.20 | 1.80 | **94.50** | 3.70 | 1.80 | **95.00** | 3.00 | 2.00 | **95.10** | 2.60 | 2.30 | **95.70** | 2.50 | 1.80 |
| BBS30005 | **80.10** | 10.20 | 9.70 | **81.40** | 8.20 | 10.40 | **82.30** | 6.90 | 10.80 | **82.90** | 6.20 | 10.90 | **82.90** | 6.30 | 10.80 |
| BBS30006 | **88.24** | 7.19 | 4.58 | **88.24** | 6.54 | 5.23 | **88.24** | 3.92 | 7.84 | **89.54** | 3.92 | 6.54 | **89.54** | 4.58 | 5.88 |
| BBS30008 | **86.51** | 11.75 | 1.75 | **82.54** | 15.71 | 1.75 | **79.52** | 16.03 | 4.44 | **70.48** | 26.35 | 3.17 | **79.84** | 17.46 | 2.70 |
| BBS30009 | 27.45 | **48.79** | 23.76 | **41.82** | 34.42 | 23.76 | 25.32 | **45.09** | 29.59 | 20.48 | **50.07** | 29.45 | 25.18 | **45.38** | 29.45 |
| BBS30010 | **99.90** | 0.10 | 0.00 | **99.80** | 0.20 | 0.00 | **99.90** | 0.10 | 0.00 | **99.90** | 0.10 | 0.00 | **99.90** | 0.10 | 0.00 |
| BBS30013 | **99.70** | 0.20 | 0.10 | **99.30** | 0.60 | 0.10 | **99.60** | 0.30 | 0.10 | **99.50** | 0.40 | 0.10 | **99.50** | 0.40 | 0.10 |
| BBS30016 | 28.38 | **42.49** | 29.13 | 34.53 | **36.34** | 29.13 | 28.83 | **38.14** | 33.03 | 29.58 | **38.59** | 31.83 | 28.98 | **38.14** | 32.88 |
| BBS30030 | **89.60** | 7.10 | 3.30 | **79.10** | 17.20 | 3.70 | **87.70** | 7.90 | 4.40 | **87.10** | 7.90 | 5.00 | **87.80** | 7.70 | 4.50 |
| Average | **75.87** | 15.01 | 9.12 | **76.94** | 13.53 | 9.53 | **75.38** | 13.48 | 11.14 | **74.26** | 14.98 | 10.76 | **75.75** | 13.62 | 10.63 |

### 8 Comparison with BLOSUM on common overlap

Supplementary Table 47: Ankh-score vs BLOSUM45 on the common overlap, dataset CDD natural.

| Domain | $d_{cc}$ | | | $d_{d}$ | | | $d_{pos}$ | | | $d_{seq}$ | | | $d_{ssp}$ | | |
|---|---|---|---|---|---|---|---|---|---|---|---|---|---|---|---|
| Average distances | | | | | | | | | | | | | | | |
| | Ankh | BLOSUM45 | P-value | Ankh | BLOSUM45 | P-value | Ankh | BLOSUM45 | P-value | Ankh | BLOSUM45 | P-value | Ankh | BLOSUM45 | P-value |
| cd00012 | **0.0390** | 0.1201 | 1.6E-190 | **0.0201** | 0.0903 | 2.3E-177 | **0.5132** | 0.8704 | 3.2E-216 | **0.4645** | 0.8285 | 7.0E-221 | **0.8004** | 0.9479 | 2.3E-199 |
| cd00083 | **0.0024** | 0.0115 | 6.7E-101 | **0.0006** | 0.0031 | 5.9E-99 | **0.0422** | 0.1389 | 3.3E-100 | **0.0311** | 0.1276 | 2.7E-100 | **0.0907** | 0.2555 | 2.7E-100 |
| cd00173 | **0.0211** | 0.0496 | 1.3E-18 | **0.0053** | 0.0140 | 1.2E-17 | **0.2366** | 0.4801 | 3.7E-21 | **0.2195** | 0.4591 | 5.0E-21 | **0.4176** | 0.6219 | 1.3E-20 |
| cd00637 | **0.0045** | 0.0180 | 0.0E+00 | **0.0010** | 0.0054 | 0.0E+00 | **0.1288** | 0.3929 | 0.0E+00 | **0.1071** | 0.3789 | 0.0E+00 | **0.2200** | 0.5541 | 0.0E+00 |
| cd01040 | **0.0039** | 0.0269 | 2.8E-273 | **0.0007** | 0.0072 | 3.0E-281 | **0.0944** | 0.3429 | 1.4E-283 | **0.0760** | 0.3389 | 2.3E-285 | **0.1850** | 0.4897 | 2.7E-285 |
| cd01068 | **0.0028** | 0.0303 | 2.0E-22 | **0.0007** | 0.0048 | 1.3E-22 | **0.0886** | 0.2954 | 1.4E-24 | **0.0867** | 0.2916 | 1.6E-24 | **0.1757** | 0.4366 | 9.5E-25 |
| cd10140 | **0.0493** | 0.0698 | 1.8E-64 | **0.0099** | 0.0184 | 2.4E-73 | **0.6469** | 0.8191 | 4.3E-51 | **0.6293** | 0.8018 | 3.6E-52 | **0.8227** | 0.9021 | 3.9E-32 |
| cd21116 | **0.0103** | 0.0449 | 4.9E-51 | **0.0035** | 0.0167 | 5.3E-52 | **0.2566** | 0.5859 | 2.4E-54 | **0.2184** | 0.5659 | 1.6E-54 | **0.4147** | 0.7194 | 2.6E-54 |
| Domains won | **8** | 0 | | **8** | 0 | | **8** | 0 | | **8** | 0 | | **8** | 0 | |
| Tests won | | | | | | | | | | | | | | | |
| | Ankh | BLOSUM45 | Equal | Ankh | BLOSUM45 | Equal | Ankh | BLOSUM45 | Equal | Ankh | BLOSUM45 | Equal | Ankh | BLOSUM45 | Equal |
| cd00012 | **1276** | 104 | 0 | **1215** | 165 | 0 | **1337** | 39 | 4 | **1331** | 45 | 4 | **1204** | 41 | 135 |
| cd00083 | **610** | 34 | 175 | **601** | 27 | 191 | **605** | 24 | 190 | **605** | 23 | 191 | **605** | 23 | 191 |
| cd00173 | **117** | 18 | 1 | **119** | 16 | 1 | **118** | 16 | 2 | **121** | 14 | 1 | **119** | 15 | 2 |
| cd00637 | **2991** | 11 | 1 | **2966** | 36 | 1 | **2995** | 5 | 3 | **2998** | 3 | 2 | **2997** | 4 | 2 |
| cd01040 | **1707** | 91 | 38 | **1714** | 75 | 47 | **1713** | 62 | 61 | **1722** | 65 | 49 | **1721** | 68 | 47 |
| cd01068 | **136** | 6 | 8 | **135** | 7 | 8 | **139** | 3 | 8 | **139** | 3 | 8 | **139** | 3 | 8 |
| cd10140 | **468** | 94 | 8 | **477** | 86 | 7 | **432** | 110 | 28 | **440** | 112 | 18 | **344** | 118 | 108 |
| cd21116 | **312** | 11 | 0 | **307** | 16 | 0 | **317** | 5 | 1 | **317** | 6 | 0 | **314** | 9 | 0 |
| Total | **7617** | 369 | 231 | **7534** | 428 | 255 | **7656** | 264 | 297 | **7673** | 271 | 273 | **7443** | 281 | 493 |
| Tests won (%) | | | | | | | | | | | | | | | |
| cd00012 | **92.46** | 7.54 | 0.00 | **88.04** | 11.96 | 0.00 | **96.88** | 2.83 | 0.29 | **96.45** | 3.26 | 0.29 | **87.25** | 2.97 | 9.78 |
| cd00083 | **74.48** | 4.15 | 21.37 | **73.38** | 3.30 | 23.32 | **73.87** | 2.93 | 23.20 | **73.87** | 2.81 | 23.32 | **73.87** | 2.81 | 23.32 |
| cd00173 | **86.03** | 13.24 | 0.74 | **87.50** | 11.76 | 0.74 | **86.76** | 11.76 | 1.47 | **88.97** | 10.29 | 0.74 | **87.50** | 11.03 | 1.47 |
| cd00637 | **99.60** | 0.37 | 0.03 | **98.77** | 1.20 | 0.03 | **99.73** | 0.17 | 0.10 | **99.83** | 0.10 | 0.07 | **99.80** | 0.13 | 0.07 |
| cd01040 | **92.97** | 4.96 | 2.07 | **93.36** | 4.08 | 2.56 | **93.30** | 3.38 | 3.32 | **93.79** | 3.54 | 2.67 | **93.74** | 3.70 | 2.56 |
| cd01068 | **90.67** | 4.00 | 5.33 | **90.00** | 4.67 | 5.33 | **92.67** | 2.00 | 5.33 | **92.67** | 2.00 | 5.33 | **92.67** | 2.00 | 5.33 |
| cd10140 | **82.11** | 16.49 | 1.40 | **83.68** | 15.09 | 1.23 | **75.79** | 19.30 | 4.91 | **77.19** | 19.65 | 3.16 | **60.35** | 20.70 | 18.95 |
| cd21116 | **96.59** | 3.41 | 0.00 | **95.05** | 4.95 | 0.00 | **98.14** | 1.55 | 0.31 | **98.14** | 1.86 | 0.00 | **97.21** | 2.79 | 0.00 |
| Average | **89.36** | 6.77 | 3.87 | **88.72** | 7.13 | 4.15 | **89.64** | 5.49 | 4.87 | **90.11** | 5.44 | 4.45 | **86.55** | 5.77 | 7.69 |

### 9 Comparison against ProtT5+ESM2

Supplementary Table 48: ProtT5+ESM2-score vs ProtT5-score on CDD natural dataset.

| Domain | $d_{cc}$ | | | $d_d$ | | | $d_{pos}$ | | | $d_{seq}$ | | | $d_{ssp}$ | | |
|---|---|---|---|---|---|---|---|---|---|---|---|---|---|---|---|
| Average distances | | | | | | | | | | | | | | | |
| | ProtT5+ESM2 | ProtT5 | P-value | ProtT5+ESM2 | ProtT5 | P-value | ProtT5+ESM2 | ProtT5 | P-value | ProtT5+ESM2 | ProtT5 | P-value | ProtT5+ESM2 | ProtT5 | P-value |
| cd00012 | 0.0478 | 0.0441 | 9.1E-02 | 0.0266 | **0.0242** | 2.9E-14 | 0.6962 | 0.6956 | 9.2E-02 | 0.6211 | **0.6100** | 1.1E-05 | 0.8150 | 0.8139 | 2.7E-01 |
| cd00083 | **0.0043** | 0.0063 | 9.4E-62 | **0.0011** | 0.0020 | 1.2E-65 | **0.0643** | 0.0869 | 1.8E-60 | **0.0523** | 0.0765 | 7.7E-64 | **0.1178** | 0.1559 | 2.8E-62 |
| cd00173 | 0.0350 | **0.0346** | 8.2E-04 | 0.0083 | 0.0082 | 1.6E-02 | **0.4529** | 0.4589 | 8.0E-03 | **0.4259** | 0.4310 | 8.9E-03 | **0.5720** | 0.5767 | 8.5E-03 |
| cd00637 | **0.0040** | 0.0041 | 4.3E-12 | 0.0009 | **0.0009** | 5.6E-04 | **0.1267** | 0.1316 | 4.4E-43 | **0.1025** | 0.1083 | 2.7E-59 | **0.2151** | 0.2239 | 4.3E-65 |
| cd01040 | **0.0043** | 0.0052 | 1.9E-92 | **0.0008** | 0.0010 | 1.8E-66 | **0.1124** | 0.1208 | 1.5E-68 | **0.0893** | 0.1005 | 1.2E-95 | **0.1890** | 0.2036 | 1.1E-95 |
| cd01068 | 0.0030 | 0.0028 | 8.3E-01 | 0.0008 | 0.0007 | 7.8E-01 | 0.1119 | 0.1093 | 2.4E-01 | 0.1091 | 0.1068 | 3.0E-01 | 0.1819 | 0.1778 | 3.1E-01 |
| cd10140 | 0.0626 | 0.0611 | 6.5E-01 | 0.0136 | 0.0126 | 3.0E-02 | 0.7317 | 0.7372 | 7.9E-02 | 0.6805 | 0.6907 | 8.8E-01 | 0.8376 | 0.8435 | 1.0E-01 |
| cd21116 | 0.0100 | 0.0101 | 3.1E-02 | 0.0037 | 0.0037 | 2.7E-01 | 0.3215 | 0.3255 | 5.1E-02 | 0.2838 | 0.2846 | 4.2E-02 | 0.4472 | 0.4562 | 2.8E-02 |
| Domains won | **3** | 1 | | **2** | **2** | | **4** | 0 | | **4** | 1 | | **4** | 0 | |
| Tests won | | | | | | | | | | | | | | | |
| | ProtT5+ESM2 | ProtT5 | Equal | ProtT5+ESM2 | ProtT5 | Equal | ProtT5+ESM2 | ProtT5 | Equal | ProtT5+ESM2 | ProtT5 | Equal | ProtT5+ESM2 | ProtT5 | Equal |
| cd00012 | 898 | 852 | 20 | 786 | **968** | 16 | 779 | 749 | 242 | 799 | **861** | 110 | 841 | 821 | 108 |
| cd00083 | **587** | 72 | 422 | **586** | 63 | 432 | **584** | 64 | 433 | **584** | 65 | 432 | **584** | 65 | 432 |
| cd00173 | **151** | 86 | 88 | 147 | 88 | 90 | **138** | 79 | 108 | **141** | 78 | 106 | **145** | 75 | 105 |
| cd00637 | **1681** | 1252 | 70 | **1511** | 1423 | 69 | **1717** | 997 | 289 | **1769** | 998 | 236 | **1791** | 1001 | 211 |
| cd01040 | **1194** | 373 | 324 | **1121** | 423 | 347 | **1059** | 360 | 472 | **1136** | 340 | 415 | **1136** | 346 | 409 |
| cd01068 | 48 | 43 | 80 | 46 | 45 | 80 | 43 | 45 | 83 | 45 | 45 | 81 | 45 | 45 | 81 |
| cd10140 | 280 | 280 | 70 | 277 | 287 | 66 | 169 | 147 | 314 | 215 | 236 | 179 | 198 | 200 | 232 |
| cd21116 | 177 | 140 | 8 | 171 | 145 | 9 | 156 | 128 | 41 | 160 | 134 | 31 | 158 | 137 | 30 |
| Total | **3613** | 1783 | 904 | **4004** | 2877 | 864 | **3498** | 1500 | 1302 | **4429** | 2342 | 1299 | **3656** | 1487 | 1157 |
| Tests won (%) | | | | | | | | | | | | | | | |
| cd00012 | 50.73 | 48.14 | 1.13 | 44.41 | **54.69** | 0.90 | 44.01 | 42.32 | 13.67 | 45.14 | **48.64** | 6.21 | 47.51 | 46.38 | 6.10 |
| cd00083 | **54.30** | 6.66 | 39.04 | **54.21** | 5.83 | 39.96 | **54.02** | 5.92 | 40.06 | **54.02** | 6.01 | 39.96 | **54.02** | 6.01 | 39.96 |
| cd00173 | **46.46** | 26.46 | 27.08 | 45.23 | 27.08 | 27.69 | **42.46** | 24.31 | 33.23 | **43.38** | 24.00 | 32.62 | **44.62** | 23.08 | 32.31 |
| cd00637 | **55.98** | 41.69 | 2.33 | **50.32** | 47.39 | 2.30 | **57.18** | 33.20 | 9.62 | **58.91** | 33.23 | 7.86 | **59.64** | 33.33 | 7.03 |
| cd01040 | **63.14** | 19.73 | 17.13 | **59.28** | 22.37 | 18.35 | **56.00** | 19.04 | 24.96 | **60.07** | 17.98 | 21.95 | **60.07** | 18.30 | 21.63 |
| cd01068 | 28.07 | 25.15 | 46.78 | 26.90 | 26.32 | 46.78 | 25.15 | 26.32 | 48.54 | 26.32 | 26.32 | 47.37 | 26.32 | 26.32 | 47.37 |
| cd10140 | 44.44 | 44.44 | 11.11 | 43.97 | 45.56 | 10.48 | 26.83 | 23.33 | 49.84 | 34.13 | 37.46 | 28.41 | 31.43 | 31.75 | 36.83 |
| cd21116 | 54.46 | 43.08 | 2.46 | 52.62 | 44.62 | 2.77 | 48.00 | 39.38 | 12.62 | 49.23 | 41.23 | 9.54 | 48.62 | 42.15 | 9.23 |
| Average | **54.97** | 23.63 | 21.39 | **52.05** | 32.57 | 15.38 | **52.42** | 20.62 | 26.97 | **52.31** | 25.97 | 21.72 | **54.59** | 20.18 | 25.23 |

Supplementary Table 49: Ankh-score vs ProtT5+ESM2-score on CDD natural dataset.

| Domain | $d_{cc}$ | | | $d_d$ | | | $d_{pos}$ | | | $d_{seq}$ | | | $d_{ssp}$ | | |
|---|---|---|---|---|---|---|---|---|---|---|---|---|---|---|---|
| Average distances | | | | | | | | | | | | | | | |
| | Ankh | ProtT5+ESM2 | P-value | Ankh | ProtT5+ESM2 | P-value | Ankh | ProtT5+ESM2 | P-value | Ankh | ProtT5+ESM2 | P-value | Ankh | ProtT5+ESM2 | P-value |
| cd00012 | **0.0446** | 0.0478 | 6.7E-13 | **0.0235** | 0.0266 | 1.7E-35 | **0.6738** | 0.6962 | 4.7E-37 | **0.5926** | 0.6211 | 2.4E-39 | **0.7964** | 0.8150 | 4.1E-36 |
| cd00083 | **0.0026** | 0.0043 | 1.1E-22 | **0.0006** | 0.0011 | 9.1E-20 | **0.0430** | 0.0643 | 4.2E-23 | **0.0333** | 0.0523 | 1.3E-26 | **0.0885** | 0.1178 | 1.8E-25 |
| cd00173 | 0.0335 | 0.0350 | 4.9E-01 | 0.0081 | 0.0083 | 4.8E-02 | 0.4884 | 0.4529 | 5.2E-01 | 0.4576 | 0.4259 | 5.1E-01 | 0.5958 | 0.5720 | 5.0E-01 |
| cd00637 | 0.0044 | **0.0040** | 2.8E-62 | 0.0010 | **0.0009** | 2.6E-38 | 0.1315 | **0.1267** | 1.0E-06 | 0.1055 | **0.1025** | 1.9E-10 | 0.2211 | **0.2151** | 1.2E-09 |
| cd01040 | **0.0037** | 0.0043 | 3.2E-76 | **0.0007** | 0.0008 | 7.4E-97 | **0.1006** | 0.1124 | 1.5E-75 | **0.0795** | 0.0893 | 1.3E-78 | **0.1741** | 0.1890 | 3.9E-75 |
| cd01068 | **0.0027** | 0.0030 | 5.6E-03 | 0.0006 | 0.0008 | 1.5E-01 | **0.1046** | 0.1119 | 1.1E-05 | **0.1028** | 0.1091 | 2.6E-04 | **0.1720** | 0.1819 | 1.6E-04 |
| cd10140 | **0.0596** | 0.0626 | 2.8E-13 | **0.0120** | 0.0136 | 1.2E-13 | 0.7260 | 0.7317 | 7.5E-02 | **0.6739** | 0.6805 | 3.9E-04 | 0.8359 | 0.8376 | 2.0E-01 |
| cd21116 | 0.0084 | 0.0100 | 5.2E-02 | 0.0029 | 0.0037 | 3.0E-02 | **0.2944** | 0.3215 | 3.7E-07 | **0.2594** | 0.2838 | 1.0E-04 | **0.4129** | 0.4472 | 3.1E-06 |
| Domains won | **5** | 1 | | **4** | 1 | | **5** | 1 | | **6** | 1 | | **5** | 1 | |
| Tests won | | | | | | | | | | | | | | | |
| | Ankh | ProtT5+ESM2 | Equal | Ankh | ProtT5+ESM2 | Equal | Ankh | ProtT5+ESM2 | Equal | Ankh | ProtT5+ESM2 | Equal | Ankh | ProtT5+ESM2 | Equal |
| cd00012 | **978** | 786 | 6 | **1084** | 680 | 6 | **1020** | 617 | 133 | **1073** | 637 | 60 | **1064** | 650 | 56 |
| cd00083 | **533** | 191 | 357 | **512** | 165 | 404 | **508** | 163 | 410 | **510** | 162 | 409 | **510** | 165 | 406 |
| cd00173 | 141 | 117 | 67 | 147 | 109 | 69 | 130 | 105 | 90 | 131 | 109 | 85 | 129 | 112 | 84 |
| cd00637 | 1136 | **1832** | 35 | 1200 | **1771** | 32 | 1280 | **1504** | 219 | 1280 | **1542** | 181 | 1309 | **1557** | 137 |
| cd01040 | **1199** | 457 | 235 | **1184** | 437 | 270 | **1084** | 429 | 378 | **1112** | 432 | 347 | **1110** | 442 | 339 |
| cd01068 | **74** | 45 | 52 | 66 | 49 | 56 | **72** | 33 | 66 | **71** | 38 | 62 | **71** | 38 | 62 |
| cd10140 | **364** | 224 | 42 | **369** | 218 | 43 | 192 | 187 | 251 | **268** | 207 | 155 | 233 | 207 | 190 |
| cd21116 | 175 | 149 | 1 | 186 | 136 | 3 | **181** | 124 | 20 | **176** | 136 | 13 | **177** | 136 | 12 |
| Total | **4284** | 3535 | 727 | **4349** | 3271 | 755 | **4145** | 2870 | 1226 | **4490** | 3154 | 1227 | **4241** | 2988 | 1012 |
| Tests won (%) | | | | | | | | | | | | | | | |
| cd00012 | **55.25** | 44.41 | 0.34 | **61.24** | 38.42 | 0.34 | **57.63** | 34.86 | 7.51 | **60.62** | 35.99 | 3.39 | **60.11** | 36.72 | 3.16 |
| cd00083 | **49.31** | 17.67 | 33.02 | **47.36** | 15.26 | 37.37 | **46.99** | 15.08 | 37.93 | **47.18** | 14.99 | 37.84 | **47.18** | 15.26 | 37.56 |
| cd00173 | 43.38 | 36.00 | 20.62 | 45.23 | 33.54 | 21.23 | 40.00 | 32.31 | 27.69 | 40.31 | 33.54 | 26.15 | 39.69 | 34.46 | 25.85 |
| cd00637 | 37.83 | **61.01** | 1.17 | 39.96 | **58.97** | 1.07 | 42.62 | **50.08** | 7.29 | 42.62 | **51.35** | 6.03 | 43.59 | **51.85** | 4.56 |
| cd01040 | **63.41** | 24.17 | 12.43 | **62.61** | 23.11 | 14.28 | **57.32** | 22.69 | 19.99 | **58.80** | 22.85 | 18.35 | **58.70** | 23.37 | 17.93 |
| cd01068 | **43.27** | 26.32 | 30.41 | 38.60 | 28.65 | 32.75 | **42.11** | 19.30 | 38.60 | **41.52** | 22.22 | 36.26 | **41.52** | 22.22 | 36.26 |
| cd10140 | **57.78** | 35.56 | 6.67 | **58.57** | 34.60 | 6.83 | 30.48 | 29.68 | 39.84 | **42.54** | 32.86 | 24.60 | 36.98 | 32.86 | 30.16 |
| cd21116 | 53.85 | 45.85 | 0.31 | 57.23 | 41.85 | 0.92 | **55.69** | 38.15 | 6.15 | **54.15** | 41.85 | 4.00 | **54.46** | 41.85 | 3.69 |
| Average | **51.14** | 34.85 | 14.01 | **53.95** | 34.07 | 11.98 | **50.39** | 30.03 | 19.58 | **49.63** | 31.73 | 18.64 | **50.93** | 31.88 | 17.19 |
